# Microglial AhR Drives Sex-Biased Amyloid Pathology

**DOI:** 10.64898/2026.11.05.764518

**Authors:** Carmen Nieto-Vaquero, Alicia García-Culebras, Andrea Rubio-Ponce, David Chamoso-Sánchez, Manuel J. Gómez, F. Javier de Castro-Millán, Carlos Parra-Pérez, Sandra Vázquez-Reyes, Aurora Semerano, Isabel Bravo-Ferrer, Ana Moraga-Yébenes, Coral Barbas, Héctor Bueno, Estrella Morenas-Rodríguez, Fátima Sánchez-Cabo, Alberto Villarejo-Galende, Takashi Saito, Takaomi C. Saido, Pedro M. Fernández-Salguero, Ignacio Lizasoain, María Isabel Cuartero, María Ángeles Moro

**Author notes:** Correspondence to María I. Cuartero or María A. Moro. These authors contributed equally as corresponding authors (María Isabel Cuartero, María Ángeles Moro).

## Abstract

Microglia integrate local and systemic signals that can modify their response to amyloid-β (Aβ), but the molecular pathways connecting these signals to Alzheimer’s disease (AD) pathology remain incompletely understood. Here, we identify the aryl hydrocarbon receptor (AhR), a sensor of microbiota- and host-derived metabolites, as a microglial regulator of amyloid pathology. Sustained administration of the AhR agonist indoxyl-3-sulfate (I3S) exacerbated cortical Aβ deposition and plaque-associated neuritic damage in APP^NL-F^ mice. Conversely, constitutive AhR deficiency reduced amyloid burden, promoted plaque compaction, and preserved behavioural performance, whereas inducible deletion of AhR in microglia was sufficient to reduce plaque accumulation. Single-cell RNA sequencing revealed that AhR loss reprogrammed microglia toward phagolysosomal and Aβ-recognition states and enhanced microglial Aβ uptake, with markedly stronger effects in females. APP^NL-F^ mice also exhibited sex-dependent remodelling of the gut microbiota and tryptophan-derived metabolites. Circulating indole-3-aldehyde (IAld) robustly activated AhR and was associated with cortical amyloid pathology. AhR-activating bioactivity was increased in mouse serum, cortex and faecal extracts and was higher in serum from the COGNIBIO12 analytical subcohort than in independently recruited cognitively unimpaired H2H participants. Among amyloid-positive patients, higher serum AhR bioactivity was associated with lower CSF Aβ42 concentrations. Together, these findings identify microglial AhR as a metabolically responsive regulator linking systemic ligand availability, biological sex, and microglial control of amyloid pathology.

## Introduction

Microglia, the tissue-resident macrophages of the central nervous system, are key regulators of amyloid-β (Aβ) pathology in Alzheimer’s disease (AD). They recognize and engulf Aβ and accumulate around amyloid deposits, where they promote plaque compaction and limit the exposure of surrounding neurites to potentially toxic Aβ species^1^. Chronic exposure to the diseased environment, however, induces heterogeneous transcriptional and functional states that can alter these protective interactions with Aβ^2^. The importance of microglial function in AD is reinforced by human genetic studies, which have identified numerous AD risk genes enriched in microglia and implicated innate immune, lipid-handling and phagocytic pathways in disease susceptibility^3, 4, 5, 6^. Microglial responses to amyloid pathology are also influenced by biological sex; in APP knock-in mice, female microglia undergo a more rapid transition towards plaque-associated transcriptional states^7^. The molecular checkpoints that integrate environmental signals to shape these microglial responses, and whether their activity differs between sexes, remain poorly understood.

Signals from the gut microbiota (GM) are emerging as modulators of microglial function. Gut microbial dysbiosis has been described in patients with AD^8^, while experimental manipulation of the GM modifies Aβ deposition, plaque-associated microglial phenotypes, and microglial transcriptional programs in mouse models of amyloidosis^9, 10, 11^. These effects can be sex dependent: administration of *Bacteroides fragilis* increased Aβ plaque deposition selectively in female APP/PS1 mice and was associated with suppression of microglial pathways related to phagocytosis and protein degradation^12^. Among microbiota-derived signals, tryptophan metabolites are particularly relevant because intestinal bacteria convert dietary tryptophan into diverse indole derivatives, several of which enter the systemic circulation and activate the aryl hydrocarbon receptor (AhR)^13, 14^. Thus, AhR provides a potential mechanism through which the systemic metabolic environment can regulate microglial responses to tissue pathology.

AhR is a ligand-activated transcription factor that integrates signals from microbial and host-derived tryptophan metabolites, dietary compounds, and environmental ligands to regulate immune-cell function^15, 16, 17^. In the central nervous system (CNS), AhR controls glial transcriptional programs and mediates responses to microbiota-derived metabolites^18, 19^. Emerging evidence also links this pathway to AD, with seemingly divergent consequences. Activation of the IDO1–kynurenine–AhR axis has been implicated in Aβ-induced neuronal toxicity, Wnt/β-catenin dysregulation and tau phosphorylation. More recently, IDO1-derived kynurenine was shown to suppress astrocytic glycolysis in an AhR-dependent manner across amyloid and tau models^20, 21^. Conversely, Indole, indole-3-acetic acid (IAA) and indole-3-propionic acid attenuated neuroinflammation and amyloid-related pathology in APP/PS1 mice through AhR-associated mechanisms^22^, whereas microbiome-derived indole-3-lactic acid (ILA) reduced amyloid accumulation through AhR-dependent effects involving microglia and astrocytes in 5xFAD mice^23^. More recently, IAA was linked to reduced microglia-mediated synaptic loss through an AhR-CCR4 pathway^24^.

Together, these studies indicate that the consequences of AhR signalling depend on ligand and cellular context but leave unresolved whether AhR acts within microglia to directly control their response to Aβ, which microglial programs are regulated by AhR, and whether this regulation is influenced by biological sex.

Here, using sustained treatment with an AhR agonist, and constitutive global and inducible microglial genetic deletion of AhR in APP^NL-F^ mice, we investigated whether and how microglial AhR regulates plaque-associated immune responses and amyloid pathology. We find that AhR constrains Aβ-recognition and phagolysosomal programs in microglia and limits their engagement with amyloid deposits, with substantially stronger effects in females. These findings identify microglial AhR as a ligand-responsive checkpoint that links systemic metabolic signals to sex-dependent innate immune responses in amyloid pathology.

## Results

### Activation of AhR with indoxyl-3-sulfate increases cortical amyloid pathology and neuritic dystrophy in APP^NL-F^ mice

To evaluate the effect of AhR activation in AD, APP^NL-F^ knock-in mice were chronically treated with indoxyl-3-sulfate (I3S), a well-established AhR agonist (Figure 1A). APP^NL-F^ mice developed robust and regionally restricted amyloid pathology, with Aβ plaques predominantly confined to the cerebral cortex and virtually absent from the hippocampus, allowing a precise assessment of cortical amyloidosis at 12 months of age (Supplementary Figure 1)^25^. I3S treatment resulted in a significant increase in the number of both ThioS-positive fibrillar plaques and total Aβ-positive deposits (Figure 1B-D), together with a greater percentage of cortical area occupied by Aβ deposits, indicating an overall increase in amyloid burden (Supplementary Figure 2). Importantly, I3S treatment increased plaque-associated neuritic dystrophy, evidenced by a greater LAMP1-positive dystrophic neurite area surrounding amyloid deposits (Figure 1E). To determine whether these histological changes were accompanied by alterations in cortical Aβ levels, we measured Aβ in TBS-soluble and guanidine-HCl-extracted insoluble cortical fractions. Consistent with the histological findings, I3S significantly increased soluble Aβ42, with no significant difference in Aβ40 or the Aβ42/Aβ40 ratio (Figure 1F). Insoluble Aβ42 showed a trend toward increased levels, whereas Aβ40 and the Aβ42/Aβ40 ratio remained unchanged (Figure 1G). Together, these findings indicate that sustained AhR activation increases cortical Aβ42, with a more pronounced effect in the soluble fraction. We also assessed the behavioural consequences in I3S-treated APP^NL-F^ mice. Although no significant treatment differences were detected in novel object recognition or spontaneous alternation in the Y-maze (Supplementary Figure 2), I3S treatment markedly reduced locomotor and exploratory activity in the open-field test. (Figure 1H-L and Supplementary Figure 2D). Thus, AhR activation by I3S treatment increased cortical amyloid deposition and plaque-associated neuritic damage and reduced locomotor and exploratory activity, without detectable deficits in the memory measures assessed at this age.

**Figure 1.**
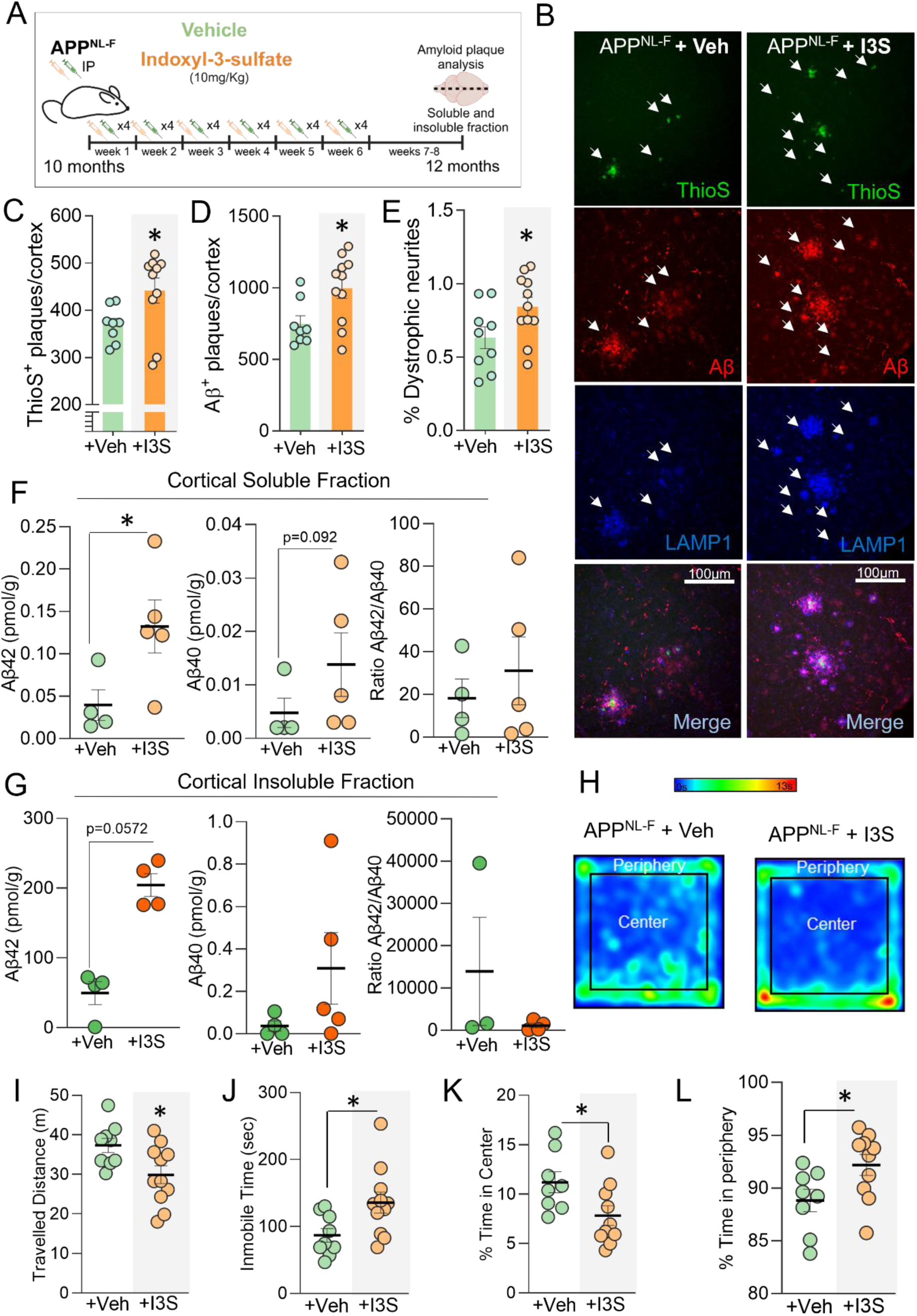
I3S treatment increases cortical amyloid pathology and reduces locomotor and exploratory activity in APP^NL-F^ mice. **(A)** Experimental design. APP^NL-F^ mice received intraperitoneal injections of vehicle or indoxyl-3-sulfate (I3S; 10 mg/kg) every other day for 6 weeks, beginning at 10 months of age. Amyloid pathology, cortical Aβ levels and behavior were assessed at 12 months of age. Both male and female mice were included in the experimental groups. **(B)** Representative cortical images showing Thioflavin S-positive fibrillar plaques (ThioS, green), Aβ immunoreactivity detected with the 6E10 antibody (red), LAMP1-positive dystrophic neurites (blue) and merged channels in vehicle-and I3S-treated APP^NL-F^ mice. Arrowheads indicate amyloid deposits and associated dystrophic neurites. Scale bar=100 μm. **(C)** Number of ThioS-positive plaques per cortex. Vehicle: n=8; I3S: n=9. Two-tailed exact Mann-Whitney U test, p=0.0343. **(D)** Number of Aβ-positive plaques per cortex. Vehicle: n=8; I3S: n=11. Two-tailed exact Mann-Whitney U test, p=0.0259. **(E)** Percentage of cortical area occupied by LAMP1-positive dystrophic neurites. Vehicle: n=9; I3S: n=11. Two-tailed exact Mann-Whitney U test, p=0.0326. **(F)** Cortical Aβ42 and Aβ40 concentrations and Aβ42/Aβ40 ratio measured by ELISA in the TBS-soluble fraction. Vehicle: n=4; I3S: n=5. Two-tailed exact Mann-Whitney U tests. Aβ42, p=0.0317; Aβ40, p=0.0920; Aβ42/Aβ40 ratio, not significant. **(G)** Cortical Aβ42 and Aβ40 concentrations and Aβ42/Aβ40 ratio measured in the guanidine-HCl-extracted insoluble fraction. Vehicle: n=4; I3S: n=5. Two-tailed exact Mann-Whitney U tests. Aβ42, p=0.0572; Aβ40 and Aβ42/Aβ40 ratio, not significant. **(H)** Representative occupancy heat maps obtained during the open-field test. **(I-L)** Total distance travelled **(I)**, immobility time **(J)**, percentage of time spent in the center **(K)** and percentage of time spent in the periphery **(L)** of the open-field arena. Vehicle: n=10; I3S: n=11. Two-tailed exact Mann-Whitney U tests; **(I)** p=0.0465; **(J)** p=0.0192; **(K)** p=0.0343; **(L)** p=0.0343. Data are presented as mean±SEM. Each dot represents one biologically independent mouse. Sex distribution across experimental groups was as follows: (C) vehicle, 5 males/3 females; I3S, 6 males/4 females; (D) vehicle, 5 males/3 females; I3S, 6 males/5 females; (E) vehicle, 5 males/4 females; I3S, 6 males/5 females; (F,G) vehicle, 2 males/2 females; I3S, 3 males/2 females; (I,J) vehicle, 5 males/4 females; I3S, 6 males/5 females; and (K,L) vehicle, 5 males/3 females; I3S, 5 males/5 females. Statistical significance was considered at p<0.05.

### Constitutive deletion of AhR reduces cortical amyloid pathology and improves behavioural performance in AD mice

We next tested whether constitutive global AhR loss affects disease progression. For this purpose, we generated APP^NL-F^/AhR^-/-^mice and compared them with APP^NL-F^/AhR^+/-^ and APP^NL-F^/AhR^+/+^ mice at 12 months of age (Figure 2A-B). Constitutive AhR loss markedly reduced the number of cortical ThioS-positive fibrillar plaques and total Aβ-positive deposits, together with cortical Aβ burden and LAMP1-positive dystrophic neurites (Figure 2A-F). Heterozygous animals generally displayed an intermediate phenotype, suggesting an AhR gene-dosage effect. Importantly, this protective effect was already detectable at 6 months, when APP^NL-F^/AhR^-/-^mice showed fewer cortical Aβ-positive plaques and a similar downward trend in ThioS-positive plaques (Supplementary Figure 3A). Biochemical profiling further revealed differences in cortical Aβ concentrations following AhR loss. In the TBS-soluble fraction, Aβ42 and the Aβ42/Aβ40 ratio was markedly reduced (Figure 2G), whereas soluble Aβ40 was increased (Supplementary Figure 3B). In the guanidine-HCl-extracted insoluble fraction, Aβ42 was significantly reduced and the Aβ42/Aβ40 ratio showed a downward trend (Figure 2H), while insoluble Aβ40 remained unchanged (Supplementary Figure 3B). Together, these findings associate AhR loss with lower cortical Aβ concentrations, limiting the accumulation of Aβ42.

**Figure 2.**
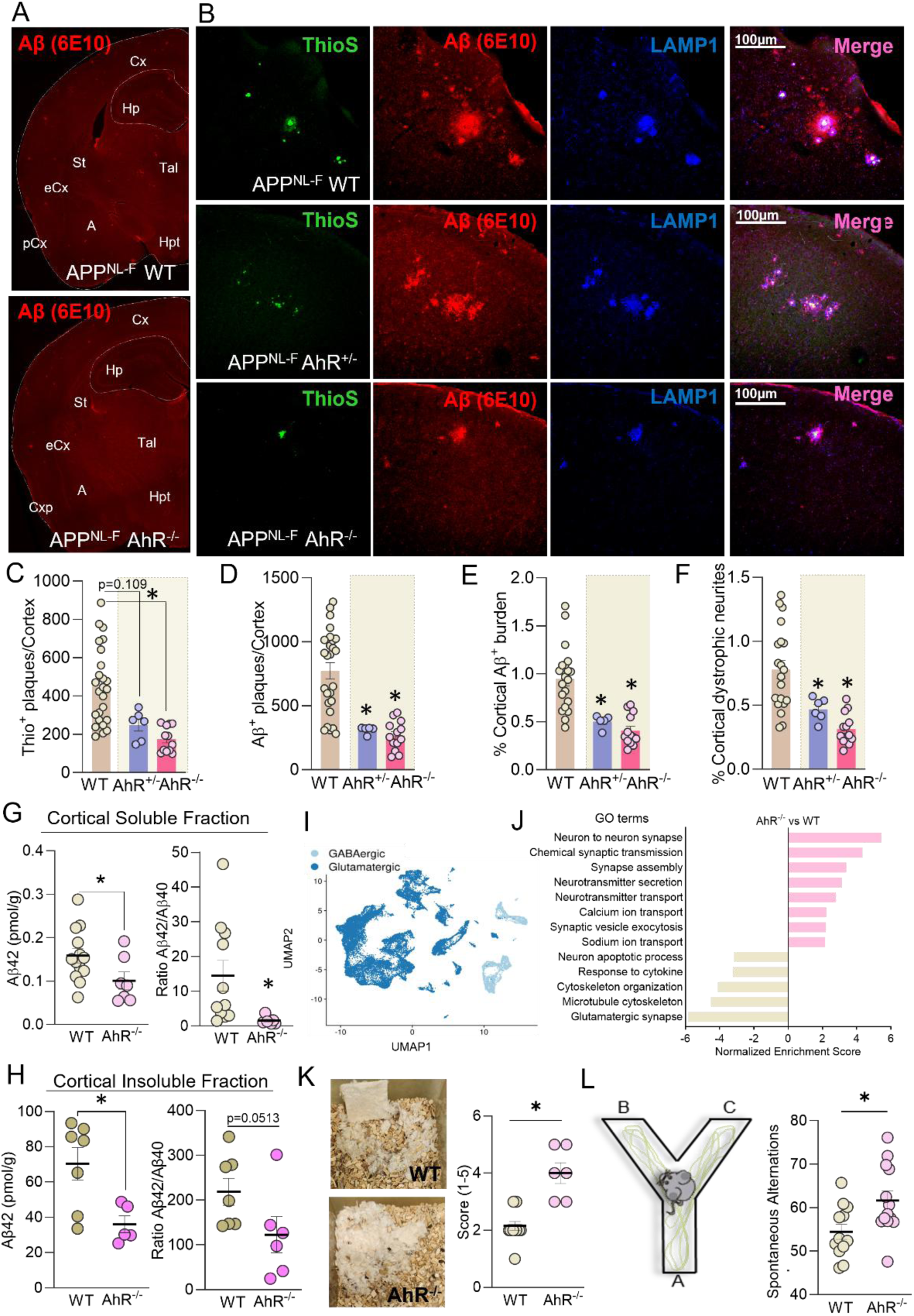
Constitutive loss of AhR attenuates cortical amyloid pathology, shifts Aβ composition and improves behavioural performance in APP^NL-F^ mice. **(A)** Representative coronal brain sections from 12-month-old APP^NL-F^/AhR^+/+^ and APP^NL-F^/AhR^-/-^mice showing Aβ immunoreactivity detected with the 6E10 antibody. **(B)** Representative cortical images showing Thioflavin S-positive fibrillar plaques (ThioS, green), Aβ immunoreactivity detected with the 6E10 antibody (red), LAMP1-positive dystrophic neurites (blue) and merged channels in APP^NL-F^/AhR^+/+^, APP^NL-F^/AhR^+/-^ and APP^NL-F^/AhR^-/-^mice. Scale bar = 100 μm. **(C)** Number of ThioS-positive plaques per cortex. APP^NL-F^/AhR^+/+^: n=25; APP^NL-F^/AhR^+/-^: n=6; APP^NL-F^/AhR^-/-^: n=12. Kruskal-Wallis test followed by Dunn’s multiple-comparisons test. Adjusted p=0.109 for APP^NL-F^/AhR^+/+^ versus APP^NL-F^/AhR^+/-^ and p<0.0001 for APP^NL-F^/AhR^+/+^ versus APP^NL-F^/AhR^-/-^. **(D)** Number of Aβ-positive plaques per cortex; n=26, n=5 and n=13, respectively. Kruskal-Wallis test followed by Dunn’s multiple-comparisons test. Adjusted p=0.0267 and p<0.0001, respectively. **(E)** Percentage of cortical area occupied by Aβ deposits; n=20, n=5 and n=13, respectively. Kruskal-Wallis test followed by Dunn’s multiple-comparisons test. Adjusted p=0.0207 and p<0.0001, respectively. **(F)** Percentage of cortical area occupied by LAMP1-positive dystrophic neurites; n=20, n=6 and n=13, respectively. Kruskal-Wallis test followed by Dunn’s multiple-comparisons test. Adjusted p=0.1573 and p<0.0001, respectively. **(G)** Cortical Aβ42 concentration and Aβ42/Aβ40 ratio measured by ELISA in the TBS-soluble fraction. For Aβ42, APP^NL-F^/AhR^+/+^: n=13 and APP^NL-F^/AhR^-/-^: n=7; for the Aβ42/Aβ40 ratio, n=11 and n=7, respectively. Two-tailed t test for Aβ42, p=0.0048; exact Mann-Whitney U test for the Aβ42/Aβ40 ratio, p=0.0012. **(H)** Cortical Aβ42 concentration and Aβ42/Aβ40 ratio measured in the guanidine-HCl-extracted insoluble fraction. For Aβ42, APP^NL-F^/AhR^+/+^: n=7 and APP^NL-F^/AhR^-/-^: n=5; and for the Aβ42/Aβ40 ratio, n=7 and n=6, respectively. Two-tailed exact Mann-Whitney U tests. Aβ42, p=0.0303; Aβ42/Aβ40 ratio, p=0.0513. **(I)** Uniform Manifold Approximation and Projection (UMAP) of cortical neuronal nuclei, showing GABAergic and glutamatergic populations. **(J)** Gene set enrichment analysis (GSEA) of curated synaptic and neurodegeneration-associated pathways in cortical single-nucleus RNA-seq data from 12-month-old APP^NL-F^/AhR^+/+^ and APP^NL-F^/AhR^-/-^mice. For each genotype, samples were processed as one pool of 2 males and one pool of 2 females; as only minor sex-dependent differences were observed, nuclei from both sexes were analyzed together. Normalized enrichment scores (NES) were calculated using fgsea; pathways with Benjamini-Hochberg adjusted p<0.05 are shown. WT in the graphs denotes APP^NL-F^/AhR^+/+^. **(K)** Quantification of nest-quality score in 12-month-old APP^NL-F^/AhR^+/+^ and APP^NL-F^/AhR^-/-^mice. n=6 per group. Mann-Whitney U test, p<0.001. **(L)** Percentage of spontaneous alternations in 16-month-old APP^NL-F^/AhR^+/+^ and APP^NL-F^/AhR^-/-^mice. n=12 and n=13, respectively. Two-tailed exact Mann-Whitney U test, p=0.0441. Data are presented as mean±SEM. Each dot represents one biologically independent mouse. Both male and female mice were included in the experimental groups. Sex distribution across experimental groups was as follows: (C–F) APP^NL-F^/AhR^+/+^, 15 males/11 females; APP^NL-F^/AhR^+/−^, 3 males/3 females; APP^NL-F^ /AhR^−/−^, 7 males/6 females; (G,H) APP^NL-F^ /AhR^+/+^, 9 males/5 females; APP^NL-F^ /AhR^−/−^, 3 males/4 females; (K) APP^NL-F^ /AhR^+/+^, 9 males/4 females; APP^NL-F^ /AhR^−/−^, 3 males/3 females; and (L) APP^NL-F^ /AhR^+/+^, 7 males/5 females; APP^NL-F^ /AhR^−/−^, 5 males/8 females. Statistical significance was considered at p<0.05.

To determine whether the reduction in amyloid pathology was accompanied by changes in neuronal transcriptional programs, we performed single-nucleus RNA sequencing of cortical tissue from 12-month-old APP^NL-F^/AhR^+/+^ and APP^NL-F^/AhR^-/-^mice (Figure 2I, Supplementary Figure 3E-J and Supplementary Table 2). Gene set enrichment analysis identified genotype-associated transcriptional differences, with enrichment of pathways related to synaptic transmission, neurotransmitter release and synaptic vesicle exocytosis, together with reduced representation of pathways associated with neuronal apoptosis, cytoskeletal remodelling, and cellular stress (Figure 2J). Additional analyses of GABAergic and glutamatergic neurons, individual differentially expressed genes and sex-stratified transcriptional profiles further supported this neuronal shift (Supplementary Figure 3F-M). Overall, these data indicate that AhR deficiency reshapes neuronal transcriptional programs in parallel with the reduction in amyloid pathology, favouring synaptic signalling over stress- and neurodegeneration-associated programs.

We finally asked whether these changes affected behavioural performance. At 12 months of age, APP^NL-F^/AhR^-/-^mice showed improved nest-building behaviour, with higher nest-quality scores (Figure 2K) and a lower percentage of unused nesting material (Supplementary Figure 3C). Given the late onset of cognitive impairment in the APP^NL-F^ model^25^, working memory was assessed at 16 months. APP^NL-F^/AhR^-/-^mice maintained spontaneous alternation in the Y-maze, whereas APP^NL-F^/AhR^+/+^ mice showed impaired performance (Figure 2L and Supplementary Figure 3D). Together, these findings indicate that AhR loss improves species-typical behaviour and is associated with preserved working-memory performance.

### AhR regulates plaque-associated microglial responses and amyloid accumulation

We next examined whether AhR expression changes during disease progression. Cortical AhR protein was higher in APP^NL-F^ than WT mice at both 5 and 12 months and increased from 5 to 12 months in APP^NL-F^ mice, whereas no significant age-related increase was detected in WT mice (Supplementary Figure 4A). Consistently, immunofluorescence showed stronger cortical AhR labelling at this stage (Supplementary Figure 4B-C). AhR was prominently expressed in Iba1-positive microglia and GFAP-positive astrocytes, particularly around amyloid plaques (Figure 3A), whereas overlap with NeuN-positive neurons was limited (Supplementary Figure 4C).

**Figure 3.**
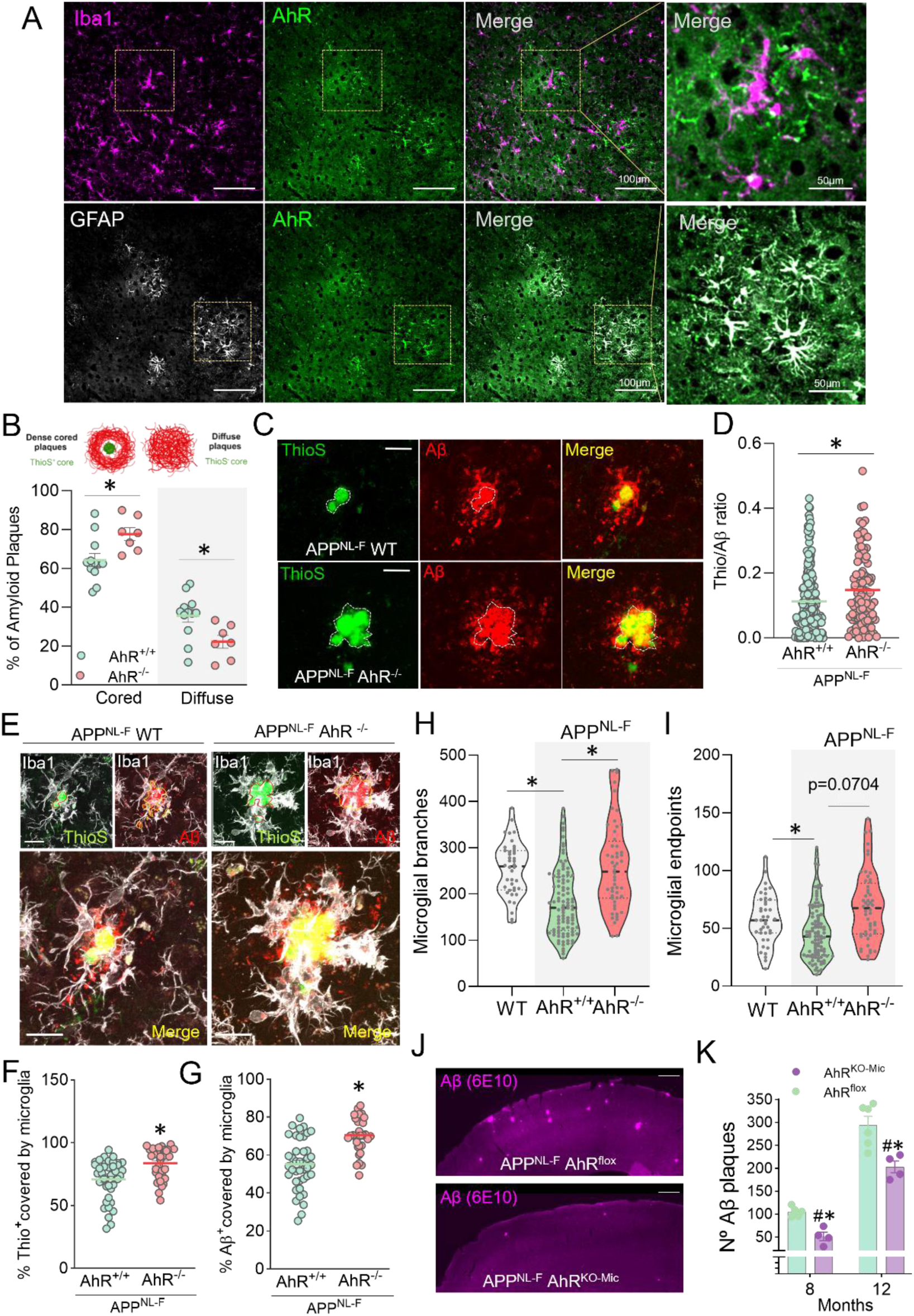
AhR is enriched in plaque-associated glia and regulates microglial responses and amyloid accumulation in APP^NL-F^ mice. **(A)** Representative cortical images from 12-month-old APP^NL-F^ mice showing AhR immunoreactivity in Iba1-positive microglia and GFAP-positive astrocytes. Enlarged views show AhR expression in plaque-associated glial cells. Scale bars = 100 μm; enlarged images, 50 μm. **(B)** Classification and quantification of dense-cored and diffuse amyloid plaques in 12-month-old APP^NL-F^/AhR^+/+^ and APP^NL-F^/AhR^-/-^mice. Dense-cored plaques were defined as Aβ-positive deposits containing a ThioS-positive core, whereas diffuse plaques lacked a ThioS-positive core. APP^NL-F^/AhR^+/+^: n=12 mice; APP^NL-F^/AhR^-/-^: n=7 mice. Two-way ANOVA followed by Sidák’s multiple-comparisons test. For cored plaques and for diffuse plaques adjusted p=0.0246. **(C)** Representative images of ThioS-positive fibrillar cores and total Aβ-positive deposits in APP^NL-F^/AhR^+/+^ and APP^NL-F^/AhR^-/-^mice. Scale bars=10μm. **(D)** ThioS/Aβ ratio. Individual plaque measurements are shown for visualization, whereas values were averaged within each mouse for statistical inference. APP^NL-F^/AhR^+/+^: n=12 mice; APP^NL-F^/AhR^−/−^: n=6 mice. Two-sided Welch’s t-test on mouse-level means, p=0.0303. **(E)** Representative images showing Iba1-positive microglial processes surrounding ThioS-positive or Aβ-positive plaques in APP^NL-F^/AhR^+/+^ and APP^NL-F^/AhR^-/-^ mice. Scale bars=10μm. **(F)** Percentage of the ThioS-positive plaque area covered by Iba1-positive microglia, quantified in 39 plaques from n=12 APP^NL-F^/AhR^+/+^ mice and 32 plaques from n=7 APP^NL-F^/AhR^-/-^mice. Two-tailed exact Mann-Whitney U test, p=0.0004. **(G)** Percentage of the Aβ-positive plaque area covered by Iba1-positive microglia, quantified in 41 plaques from n=12 APP^NL-F^/AhR^+/+^ mice and 32 plaques from n=7 APP^NL-F^/AhR^-/-^. Two-tailed exact Mann-Whitney U test, p<0.0001. **(H)** Number of branches in cortical microglia located away from amyloid plaques. Individual-cell measurements are shown for visualization, whereas values were averaged within each mouse for statistical inference. WT: n=4 mice, 40 cells; APP^NL-F^/AhR^+/+^: n=12 mice, 107 cells; APP^NL-F^/AhR^-/-^: n=6 mice, 52 cells. Pairwise two-sided Welch’s t-tests on mouse-level means with Holm correction: WT versus APP^NL-F^/AhR^+/+^, adjusted p=0.0028; WT versus APP^NL-F^/AhR^-/-^, adjusted p=0.9814; APP^NL-F^/AhR^+/+^ versus APP^NL-F^/AhR^-/-^adjusted p=0.0490. **(I)** Number of endpoints in cortical microglia located away from amyloid plaques. Individual-cell measurements are shown for visualization, whereas values were averaged within each mouse for statistical inference. WT: n=4 mice, 40 cells; APP^NL-F^/AhR^+/+^: n=12 mice, 107 cells; APP^NL-F^/AhR^-/-^: n=6 mice, 52 cells. Pairwise two-sided Welch’s t-tests on mouse-level means with Holm correction: WT versus APP^NL-F^/AhR^+/+^, adjusted p=0.0583; WT versus APP^NL-F^/AhR^-/-^, adjusted p=0.3074; APP^NL-F^/AhR^+/+^versus APP^NL-F^/AhR^-/-^, adjusted p=0.0704 (unadjusted p=0.0352). **(J)** Representative coronal brain sections showing Aβ immunoreactivity detected with the 6E10 antibody in control APP^NL-F^/AhR^fl/fl^ mice and APP^NL-F^ mice with tamoxifen-induced microglia-specific, Cx3cr1-CreER-mediated AhR deletion (AhR^KO-Mic^). Scale bars=250μm. **(K)** Number of cortical Aβ-positive plaques in APP^NL-F^/AhR^fl/fl^ and APP^NL-F^/ AhR^KO-Mic^ mice at 8 and 12 months of age. At 8 months, n=5 mice per group; at 12 months, n=6 mice per group. Two-way ANOVA followed by Šídák’s multiple-comparisons test. At 8 months, p=0.0379; at 12 months, p=0.0006. Data are presented as mean±SEM. Each dot represents one mouse in **(B)** and **(K)**, one plaque in **(D)**, **(F)** and **(G)**, or one microglial cell in **(H)** and **(I)**. For panels D, H and I, individual plaque or cell measurements are shown for visualization, whereas statistical inference was performed on values averaged within each mouse, with the mouse considered the experimental unit. Both male and female mice were included in the experimental groups. Sex distribution across experimental groups was as follows: (B) APP^NL-F^/AhR^+/+^, 6 males/6 females; APP^NL-F^/AhR^−/−^, 3 males/4 females; and (K) at 8 months, APP^NL-F^/AhR^fl/fl^, 1 male/4 females; APP^NL-F^/AhR^KO-Mic^, 2 males/2 females; at 12 months, APP^NL-F^/AhR^fl/fl^, 2 males/4 females; APP^NL-F^ /AhR^KO-Mic^, 2 males/2 females. Statistical significance was considered at p<0.05.

Given the prominent enrichment of AhR in plaque-associated microglia, and the central role of microglia in shaping plaque architecture and compaction^1, 12, 26, 27, 28, 29^, we next asked whether AhR regulates plaque organization and the microglial response to amyloid deposits. APP^NL-F^/AhR^-/-^ mice showed a shift toward a higher proportion of dense-cored plaques and fewer diffuse deposits (Figure 3B-C), together with an increased ThioS/Aβ ratio, consistent with enhanced plaque compaction (Figure 3D). Constitutive AhR loss was also associated with increased Iba1-positive coverage of both ThioS-positive cores and total Aβ deposits (Figure 3E-G and Supplementary Figure 5B-C). No significant difference was detected in overall cortical microglial abundance (Supplementary Figure 5D-E). Microglia located away from plaques from APP^NL-F^/AhR^-/-^ mice also showed a more ramified morphology, with increased numbers of branches and a similar trend in endpoints compared with APP^NL-F^/AhR^+/+^ mice, with values like those observed in non-AD WT mice (Figure 3H-I and Supplementary Figure 5F). Together, these findings suggest that AhR may regulate plaque organization through its effects on plaque-associated microglia.

To investigate whether targeting AhR in the microglial compartment was sufficient to modify amyloid pathology, we generated APP^NL-F^ mice with tamoxifen-inducible Cx3cr1-CreER-mediated deletion of Ahr (AhR^KO-Mic^). Cx3cr1-lineage tracing confirmed Cre activity in microglia and revealed reduced AhR immunoreactivity, with a similar trend toward reduced AhR protein levels observed by Western blot in isolated microglia (p=0.1143; Supplementary Figure 5G-H). AhR^KO-Mic^ mice exhibited significantly reduced cortical Aβ plaque burden at both 8 and 12 months of age (Figure 3J-K). Together, these findings identify AhR as an important regulator of plaque-associated microglial responses and demonstrate that its deletion in C x3cr1-expressing cells reduces amyloid accumulation *in vivo*.

### AhR loss alters phagolysosomal microglial programs and Aβ handling in a sex-dependent manner

To define the microglial programs regulated by AhR during amyloid pathology, we performed single-cell RNA sequencing of cortical non-neuronal cells from 12-month-old male and female APP^NL-F^/AhR^+/+^ and APP^NL-F^/AhR^-/-^ mice (Figure 4A). Following Percoll enrichment and isolation of viable DAPI-negative cells, 8,604 high-quality cells were retained for analysis. Unsupervised clustering identified 19 transcriptional clusters corresponding to the major non-neuronal brain populations (Figure 4B, Supplementary Figure 6 and Supplementary Table 3). Microglia represented the predominant population across all groups, accounting for 65.3-80.4% of analysed cells (Figure 4C).

**Figure 4.**
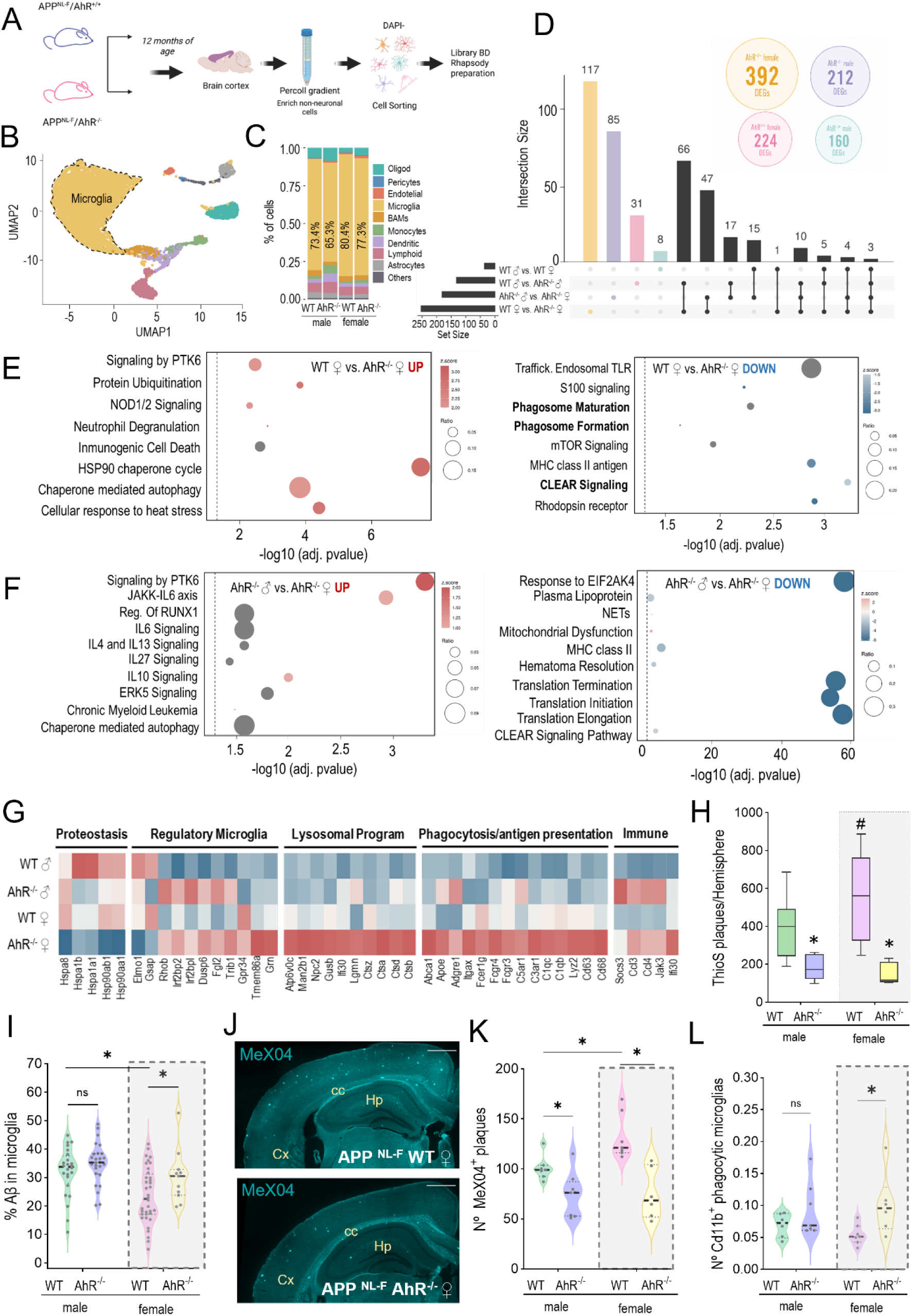
AhR loss alters phagolysosomal programs and Aβ handling in APP^NL-F^ mice, in a sex-dependent fashion. **(A)** Experimental workflow for single-cell RNA sequencing. Cortical non-neuronal cells were isolated from 12-month-old male and female APP^NL-F^/AhR^+/+^ and APP^NL-F^/AhR^-/-^mice by Percoll-gradient enrichment, followed by sorting of viable DAPI-negative cells and library preparation using the BD Rhapsody system. **(B)** UMAP representation of 8,604 cells showing 19 transcriptional clusters assigned to the major non-neuronal brain cell populations. **(C)** Relative abundance of the identified cell populations across sex and genotype. **(D)** Differential gene-expression analysis of microglia according to AhR genotype and biological sex. Circles indicate the number of DEGs identified for each sex-genotype group compared with all other groups combined (one-versus-rest analysis). The UpSet plot shows the total number of DEGs and their overlap across four independent pairwise comparisons: APP^NL-F^/AhR^-/-^versus APP^NL-F^/AhR^+/+^ in females, APP^NL-F^/AhR^-/-^versus APP^NL-F^/AhR^+/+^ in males, females versus males in AhR^-/-^mice, and females versus males in APP^NL-F^/AhR^+/+^ mice. Horizontal bars indicate the total number of DEGs in each comparison, whereas vertical bars represent genes unique to or shared among the comparisons indicated by the connected dots. **(E)** Functional enrichment analysis of genes upregulated and downregulated in female APP^NL-F^/AhR^-/-^ compared with female APP^NL-F^/AhR^+/+^ microglia. **(F)** Functional enrichment analysis of genes upregulated and downregulated between male and female APP^NL-F^/AhR^-/-^. Dot size represents the gene ratio, colour indicates the pathway activation z-score and the x axis shows −log 10 (adjusted p-value). **(G)** Heatmap showing the expression of selected genes associated with proteostasis, microglial regulation, lysosomal function, phagocytosis and antigen presentation, and immune signalling in male and female APP^NL-F^/AhR^+/+^ and APP^NL-F^/AhR^-/-^ microglia. Values represent row-scaled z-scores of average gene expression. **(H)** Cortical ThioS-positive plaque burden in male and female APP^NL-F^/AhR^+/+^ and APP^NL-F^/AhR^-/-^ mice. Male APP^NL-F^/AhR^+/+^: n=15; male APP^NL-F^/AhR^-/-^: n=8; female APP^NL-F^/AhR^+/+^: n=10; female APP^NL-F^/AhR^-/-^: n=6. Two-way ANOVA followed by Tukey’s multiple-comparisons test. APP^NL-F^/AhR^+/+^ versus APP^NL-F^/AhR^-/-^: males, p=0.0247; females, p=0.0002. Male versus female APP^NL-F^/AhR^+/+^ mice, p=0.0553. **(I)** Percentage of Aβ immunoreactivity contained within Iba1-positive microglia in male and female APP^NL-F^/AhR^+/+^ and APP^NL-F^/AhR^-/-^ mice. A total of 24, 26, 36 and 11 plaques were analysed, respectively, from male APP^NL-F^/AhR^+/+^: n=6; male APP^NL-F^/AhR^-/-^: n=6; female APP^NL-F^/AhR^+/+^: n=4; female APP^NL-F^/AhR^-/-^: n=2. Two-way ANOVA followed by Fisher’s LSD test. APP^NL-F^/AhR^+/+^ versus APP^NL-F^/AhR^-/-^: males, p=0.2330; females, p=0.0289. Male versus female APP^NL-F^/AhR^+/+^ mice, p=0.0004. Sex-stratified analyses in this panel were exploratory and performed post hoc following the observation of sex-dependent differences in the microglial response to AhR loss; interpretation of the female comparison is additionally limited by the small number of AhR-deficient females (n=2). **(J)** Representative brain sections from female APP^NL-F^/AhR^+/+^ and APP^NL-F^/AhR^-/-^ mice showing fibrillar Aβ deposits labelled in vivo with Methoxy-X04. Cx, cortex; cc, corpus callosum; Hp, hippocampus. Scale bar = 500 μm. **(K)** Number of cortical Methoxy-X04-positive plaques in male and female APP^NL-F^/AhR^+/+^ and APP^NL-F^/AhR^-/-^ mice. Male APP^NL-F^/AhR^+/+^: n=7; male APP^NL-F^/AhR^-/-^: n=8; female APP^NL-F^/AhR^+/+^: n=7; female APP^NL-F^/AhR^-/-^: n=6. Two-way ANOVA followed by Fisher’s LSD test. APP^NL-F^/AhR^+/+^versus APP^NL-F^/AhR^-/-^: males, p=0.0224; females, p<0.0001. Male versus female APP^NL-F^/AhR^+/+^ mice, p=0.0107. **(L)** Flow-cytometric quantification of CD11b-positive microglia containing Methoxy-X04-labeled Aβ in male and female APP^NL-F^/AhR^+/+^ and APP^NL-F^/AhR^-/-^ mice. Male APP^NL-F^/AhR^+/+^: n=6; male APP^NL-F^/AhR^-/-^: n=7; female APP^NL-F^/AhR^+/+^: n=7; female APP^NL-F^/AhR^-/-^: n=6. Two-way ANOVA followed by Fisher’s LSD test. APP^NL-F^/AhR^+/+^ versus APP^NL-F^/AhR^-/-^: males, p=0.2141; females, p=0.0226. Data are presented as mean±SEM. Statistical significance was considered at p<0.05.

We therefore focused on the microglial compartment. Pairwise differential gene-expression analysis revealed a greater number of genotype-associated transcriptional differences in female microglia, with 253 differentially expressed genes (DEGs) between AhR-deficient and AhR-intact females, compared with 132 in males (Figure 4D). Similarly, comparisons between sexes identified 181 DEGs in AhR-deficient microglia, compared with only 36 in AhR-intact microglia (Figure 4D). Notably, UpSet analysis revealed that the female genotype comparison also contained the largest number of comparison-specific DEGs (117 genes, compared with 31 in males), alongside transcriptional changes shared across comparisons (Figure 4D; Supplementary Figures 7-8).

Functional enrichment revealed that AhR loss in female microglia strongly enhanced pathways related to phagosome formation and maturation, lysosomal function and CLEAR signalling (Figure 4E). Comparison of AhR-deficient males and females also identified lysosomal and clearance-associated transcriptional differences (Figure 4F). At the gene level, this response included coordinated upregulation of Aβ-recognition and uptake genes, including *Apoe, Itgax, Adgre1, Fcgr3, Fcgr4, C1qb, C1qc, C3ar1* and *C5ar1*, together with phagolysosomal effectors such as *Cd68, Cd63, Lyz2, Ctsb, Ctsd, Ctsa, Ctsz, Lgmn* and *Ifi30* (Figure 4G). This program was less prominent in males, which retained comparatively higher expression of inflammatory and stress-associated mediators, including *Jak3, Ccl4* and *Socs3*. Thus, constitutive AhR loss is associated with an Aβ-handling and phagolysosomal program that is substantially stronger in female microglia.

We next asked whether this transcriptional response was accompanied by functional changes in amyloid handling. Constitutive AhR deletion reduced cortical ThioS-positive plaque burden in both sexes, with a larger reduction observed in females (Figure 4H). Consistent with this pattern, an exploratory quantification of Aβ immunoreactivity within Iba1-positive microglia showed increased intramicroglial Aβ in AhR-deficient females, whereas no significant increase was detected in males (Figure 4I). To independently assess microglial uptake of fibrillar Aβ *in vivo*, mice were administered Methoxy-X04 before sacrifice. Methoxy-X04 imaging confirmed reduced cortical fibrillar plaque burden following AhR loss, again with a particularly pronounced reduction in females (Figure 4J-K). Flow cytometry further showed an increased number of CD11b-positive microglia containing Methoxy-X04-labeled Aβ in female APP^NL-F^/AhR^-/-^mice (Figure 4L and Supplementary Figure 9). Together, these findings show that AhR loss promotes a phagolysosomal and Aβ-handling microglial program that is particularly prominent in females. This transcriptional state is accompanied by enhanced microglial Aβ engulfment, suggesting a functional link between AhR-dependent microglial reprogramming and reduced amyloid pathology.

To further resolve the microglial states associated with AhR loss, we reclustered the microglial compartment and identified eight transcriptionally distinct subpopulations, MgC0-MgC7 (Figure 5A-B). Each cluster displayed a characteristic marker-gene profile (Figure 5C). Integration with previously defined microglial gene modules further identified MgC4 as a DAM-like population, MgC5 as an interferon-responsive state, MgC7 as a cycling/proliferative population and MgC2 as an immediate-early response state (Supplementary Figure 10). By contrast, MgC0 and MgC1 shared a broad transcriptional core while retaining distinct gene-expression profiles, suggesting related but transcriptionally distinguishable microglial states (Supplementary Figure 10-11). MgC0 and MgC1 were the two predominant microglial populations. AhR-deficient samples showed a lower relative representation of MgC0 and a higher representation of MgC1 in both sexes, consistent with altered microglial cluster composition (Figure 5D). Functional enrichment revealed that MgC0 was associated with leukocyte adhesion and migration, cytoskeletal organization, wound healing, and myeloid-cell activation (Figure 5E). In contrast, MgC1 was enriched in chaperone-mediated autophagy, endocytosis, Fc-receptor signalling, phagocytosis and pathways related to amyloid precursor protein processing (Figure 5F). Thus, these data strongly suggest that AhR loss redistributes microglia toward an MgC1-associated state enriched in pathways linked to endocytosis, amyloid recognition, uptake, and intracellular processing.

**Figure 5.**
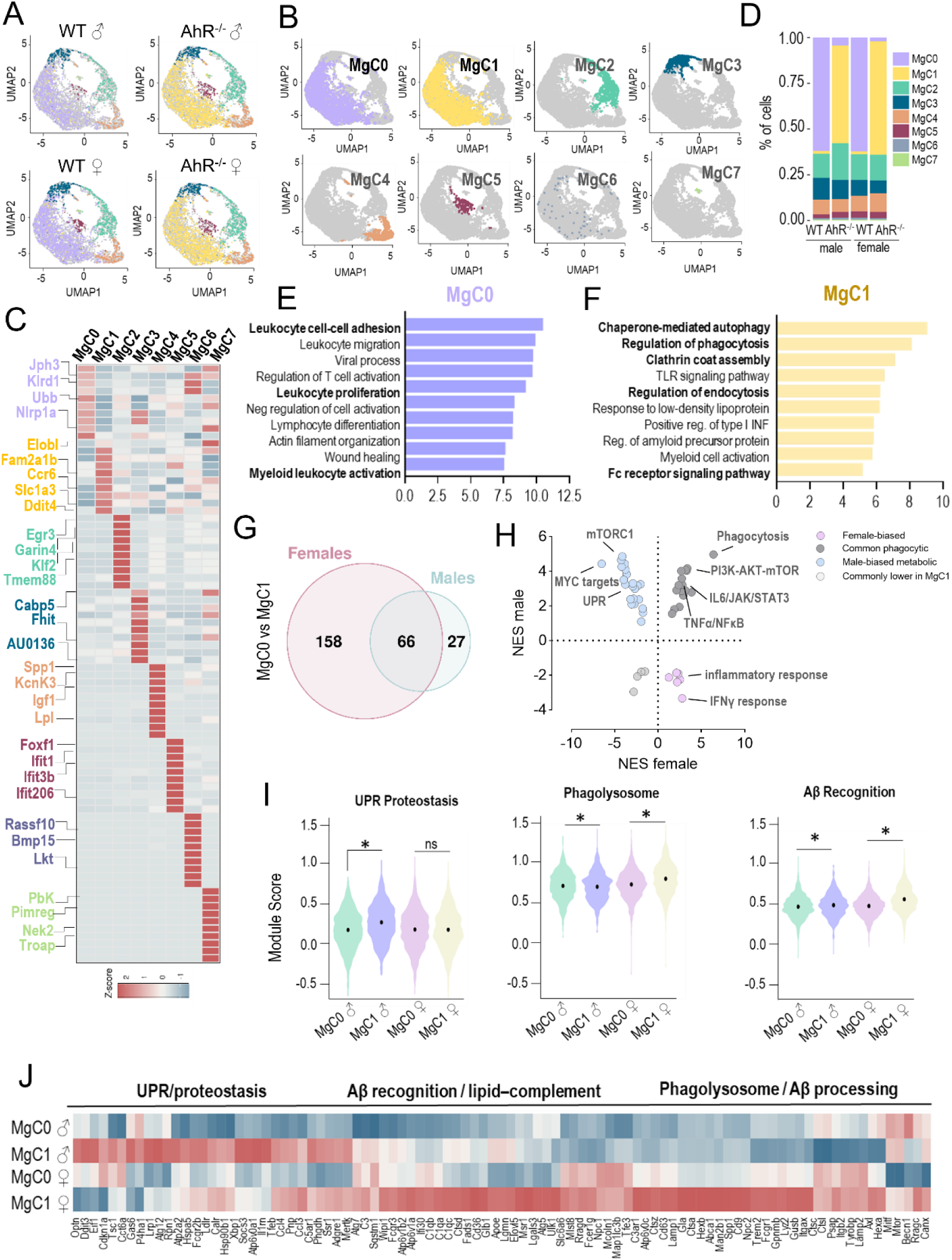
AhR loss is associated with altered proportions of microglial states in APP^NL-F^ mice. **(A)** UMAP representation of reclustered microglia from 12-month-old male and female APP^NL-F^/AhR^+/+^ and APP^NL-F^/AhR^-/-^ mice, separated by sex and genotype. **(B)** UMAP projections showing the distribution of the eight microglial subpopulations, designated MgC0-MgC7. **(C)** Heatmap showing representative marker genes defining MgC0-MgC7. Values represent row-scaled z-scores of gene expression. **(D)** Relative abundance of MgC0-MgC7 across sex and genotype. **(E, F)** Functional enrichment analysis of genes associated with MgC0 **(E)** and MgC1 **(F)**. Bars represent −log10 adjusted p value/gene ratio. **(G)** Venn diagram showing differentially expressed genes identified in the MgC1-versus-MgC0 comparisons in females and males. A total of 66 genes were shared between sexes, whereas 158 reached significance only in females and 27 only in males. **(H)** Comparison of normalized enrichment scores for pathways identified in the female and male MgC1-versus-MgC0 contrasts. Pathways are classified as female-biased, male-biased, commonly phagocytic, commonly enriched in MgC1 or commonly reduced in MgC1, as indicated. **(I)** UPR/proteostasis, phagolysosome and Aβ-recognition module scores in MgC0 and MgC1 from male and female mice. **(J)** Heatmap showing selected genes associated with UPR/proteostasis, Aβ recognition and lipid-complement signalling, and phagolysosomal Aβ processing in male and female MgC0 and MgC1. Values represent row-scaled z-scores of average gene expression.

We next examined whether the MgC1 program differed between sexes. Comparison of MgC1 and MgC0 identified 66 differentially expressed genes shared between males and females, whereas 158 were significant only in females and 27 only in males (Figure 5G). At the pathway level, both sexes showed enrichment of phagocytosis and PI3K-AKT-mTOR-related signalling. Male MgC1 showed enrichment of mTORC1, MYC-target and unfolded-protein-response pathways, whereas females showed relatively greater enrichment of inflammatory and interferon-response programs (Figure 5H). Module scoring further resolved these differences. MgC1 displayed higher phagolysosomal and Aβ-recognition scores than MgC0 in both sexes, whereas increased UPR/proteostasis signaling was restricted to males (Figure 5I). Consistently, gene-level analysis revealed a shared induction of phagolysosomal and Aβ-handling genes, superimposed on a stronger proteostatic-stress signature in males and a broader lipid-complement and amyloid-recognition program in females (Figure 5J). Together, these findings show that AhR loss shifts microglia toward a shared Aβ-processing state, with distinct sex-dependent transcriptional specialization: broader amyloid-recognition and lipid-complement remodelling in females, and greater proteostatic stress in males.

### AhR-activating bioactivity is increased in AD mice and in an independently recruited human memory-clinic subcohort

Given the marked, sex-dependent effects of AhR on microglial responses and amyloid pathology, we next asked whether the endogenous AhR-activating environment was also altered in APP^NL-F^ mice. To address this, we quantified AhR-activating bioactivity using an XRE-driven luciferase reporter assay (Figure 6A), whose responsiveness was confirmed using the established AhR agonists FICZ and I3S (Supplementary Figure 12). Serum from male and female APP^NL-F^ mice and cortical extracts from female APP^NL-F^ mice showed increased AhR-activating bioactivity than samples from their respective sex-matched WT controls (Figure 6B-C). Thus, amyloid pathology was associated with enhanced AhR-activating bioactivity in peripheral and central compartments, with a sex-dependent pattern that differed between serum and cortex.

**Figure 6.**
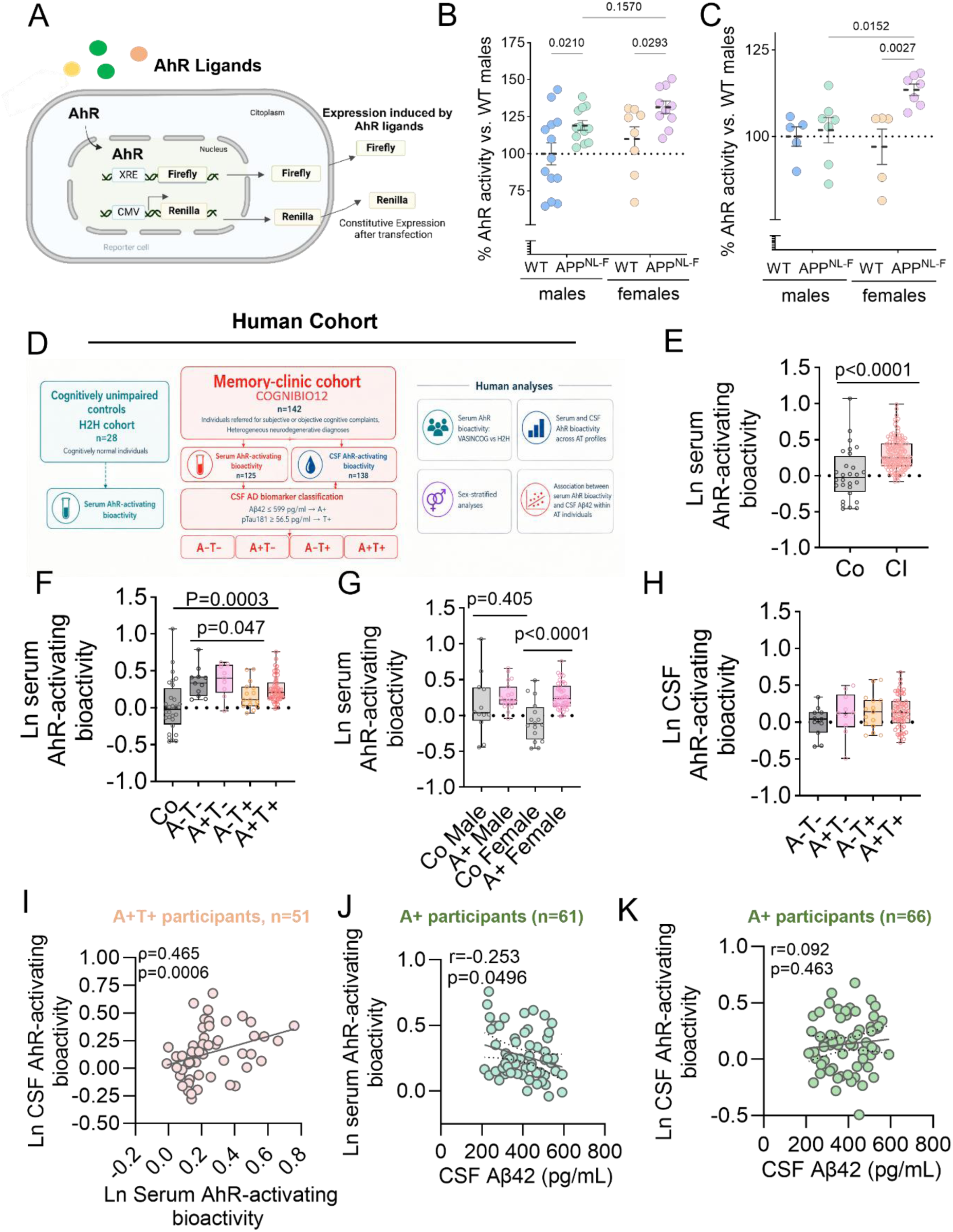
AhR-activating bioactivity is increased in APP^NL-F^ mice and the COGNIBIO12 analytical subcohort compared with independent H2H controls. **(A)** Schematic representation of the XRE-driven dual-luciferase reporter assay used to quantify AhR-activating bioactivity. AhR activation induces firefly luciferase expression through xenobiotic response elements (XREs), whereas constitutively expressed Renilla luciferase was used as an internal normalization control. **(B)** Serum AhR-activating bioactivity in male and female WT and APP^NL-F^ mice. WT males: n=13; APP^NL-F^ males: n=12; WT females: n=8; APP^NL-F^ females: n=10. Reporter activity was expressed relative to the mean value of WT males, set to 100%. Two-way ANOVA followed by uncorrected Fisher’s LSD multiple-comparisons test. WT versus APP^NL-F^: males, p=0.0210; females, p=0.0293. Male versus female: WT, p=0.2658; APP^NL-F^, p=0.1570. Each dot represents one biologically independent mouse. **(C)** AhR-activating bioactivity induced by cortical extracts from male and female WT and APP^NL-F^ mice. WT males: n=5; APP^NL-F^ males: n=7; WT females: n=5; APP^NL-F^ females: n=7. Reporter activity was expressed relative to the mean value of WT males, set to 100%. Two-way ANOVA followed by uncorrected Fisher’s LSD multiple-comparisons test. WT versus APP^NL-F^: females, p=0.0027. Male versus female APP^NL-F^ mice, p=0.0152. Each dot represents one biologically independent mouse. **(D)** Schematic overview of the human cohorts and analyses. Cognitively unimpaired participants from the H2H cohort (n=28) constituted the primary control group. The COGNIBIO12 analytical subcohort comprised 142 individuals from the larger parent COGNIBIO12 cohort (>300 enrolled) referred for subjective or objective cognitive complaints and included heterogeneous neurodegenerative diagnoses. Serum AhR-activating bioactivity was available for n=125 COGNIBIO12 participants and CSF AhR-activating bioactivity for n=138; 122 participants had matched serum and CSF AhR measurements, three had serum alone, 16 had CSF alone and one had neither valid measurement. Participants with available CSF AD biomarkers were classified according to amyloid (A) and tau (T) status using CSF Aβ42≤599 pg/mL for A+ and CSF p-tau181≥56.5 pg/mL for T+, generating A−T−, A+T−, A−T+ and A+T+ groups (n=93 with both biomarkers; serum AhR available in 87 and CSF AhR in 92). CSF Aβ40 was available for 69 participants, and age and sex were recorded for 137 and 139 participants, respectively. **(E)** Serum AhR-activating bioactivity in cognitively unimpaired H2H controls and participants from the COGNIBIO12 clinical cohort. H2H: n=28; COGNIBIO12: n=125. Serum AhR bioactivity was natural-log transformed and analysed using linear regression adjusted for age and sex with HC3 robust standard errors. The COGNIBIO12 versus H2H between-cohort contrast yielded an adjusted geometric mean ratio of 1.330 (95% CI 1.162–1.523; p<0.0001). **(F)** Serum AhR-activating bioactivity in H2H controls and COGNIBIO12 participants stratified according to CSF amyloid and tau biomarker status. H2H: n=28; A−T−: n=11; A+T−: n=9; A−T+: n=15; A+T+: n=52. The within-COGNIBIO12 adjusted global effect across the four AT groups (serum n=87, H2H excluded) was significant (p=0.0181). Planned contrasts against A−T− were corrected using the Holm procedure; A−T+ participants showed lower serum AhR bioactivity than A−T− participants (adjusted geometric mean ratio=0.830, 95% CI 0.714-0.965, Holm-adjusted p=0.0470), whereas A+T− and A+T+ did not differ significantly from A−T−. Relative to H2H controls, A+T+ participants showed higher serum AhR bioactivity (adjusted geometric mean ratio=1.276; 95 % IC 1.117-1.457; p=0.0003). **(G)** Serum AhR-activating bioactivity in amyloid-positive participants and sex-matched H2H controls. H2H males: n=12; A+ males: n=19; H2H females: n=16; A+ females: n=42. Age-adjusted analyses showed no significant difference between A+ and H2H males (adjusted geometric mean ratio=1.115, 95% CI 0.863-1.442, p=0.4050), whereas A+ females showed higher serum AhR bioactivity than H2H females (adjusted geometric mean ratio=1.409, 95% CI 1.205-1.647, p<0.0001). The formal group-by-sex interaction was not significant (p=0.1298). **(H)** CSF AhR-activating bioactivity in COGNIBIO12 participants stratified according to amyloid and tau biomarker status. A−T−: n=11; A+T−: n=10; A−T+: n=15; A+T+: n=56. No significant global difference was detected across the four AT groups (CSF AhR available in n=92; age- and sex-adjusted model n=91; global p=0.3482). In the adjusted model, A+T+ participants showed a positive but non-significant estimate relative to A−T− participants (adjusted geometric mean ratio=1.138, 95% CI 0.987-1.312, unadjusted p=0.0756, Holm-adjusted p=0.2268). One A+T+ participant lacked age data and was therefore excluded from the adjusted model (adjusted-model n=91). **(I)** Association between serum and CSF AhR-activating bioactivity among A+T+ participants. n=51. Serum and CSF AhR bioactivities were positively associated (Pearson correlation on natural-log-transformed values, r=0.328, p=0.0188; Spearman ρ=0.465, p=0.0006). The association remained significant after adjustment for age and sex (p=0.0019, Benjamini–Hochberg FDR q=0.0241). However, the formal interaction with A+T+ status was not significant (p=0.2603). The broader A+ matched subset comprised n=60 (Spearman ρ=0.378, p=0.0029), whereas the plotted A+T+ panel comprised n=51. **(J)** Association between CSF Aβ42 concentration and serum AhR-activating bioactivity among amyloid-positive participants. n=61. Pearson correlation between natural-log-transformed serum AhR bioactivity and CSF Aβ42: r=−0.253, p=0.0496. The solid line represents the linear regression fit and dotted lines indicate the 95% confidence interval. The association retained the same direction after adjustment for age and sex but did not reach statistical significance (adjusted geometric mean ratio per 1-SD increase in CSF Aβ42=0.955, 95% CI 0.906-1.006, p=0.0849). Spearman ρ=−0.200, p=0.1220. **(K)** Association between CSF Aβ42 concentration and CSF AhR-activating bioactivity among amyloid-positive participants. n=66. Pearson correlation on natural-log-transformed CSF AhR bioactivity, r=0.092, p=0.4630. The solid line represents the linear regression fit and dotted lines indicate the 95% confidence interval. No association was detected after adjustment for age and sex (adjusted geometric mean ratio per 1-SD increase in CSF Aβ42=1.022, 95% CI 0.970-1.077, p=0.4120). Individual dots represent individual participants. Amyloid positivity was defined as CSF Aβ42≤599 pg/mL and tau positivity as CSF p-tau181≥56.5 pg/mL. Statistical tests were two-sided and p<0.05 was considered statistically significant.

We next asked whether an AhR-activating milieu was also detectable in humans. We analysed 142 participants from the larger COGNIBIO12 memory-clinic cohort (>300 enrolled) as an analytical subcohort evaluated for subjective or objective cognitive complaints and heterogeneous clinical diagnoses, alongside 28 cognitively unimpaired participants from the independent H2H cohort as an external reference group (Figure 6D). Serum AhR bioactivity was available for 125 COGNIBIO12 participants and 28 H2H controls, whereas CSF AhR bioactivity was available for 138 COGNIBIO12 participants. Clinical and biomarker characteristics are summarized in Table 1 and Supplementary Table 4-5.

**Table 1.** Demographic, clinical and biomarker characteristics of the human study cohorts. Demographic, clinical and biomarker characteristics of cognitively unimpaired participants from the H2H cohort and the 142-person analytical subcohort of the parent COGNIBIO12 clinical cohort (>300 enrolled). Age is presented as mean ± SD. Continuous biomarker and clinical variables are summarized as median [interquartile range, IQR] because several showed skewed or non-normal distributions. Categorical variables are presented as n/available n (%). The n indicated for each variable corresponds to participants with available data. An amyloid aetiology was recorded separately in 87/142 participants (61.3%), including 80 who also had a clinical Alzheimer’s disease diagnosis; clinical diagnosis and etiological classification overlapped and are therefore reported separately. Clinical diagnostic categories were defined using the clinical-diagnosis field and are mutually exclusive; 85/142 participants (59.9%) had a clinical Alzheimer’s disease code. The detailed clinical diagnostic distribution is provided in the cohort Methods. Sex was unrecorded for three COGNIBIO12 participants. Of the 142 COGNIBIO12 participants, 125 had analytically valid serum AhR measurements and 138 had CSF AhR measurements; 122 participants had paired serum and CSF results, three had serum only, 16 had CSF only and one had neither valid measurement. Serum AhR-activating bioactivity was available in both cohorts, whereas CSF AhR-activating bioactivity and CSF AD biomarkers were available only in COGNIBIO12 participants. AhR-activating bioactivity was determined using an XRE-driven luciferase reporter assay. Age was available for 137 participants and sex for 139. CSF Aβ42 and p-tau181 were available in 93 participants and CSF Aβ40 in 69; participants with available Aβ42 and p-tau181 measurements were classified according to AT status. Amyloid positivity was defined as CSF Aβ42 ≤599 pg/mL and tau positivity as CSF p-tau181 ≥56.5 pg/mL. Visit-matched MMSE and GDS assessments were available for 42 and 91 participants, respectively; values from other longitudinal visits were not substituted. Among participants with available GDS data, the stage distribution was: GDS 2, n=1; GDS 3, n=55; GDS 4, n=33; and GDS 5, n=2. The H2H group comprised 28 independently recruited cognitively unimpaired participants with serum AhR measurements only. Missing values were not imputed; each analysis included participants with available data for the variables required by that analysis. MMSE, Mini-Mental State Examination; GDS, Global Deterioration Scale; CSF, cerebrospinal fluid; AhR, aryl hydrocarbon receptor; IQR, interquartile range; SD, standard deviation.

| Characteristic | H2H cognitively unimpaired controls | COGNIBIO12 analytical subcohort | Women, n | Women age, years (mean ± SD) | Men, n | Men age, years (mean ± SD) |
|---|---|---|---|---|---|---|
| Participants, n | 28 | 142 | 82 | — | 57 | — |
| Age, years | 73.5 ± 5.3 (n=28) | 74.8 ± 7.1 (n=137) | — | 74.2 ± 7.1 (n=82) | — | 75.5 ± 7.1 (n=55) |
| Women, n/available (%) | 16/28 (57.1%) | 82/139 (59.0%) | — | — | — | — |
| Men, n/available (%) | 12/28 (42.9%) | 57/139 (41.0%) | — | — | — | — |
| Sex unavailable, n | 0 | 3 | — | — | — | — |
| **Clinical diagnosis and sex-stratified age** |  |  |  |  |  |  |
| Alzheimer's disease | — | 85 (59.9%) | 59 | 76.0 ± 5.3 (n=59) | 26 | 76.5 ± 4.4 (n=25) |
| Dementia with Lewy bodies | — | 10 (7.0%) | 4 | 78.3 ± 5.8 (n=4) | 6 | 80.4 ± 5.6 (n=6) |
| Primary progressive aphasia | — | 9 (6.3%) | 5 | 75.3 ± 7.4 (n=5) | 4 | 79.1 ± 3.8 (n=4) |
| Behavioral-variant frontotemporal dementia | — | 7 (4.9%) | 4 | 63.1 ± 11.0 (n=4) | 3 | 73.7 ± 0.9 (n=3) |
| Mixed clinical diagnosis | — | 4 (2.8%) | 1 | 67.6 (n=1) | 3 | 83.1 ± 0.8 (n=3) |
| Vascular cognitive impairment | — | 2 (1.4%) | 0 | — | 2 | 77.8 ± 2.4 (n=2) |
| Posterior cortical atrophy | — | 2 (1.4%) | 2 | 65.1 ± 6.3 (n=2) | 0 | — |
| Corticobasal syndrome | — | 1 (0.7%) | 0 | — | 1 | 76.6 (n=1) |
| Other clinical diagnoses | — | 13 (9.2%) | 4 | 63.2 ± 6.5 (n=4) | 9 | 66.3 ± 10.3 (n=9) |
| Clinical diagnosis unavailable | — | 9 (6.3%) | 3 | 71.0 ± 6.6 (n=3) | 3 | 71.8 ± 4.8 (n=2) |
| Amyloid etiology, n/available (%)¹ | — | 87/142 (61.3%)¹ | 60 | 75.8 ± 5.5 (n=60) | 27 | 76.7 ± 4.3 (n=26) |
| **Sample availability** |  |  |  |  |  |  |
| Serum AhR-activating bioactivity available, n | 28 | 125 | — | — | — | — |
| CSF AhR-activating bioactivity available, n | — | 138 | — | — | — | — |
| Paired serum + CSF AhR measurements, n | — | 122 | — | — | — | — |
| Serum AhR only, n | — | 3 | — | — | — | — |
| CSF AhR only, n | — | 16 | — | — | — | — |
| Neither valid serum nor CSF AhR result, n | — | 1 | — | — | — | — |
| **Biomarkers and clinical measures** |  |  |  |  |  |  |
| Serum AhR-activating bioactivity, median [IQR] | 0.978 [0.835–1.197] (n=28) | 1.281 [1.148–1.554] (n=125) | — | — | — | — |
| CSF AhR-activating bioactivity, median [IQR] | — | 1.071 [0.900–1.283] (n=138) | — | — | — | — |
| CSF Aβ42, pg/mL, median [IQR] | — | 466 [341–616] (n=93) | — | — | — | — |
| CSF Aβ40, pg/mL, median [IQR] | — | 10,304 [8,404–12,476] (n=69) | — | — | — | — |
| CSF p-tau181, pg/mL, median [IQR] | — | 90.3 [59.0–127.2] (n=93) | — | — | — | — |
| CSF Aβ42/Aβ40, median [IQR] | — | 0.044 [0.040–0.053] (n=69) | — | — | — | — |
| Amyloid-positive (A+), n/available (%) | — | 67/93 (72.0%) | — | — | — | — |
| Tau-positive (T+), n/available (%) | — | 72/93 (77.4%) | — | — | — | — |
| MMSE, median [IQR] | — | 23 [21–27] (n=42) | — | — | — | — |
| GDS, median [IQR] | — | 3 [3–4] (n=91) | — | — | — | — |

Among participants in the COGNIBIO12 analytical subcohort, 82 of 139 with recorded sex were women (59.0%), and mean age was 74.8 ± 7.1 years (n=137). A clinical Alzheimer’s disease code was recorded in 85/142 participants (59.9%), while a suspected amyloid aetiology was recorded in a separate clinical variable for 87/142 (61.3%); these fields overlapped in 80 participants. Participant-level matching identified 122 paired serum-CSF AhR results, three serum-only and 16 CSF-only participants; one participant had neither valid AhR measurement.

Serum AhR-activating bioactivity was significantly higher in the COGNIBIO12 analytical subcohort than in cognitively unimpaired H2H controls. After adjustment for age and sex, COGNIBIO12 participants showed a 33.0% increase in serum AhR bioactivity (adjusted geometric mean ratio=1.330, 95% CI 1.162-1.523 p<0.0001) (Figure 6E; Supplementary Table 4-5). We next examined associations with amyloid (A) and tau (T) biomarker status within COGNIBIO12, separately from the external H2H comparison. The overall difference across the four AT groups was significant (p=0.0181), but this pattern was not explained by amyloid positivity alone: no significant difference was detected between A+ and A− participants or between A+T+ and A−T− participants. Instead, A−T+ participants showed lower serum AhR bioactivity than A−T− participants (adjusted geometric mean ratio=0.830, 95% CI 0.714-0.965, Holm-adjusted p=0.0470; Figure 6F and Supplementary Table 4-5). In the pooled analysis of the 87 COGNIBIO12 participants with complete AT classification and serum AhR results, T+ participants had lower serum AhR bioactivity than T− participants (age- and sex-adjusted geometric mean ratio=0.868, 95% CI 0.788–0.956; unadjusted p=0.0041), whereas the A+ versus A− comparison was not significant (ratio=1.021, p=0.6601).

In additional comparisons across these independently recruited cohorts, serum AhR bioactivity was 29.6% higher in A+ COGNIBIO12 participants than in cognitively unimpaired H2H participants (adjusted geometric mean ratio=1.295, 95% CI 1.130-1.485, p=0.0002) and 27.5% higher in A+T+ participants than in H2H participants (adjusted geometric mean ratio=1.276, 95% CI 1.117-1.457, p=0.0003) (Figure 6F and Supplementary Table 5). By contrast, within COGNIBIO12, serum AhR bioactivity did not differ significantly between A+ and A− participants (adjusted geometric mean ratio=1.021, p=0.6601). Thus, serum AhR bioactivity was higher in the memory-clinic cohort than in external controls, without a significant association with binary amyloid status within the clinical cohort.

Given the sex-stratified findings in mice, we next examined the human data by sex. Relative to sex-matched H2H controls, serum AhR bioactivity was 40.9% higher in A+ women (adjusted ratio=1.409, 95% CI 1.205-1.647, p<0.0001), whereas the corresponding difference in A+ men was smaller and not significant (adjusted ratio=1.115, p=0.4050) (Figure 6G and Supplementary Table 4-5). The formal group-by-sex interaction was not significant (p=0.1298; Figure 6G). No significant differences in CSF AhR-activating bioactivity were detected across AT groups (Figure 6H and Supplementary Tables 4-5). A+T+ participants showed a positive but non-significant estimate relative to A−T− participants, and neither amyloid nor tau positivity alone was associated with altered CSF AhR bioactivity. Serum-CSF associations were evaluated only in participant-matched specimens. Across all participants with paired serum and CSF AhR measurements (n=122), serum and CSF AhR bioactivities were not significantly correlated (Pearson r=0.091, p=0.3170; Spearman ρ=0.108, p=0.2359; Supplementary Table 5). We next examined these associations among amyloid-positive participants. Serum and CSF AhR bioactivities were positively correlated in A+ participants (Spearman ρ=0.378, p=0.0029, n=60). In the matched A+T+ subset (n=51), the Spearman correlation was ρ=0.465 (p=0.0006); the adjusted association remained significant (age- and sex-adjusted geometric mean ratio=1.077, 95% CI 1.028–1.128, p=0.0019; FDR q=0.0241; Figure 6I). The interaction between A+T+ status and the serum-CSF relationship was not significant (p=0.2603). We next examined the relationship between serum AhR-activating bioactivity and CSF Aβ42 concentrations among amyloid-positive participants. Serum AhR bioactivity showed an inverse correlation with CSF Aβ42 (Pearson r=−0.253, p=0.0496, n=61; Figure 6J), consistent in direction with our experimental findings linking AhR signalling to amyloid pathology. The association remained inverse in Spearman and age- and sex-adjusted analyses, although these tests did not reach statistical significance (p=0.1216 and p=0.0849, respectively). When amyloid positivity was defined using the CSF Aβ42/Aβ40 ratio, a similar inverse correlation was observed (r=−0.260, p=0.0529, n=56), whereas the association was attenuated when restricted to Aβ42-defined A+ participants with available ratio measurements (r=−0.107, p=0.4774, n=46; Supplementary Table 4-5). No significant association was detected between CSF Aβ42 and CSF AhR-activating bioactivity (Figure 6K).

### Sex-dependent gut microbial remodelling is associated with altered AhR-active indole metabolite profiles

We next investigated the potential metabolic origin of AhR-activating ligands. Because gut microbial tryptophan metabolism generates multiple endogenous AhR ligands, we first examined AhR-activating bioactivity in faecal extracts. APP^NL-F^ extracts elicited substantially greater reporter activity than those from WT littermates (Figure 7A). In females, faecal AhR-activating bioactivity also showed a positive association estimate with cortical amyloid burden (Spearman ρ=0.7097, p=0.0741, n=7), which did not reach statistical significance (Figure 7B and Supplementary Figure 13). This observation suggests a potential link between the intestinal AhR-activating environment and amyloid pathology.

**Figure 7.**
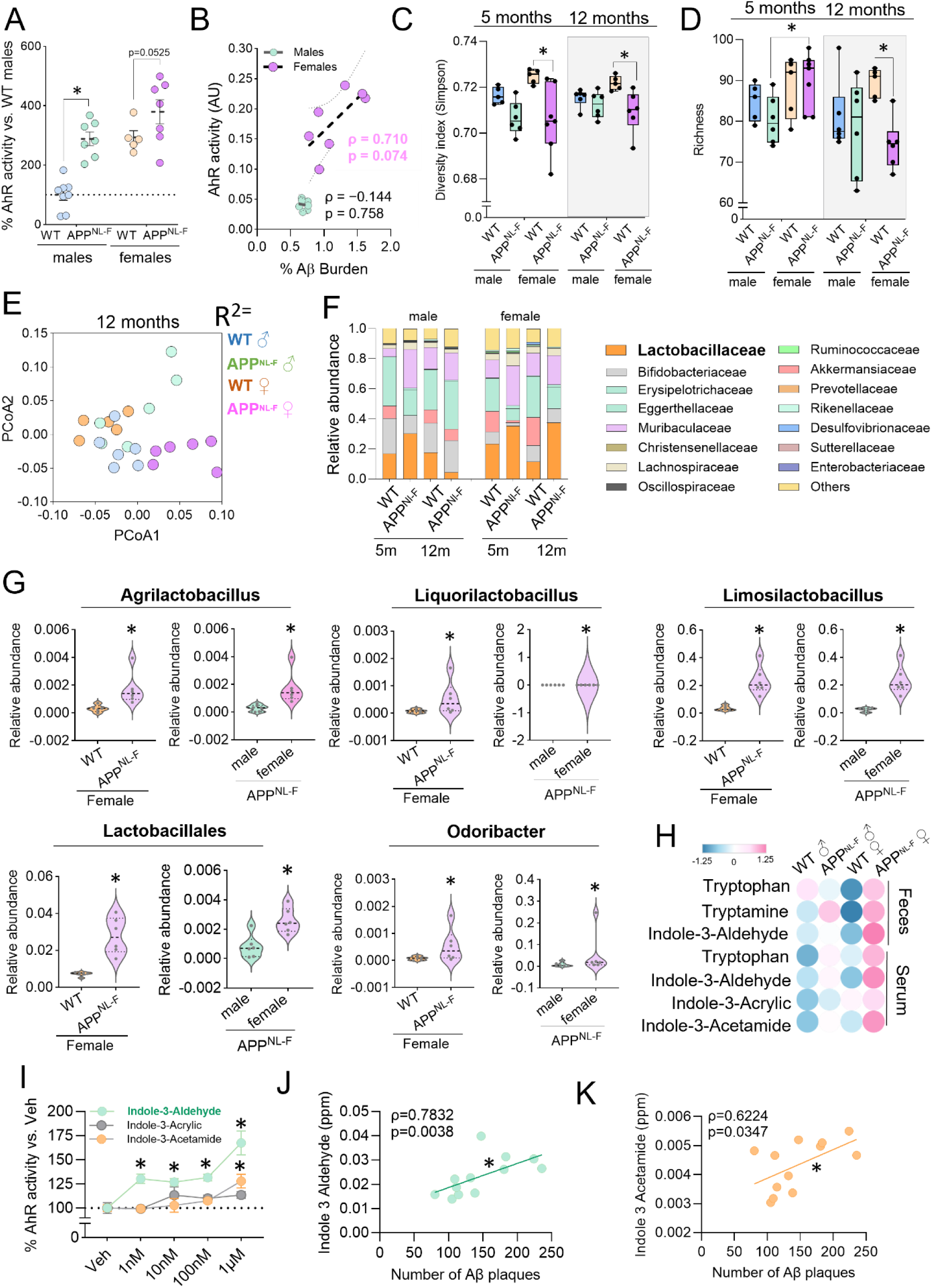
Sex-dependent remodelling of the gut microbiota and tryptophan metabolism is associated with an expanded pool of AhR-active indole metabolites in APP^NL-F^ mice. **(A)** AhR-activating bioactivity induced by faecal extracts from 12-month-old male and female WT and *APP*^NL-F^ mice, measured using an XRE-driven luciferase reporter assay. WT males: n=8; *APP*^NL-F^ males: n=7; WT females: n=5; *APP*^NL-F^ females: n=7. Two-way ANOVA followed by Fisher’s LSD test. WT versus *APP*^NL-F^: males, p<0.0001; females, p=0.0525. Males versus females: WT, p<0.0001; *APP*^NL-F^, p=0.0267. **(B)** Association between faecal AhR-activating bioactivity and cortical Aβ burden in 12-month-old male and female APP^NL-F^ mice. Correlations were assessed separately by sex using Spearman rank correlation: females, ρ=0.7097, p=0.0741, n=7; males, ρ=−0.1439, p=0.7583, n=7. Lines represent descriptive linear regression fits. **(C)** Simpson diversity index of the faecal microbiota in male and female WT and *APP*^NL-F^ mice at 5 and 12 months of age. At 5 months, WT males: n=5; WT females: n=5; *APP*^NL-F^ males: n=6; *APP*^NL-F^ females: n=7. At 12 months, WT males: n=6; WT females: n=5; *APP*^NL-F^ males: n=6; *APP*^NL-F^ females: n=6. Two-way ANOVA followed by Tukey’s multiple-comparisons test. At 5 months, WT females versus *APP*^NL-F^ males, p=0.0025, and WT females versus *APP*^NL-F^ females, p=0.0010. At 12 months, WT females versus *APP*^NL-F^ females, p=0.0582. **(D)** Bacterial richness in the faecal microbiota of male and female WT and *APP*^NL-F^ mice at 5 and 12 months of age. Group sizes were as indicated in **(C)**. Two-way ANOVA followed by Tukey’s multiple-comparisons test. At 12 months, bacterial richness was significantly lower in *APP*^NL-F^ females than in age-matched WT females, p=0.0090. All other comparisons were not significant. **(E)** Principal coordinates analysis based on Bray-Curtis dissimilarities, showing the distribution of faecal microbial communities according to genotype, sex and age. Each point represents one biologically independent mouse, and the distance between points reflects the degree of dissimilarity between microbial communities. Overall PERMANOVA: R²=0.4534, pseudo-F=5.2523, p=0.0010. Pairwise PERMANOVA with false-discovery-rate correction: APP^NL-F^ versus WT females, q=0.0080; APP^NL-F^ versus WT males, q=0.0110; female versus male APP^NL-F^ mice, q=0.0080; and female versus male WT mice, q=0.0080. **(F)** Relative abundance of Lactobacillaceae in the faecal microbiota of male and female WT and *APP*^NL-F^ mice at 5 and 12 months of age. Group sizes were as indicated in **(C)**. At 12 months, Lactobacillaceae abundance was higher in *APP*^NL-F^ females than in age-matched WT females and *APP*^NL-F^ males. Tukey’s multiple-comparisons test: WT females versus *APP*^NL-F^ females, p<0.0001; *APP*^NL-F^ males versus *APP*^NL-F^ females, p<0.0001. **(G)** Relative abundance of selected bacterial taxa associated with microbial tryptophan metabolism, including *Agrilactobacillus, Liquorilactobacillus, Limosilactobacillus, Odoribacter* and unclassified Lactobacillales, in 12-month-old mice. Left, WT females compared with *APP*^NL-F^ females; right, male compared with female *APP*^NL-F^ mice. WT females: n=5; *APP*^NL-F^ males: n=6; *APP*^NL-F^ females: n=6. Two-tailed Mann-Whitney U tests, p<0.05. **(H)** Integrated heatmap showing selected tryptophan-related metabolites in faecal extracts (upper panel) and serum (lower panel) from 12-month-old male and female WT and APP^NL-F^ mice, analysed using complementary targeted and untargeted metabolomic approaches. Metabolite values were independently standardized within each biological matrix using Z-scores calculated across all 12-month-old samples. Coloured circles represent the mean Z-score for each sex–genotype group, with blue and pink indicating lower and higher relative metabolite abundance, respectively. Five independent biological pools of faecal material were prepared per group. The CEMBIO dataset included five analysed pools per group, except for APP^NL-F^ mice females (n=4). Serum samples were obtained from individual mice (WT males, n=6; APP^NL-F^ mice males, n=7; WT females, n=5; APP^NL-F^ mice females, n=6). Pairwise comparisons were performed using two-sided Mann–Whitney U tests, with p values adjusted for multiple comparisons using the Benjamini–Hochberg false-discovery-rate (FDR) procedure. Adjusted q values <0.05 were considered statistically significant. In faecal samples, tryptophan, tryptamine and indole-3-aldehyde (IAld) were significantly increased in APP^NL-F^ females compared with WT females (q=0.0278 for each metabolite). Faecal tryptamine was also increased in APP^NL-F^ mice males compared with WT males (q=0.0466). In serum, IAld and indole-3-acetamide (I3AM) were significantly higher in APP^NL-F^ females than in APP^NL-F^ males (q=0.0047 for both). Additional significant sex differences were detected within WT mice for faecal tryptophan and tryptamine (q=0.0190 for both) and serum tryptophan (q=0.0379). Heatmap colours represent standardized metabolite abundance. **(I)** AhR-activating capacity of selected indole derivatives-indole-3-aldehyde (IAld), indole-3-acrylic acid (I3Acr) and indole-3-acetamide (I3AM)-assessed using an XRE-driven luciferase reporter assay at concentrations ranging from 1 nM to 1 µM. Reporter activity was normalized to the corresponding vehicle-treated condition, set to 100%. Vehicle: n=6; 1 and 10 nM: n=3; 100 nM and 1 µM: n=6. Comparisons against vehicle were performed using Dunnett’s multiple-comparisons test. IAld significantly increased AhR reporter activity at 1 nM, p=0.0079; 10 nM, p=0.0214; 100 nM, p=0.0005; and 1 µM, p<0.0001. I3Acr increased reporter activity at 1 µM, p=0.0023, whereas no significant differences were detected for I3AM. **(J)** Association between serum indole-3-aldehyde levels and cortical Aβ plaque number in 12-month-old *APP*^NL-F^ mice. Spearman correlation, ρ=0.7832, p=0.0038, n=12. **(K)** Association between serum indole-3-acetamide levels and cortical Aβ plaque number in 12-month-old *APP*^NL-F^ mice. Spearman correlation, ρ=0.6224, p=0.0347, n=12 Data are presented as mean±SEM unless otherwise indicated. Each dot represents one biologically independent mouse, except in the luciferase concentration-response experiments, in which n represents independent determinations. Heatmap colours represent standardized metabolite levels for visualization purposes only; Z-scores were calculated independently within each biological matrix and do not represent statistical integration or directly comparable absolute metabolite concentrations. Asterisks indicate p<0.05.

We therefore profiled the GM of male and female APP^NL-F^ and WT mice at 5 and 12 months of age. Bacterial richness was significantly reduced in 12-month-old APP^NL-F^ females compared with age-matched WT females (p=0.0090; Figure 7D). Rarefaction curves are shown in Supplementary Figure 14. The Simpson diversity index was also lower in APP^NL-F^ females than in WT females at 5 months (p=0.0010), whereas the corresponding difference at 12 months did not reach statistical significance (p=0.0582; Figure 7C). Bray-Curtis analysis further revealed differences in microbial community composition among genotype-sex groups (Figure 7E and Supplementary Figure 15).

We next investigated whether these changes involved bacterial taxa associated with tryptophan metabolism and the generation of potential AhR ligands. Lactobacillaceae emerged as a prominently altered family, with higher relative abundance in 12-month-old APP^NL-F^ females than in WT females and APP^NL-F^ males (Figure 7F), revealing a sex-dependent divergence in a bacterial family closely linked to microbial tryptophan metabolism. At finer taxonomic resolution, female APP^NL-F^ mice showed increased abundance of *Agrilactobacillus, Liquorilactobacillus* and *Limosilactobacillus*, together with unclassified Lactobacillales and *Odoribacter* (Figure 7G). Additional changes involved *Ligilactobacillus, Lactobacillus, Alistipes* and *Bacteroides* (Supplementary Figure 16). These taxa were selected because they have been associated with microbial tryptophan metabolism and the generation or transformation of indole-related compounds with the potential to modulate AhR signaling^12, 30, 31, 32, 33^. Consistent with this taxonomic shift, PICRUSt2-based functional inference revealed remodelling of microbial aromatic amino-acid metabolism, including increased predicted alcohol and aryl-alcohol dehydrogenase functions in female APPNL-F mice (Supplementary Table 6). Thus, the microbiota changes observed in APP^NL-F^ mice were not random taxonomic fluctuations but converged on functions directly relevant to the generation of AhR-active metabolites.

Because functional inference cannot establish actual metabolite production, we next examined selected tryptophan-related metabolites in faecal and serum samples using complementary targeted and untargeted metabolomic approaches (Figure 7H). For visualization, metabolite values were standardized separately within each biological matrix and represented as group-mean Z-scores. At 12 months, faecal tryptophan, tryptamine and indole-3-aldehyde (IAld) showed higher values in female APP^NL-F^ preparations than in female WT preparations (FDR q=0.0278 for each metabolite). Faecal tryptamine was also higher in male APP^NL-F^ than in male WT preparations (q=0.0466). In serum, IAld and indole-3-acetamide (I3AM) were significantly higher in female than in male APP^NL-F^ mice (q=0.0047 for both), whereas indole-3-acrylic acid (I3Acr) did not show significant differences in the assessed comparisons. Thus, the sex-dependent microbial remodelling was accompanied by a corresponding shift in the pool of indole derivatives reaching the circulation.

We next asked whether these circulating indoles differed in their ability to activate AhR. IAld produced a robust concentration-dependent increase in AhR reporter activity, whereas I3AM elicited a weaker response and 3Acr showed no AhR activation across most concentrations tested (Figure 7I). We then examined whether circulating levels of these metabolites were associated with amyloid pathology. Serum IAld showed a strong positive correlation with cortical Aβ plaque number (ρ=0.7832, p=0.0038) (Figure 7J), while I3AM was also positively associated with plaque number (r=0.6224, p=0.0347) (Figure 7K). Additional analyses showed corresponding associations with cortical amyloid burden (Supplementary Figure 16). By contrast, I3Acr was not significantly associated with amyloid pathology. Notably, IAld emerged as the metabolite that most directly connected these observations, combining AhR-activating capacity with an association between its measured serum levels and cortical amyloid pathology. Together, these findings link sex-dependent remodelling of the gut microbial-tryptophan metabolic network to an altered circulating pool of AhR-relevant indole metabolites associated with amyloid accumulation.

## Discussion

Our findings identify AhR as a ligand-responsive immunometabolic checkpoint that shapes microglial functional state in response to amyloid pathology. Rather than acting simply as a rheostat of microglial activation, AhR influenced transcriptional programs used by microglia to recognize, internalize and process Aβ. Ligand-induced receptor activation favoured amyloid accumulation, whereas AhR loss shifted microglia toward phagolysosomal and Aβ-recognition states and reduced plaque pathology. Importantly, inducible deletion of AhR in CX3CR1-expressing microglia using the Cx3cr1-CreER system significantly reduced cortical amyloid burden *in vivo*, supporting a key contribution of microglial AhR to amyloid accumulation. These effects were accompanied by marked sex-associated differences in microglial transcriptional responses and were further modulated by the systemic and tissue AhR-activating environment, associated with changes in gut microbial composition and tryptophan-derived metabolites. Together, these findings position AhR as a molecular interface through which systemic metabolic signals are translated into sex-dependent microglial responses to pathological protein accumulation. This sex dependence was evident at multiple levels, including the markedly stronger microglial transcriptional and functional response to AhR loss in females and the sex-dependent pattern of AhR-activating bioactivity in peripheral and central compartments. The mechanisms underlying these differences remain unresolved and may involve gonadal hormones, sex chromosomes, age-dependent endocrine changes, basal AhR pathway activity, or sex-dependent variation in the microbiota and circulating ligand environment.

The effects of AhR were closely linked to the microglial response to plaques. AhR expression increased with amyloid progression and was enriched in plaque-associated glia, particularly microglia. AhR-deficient mice showed an increased ThioS-to-Aβ ratio and greater microglial coverage of amyloid deposits, features consistent with enhanced plaque compaction and containment. This is functionally relevant because plaque-associated microglia can form a barrier that limits the spread of potentially toxic Aβ species, promotes plaque compaction and reduces surrounding neuritic damage^1, 26, 34^.

A central implication of our study is that AhR regulates functional specialization within the microglial compartment. AhR deletion induced a coordinated program of Aβ recognition, uptake, and lysosomal processing, accompanied by increased microglial Aβ content and uptake of Methoxy-X04-labelled fibrillar amyloid. Notably, AhR loss did not simply expand canonical DAM-like microglia^2^. Instead, it redistributed the microglial compartment from MgC0 toward the closely related MgC1 state, which was enriched in Fc-receptor signalling, endocytosis, phagocytosis, and lysosomal processing. This suggests that AhR regulates functional specialization within the plaque-associated microglial compartment rather than acting as a general switch for microglial activation. This is consistent with the increasingly heterogeneous view of plaque-associated microglia emerging from mouse and human single-cell studies^35, 36^. Microglial receptors such as TREM2 and the TAM receptors Axl and Mer participate in plaque recognition, organization, and engulfment^29^. AhR is conceptually different. As a ligand-responsive transcriptional sensor, it is unlikely to function primarily as a receptor that directly recognize amyloid or phosphatidylserine-associated signals: instead, our data support a model in which, AhR may instead determine the cellular program through which those recognition and engulfment signals generated in the plaque environment are processed. In this context, metabolic sensing becomes an additional layer of control over the effector state of a tissue-resident macrophage. Importantly, inducible deletion of AhR in microglia reduced plaque burden at both disease stages examined, directly implicating microglial AhR in amyloid accumulation *in vivo*.

Biological sex emerged as an important modifier of this AhR-dependent response. Female APP^NL-F^ mice developed a greater amyloid plaque burden than males and, although AhR deletion reduced pathology in both sexes, its cellular and transcriptional consequences were markedly stronger in females. Female microglia showed broader induction of pathways related to Aβ recognition, phagocytosis, and lysosomal processing, together with a greater increase in Aβ uptake, whereas male microglia retained a comparatively stronger proteostasis-, UPR- and mTOR-related signature.

These observations suggest that sex affects not simply the amplitude but also the nature of the microglial transcriptional response when AhR signalling is altered. This is consistent with evidence that sex shapes molecular and microglial responses in AD and aging models, and that female microglia can show earlier or more extensive transitions toward plaque-associated states^7, 36, 37^. Moreover, microbiota manipulation can modify amyloid pathology in a sex-dependent manner; notably, *Bacteroides fragilis* administration increased amyloid plaques selectively in female APP/PS1 mice while suppressing microglial programs related to phagocytosis and protein degradation^12^. Our findings extend this concept by identifying a metabolite-sensing transcription factor as a potential molecular node at which sex and environmental signals converge. Together, these observations suggest that the greater amyloid vulnerability of females is accompanied by a distinct sensitivity of microglia to metabolic signals that engage AhR.

The ligand-responsive nature of AhR also raises the possibility that differences in extracellular ligand availability contribute to the sex-associated patterns observed. In parallel with the microglial phenotypes, APP^NL-F^ mice showed sex-associated differences in intestinal microbial composition and tryptophan-related metabolites, including differences in serum IAld and I3AM. Certain gut bacteria can produce AhR-active indoles^13, 14^, but our experiments did not establish the microbial origin of the measured circulating metabolites or demonstrate that faecal changes caused the serum differences. IAld robustly activated the AhR reporter *in vitro*, and its serum abundance correlated with cortical amyloid pathology. Together, these findings suggest a model in which biological sex may influence both the response of microglia to AhR pathway perturbation and the extracellular ligand environment to which the receptor is exposed. These levels may interact; however, the present experiments do not establish that sex-associated microglial differences are cell-autonomous or independent of the microbiota.

Gut microbial metabolism represents a plausible source of the tryptophan-derived indoles detected in our study, particularly IAld, whose production by intestinal bacteria is well established. However, circulating indole profiles may also be influenced by dietary sources, metabolism by other microorganisms, and host-dependent metabolic processes^38^. Thus, although the parallel changes in gut microbial composition and indole metabolites support a potential microbiota-AhR connection, our data do not establish the intestinal origin of the circulating metabolites. Importantly, AhR activation by microbiota-derived indoles is not uniformly detrimental in experimental AD. Several tryptophan-derived indoles, including indole, IAA and indole-3-propionic acid have been reported to attenuate neuroinflammation and amyloid-related pathology in APP/PS1 mice through AhR-associated mechanisms^22^, and microbiome-derived ILA reduced amyloid accumulation in 5xFAD mice AhR-dependently involving microglia and astrocytes^23^. Bifidobacteria can also increase circulating and brain ILA and reduce neuroinflammatory responses in other disease contexts^39^. More recently, IAA was shown to reduce microglia-mediated synaptic loss in AD models through an AhR-CCR4 pathway^24^. Together with the detrimental effect of sustained I3S-mediated AhR activation observed here, these studies argue against classifying AhR signalling as intrinsically protective or pathogenic.

Different AhR ligands vary in affinity, pharmacokinetics, tissue distribution and duration of receptor engagement, and the transcriptional output of AhR is strongly dependent on cellular and disease context, and biological sex. AhR may therefore be better understood as a context decoder than as a unidirectional inflammatory switch. AhR signalling was recently shown to limit microglial engulfment of synaptic material^40^. Therefore, the biological consequence of AhR engagement in microglia may depend on the ligand encountered, the duration of signalling, the pathological substrate being processed and the underlying cellular state. This framework could reconcile apparently opposing effects of AhR-active metabolites reported across experimental systems.

Our human data provided a translational extension of the experimental findings, revealing differences in circulating AhR-activating bioactivity in individuals evaluated for cognitive impairment. Serum AhR bioactivity was approximately 33% higher in the clinically heterogeneous COGNIBIO12 memory-clinic cohort than in cognitively unimpaired participants from the independently recruited H2H cohort. Although differences between cohorts may contribute to this observation, these findings indicate variation in the circulating environment capable of activating AhR. Importantly, serum AhR bioactivity was not determined by binary amyloid status within COGNIBIO12, suggesting that this functional readout may capture aspects of systemic pathway engagement beyond conventional amyloid classification. Unlike measurements of individual circulating metabolites, the XRE reporter integrates the AhR-activating capacity of multiple circulating factors^41, 42^. Within amyloid-positive participants, serum AhR-activating bioactivity showed an inverse association with CSF Aβ42 concentrations, providing a human observation consistent in direction with our experimental findings linking AhR signalling to amyloid pathology. Although this association did not remain statistically significant after adjustment for age and sex or correction for multiple comparisons, it raises the possibility that circulating AhR-activating factors are related to amyloid-associated processes in humans. Exploratory sensitivity analyses indicated that the association was influenced by sample selection and the definition of amyloid positivity, supporting the need for further investigation in clinically well-characterized cohorts. The absence of a difference in serum AhR bioactivity between A+ and A− participants does not preclude an association within the A+ group, as these analyses address different features of the data: binary amyloid status reflects whether CSF Aβ42 crosses a diagnostic threshold, whereas the continuous analysis captures variation in Aβ42 concentrations within the amyloid-positive range. Importantly, serum and CSF AhR bioactivities were positively associated in A+T+ participants, and this relationship remained significant after adjustment for age and sex and correction for multiple comparisons. Although formal interaction analysis did not establish specificity to the A+T+ biomarker profile, these findings support a relationship between peripheral and central AhR-activating environments. Together, these observations suggest that circulating AhR bioactivity may reflect a complex state of pathway engagement whose relationship with amyloid pathology depends on disease stage, ligand composition and cellular context^43, 44^.

An important implication of these findings is the substantial interindividual variation in circulating AhR bioactivity among participants with the same conventional AT biomarker classification. This suggests that AhR pathway engagement may capture a dimension of disease biology not encoded by amyloid and tau status alone. Circulating AhR bioactivity could therefore provide an accessible functional readout for biological stratification according to the degree of engagement of this immunometabolic pathway and, in future intervention studies, for monitoring changes in the systemic ligand environment or AhR-dependent programs.

Biological sex may represent an important modifier of this state. Serum AhR bioactivity was markedly higher in amyloid-positive women than in sex-matched cognitively unimpaired controls, whereas the corresponding difference in men was smaller and non-significant. Together with the stronger transcriptional and functional responses observed in female microglia following AhR loss and the sex-dependent remodelling of the AhR-active metabolic environment in APP^NL-F^ mice, the larger difference observed in amyloid-positive women raises the possibility that biological sex also influences circulating AhR pathway engagement in humans. These findings raise the possibility that sex influences both the cellular response to AhR signalling and the degree of systemic pathway engagement, potentially contributing to interindividual differences in vulnerability or response to pathway-directed interventions. The formal group-by-sex interaction was not significant, however, and the human and experimental readouts are not directly equivalent; prospective studies in larger cohorts will therefore be required to establish the extent to which sex contributes to AhR-based biological stratification. If validated prospectively, such pathway-based stratification could help identify individuals in whom AhR-dependent immune programs are particularly engaged and could support both the selection and pharmacodynamic monitoring of interventions aimed at modifying the ligand environment or downstream microglial responses.

Several considerations should frame the interpretation of our findings. The XRE-reporter assay measures integrated AhR-activating capacity but does not identify the individual ligands responsible for this activity^41^. Although IAld is a well-established microbiota-derived tryptophan metabolite^30^, our data do not directly trace the microbial origin of circulating IAld or establish which microbial taxa accounts for its increase. Constitutive AhR deficiency may involve additional peripheral effects^45, 46^, although inducible AhR deletion in microglia directly supports a microglial contribution to amyloid accumulation. The APP^NL-F^ model predominantly captures amyloid pathology; whether these mechanisms extend to tau-driven neurodegeneration remains to be established^25^. In the human cohort, CSF Aβ40 measurements were initially prioritized for patients with clinical suspicion of Alzheimer’s disease because of limited assay-kit availability. Consequently, Aβ42/Aβ40 measurements were available in a clinically selected subset, limiting their use for primary AT classification and potentially introducing selection bias into the corresponding sensitivity analyses. Differences between the independently recruited COGNIBIO12 and H2H cohorts, beyond those accounted for by age and sex, may also have influenced the external cohort comparisons. Despite these limitations, our findings identify AhR as a key regulator of microglial responses to amyloid pathology.

This mechanistic framework also opens a potential therapeutic avenue. Given the opposing effects reported for different AhR ligands and the dependence of AhR responses on cellular context and biological sex, future studies should determine whether the microglial programs identified here can be selectively manipulated to enhance Aβ recognition, lysosomal processing and plaque containment while preserving physiological AhR functions in other cells and tissues. A complementary strategy may be to define and modify the disease-associated ligand environment that engages these responses. From a translational perspective, circulating AhR-activating bioactivity could potentially provide a functional measure of pathway engagement to support patient stratification or pharmacodynamic monitoring in such studies. Identifying the relevant ligands, downstream effectors and biological contexts in which these programs operate will be essential before the pathway can be considered therapeutically actionable.

In summary, our study identifies microglial AhR as a metabolically responsive regulator of amyloid pathology. Our findings support a model in which the systemic ligand environment determines AhR pathway engagement, biological sex shapes the resulting microglial response, and this interaction influences the capacity of microglia to recognize, process and contain amyloid pathology. The detection of altered circulating AhR bioactivity in human neurodegenerative disease further suggests that this immunometabolic pathway is not restricted to experimental models and provides a foundation for investigating AhR pathway engagement as a biologically informative and potentially actionable dimension of human disease.

## Materials And Methods

### Animals

Male and female APP^NL-F^ knock-in mice were used as the amyloid model throughout the study. Animals were analysed at different ages between 5 and 16 months, depending on the experimental design. The age, sex and genotype distribution for each experiment is specified in the corresponding figure legends. APP^NL-F^ knock-in mice were kindly provided by Dr. Takaomi C. Saito (RIKEN Centre for Brain Science, Saitama, Japan) and maintained as homozygous breeding colonies. These mice carry the Swedish (*NL*) and Beyreuther/Iberian (*F*) mutations in the endogenous murine *App* locus. The Swedish mutation increases the production of Aβ40 and Aβ42, whereas the Beyreuther/Iberian mutation increases the Aβ42/Aβ40 ratio^25^. To investigate the effect of constitutive AhR loss, APP^NL-F^ mice were crossed with AhR-deficient mice kindly provided by Dr. Pedro M. Fernández-Salguero (Universidad de Extremadura, Badajoz, Spain). AhR-deficient mice were originally generated by homologous recombination through disruption of exon 1 of the *Ahr* gene^47^. The resulting APP^NL-F^/AhR^−/−^ mice were maintained on a C57BL/6 genetic background. Depending on the experiment, APP^NL-F^/AhR^+/+^, APP^NL-F^/AhR^+/−^ and APP^NL-F^/AhR^−/−^ littermates were compared. To investigate the contribution of microglial AhR, homozygous APP^NL-F^ mice were crossed with *Ahr*^fl/fl^ mice^48^ (Jackson Laboratory Ahr^tm3.1Bra/J^ stock no. 006203) and subsequently with heterozygous CX3CR1^CreER^ mice^49,50^ (Jackson Laboratory B6.129P2(Cg)-Cx3cr1^tm2.1(cre/ERT2)Litt/WganJ^ stock no. 021160). Tamoxifen administration induced Cre-mediated recombination of the floxed *Ahr* allele in *Cx3cr1*-expressing cells, leading to sustained AhR depletion in long-lived microglia. Importantly, although this approach preferentially targets parenchymal microglia after an appropriate chase period, recombination may also occur in other CX3CR1-expressing myeloid populations, including CNS border-associated macrophages, particularly perivascular and meningeal macrophages^51, 52^. Tamoxifen-treated mice are hereafter referred to as AhR^KO-Mic^ mice, whereas the corresponding control animals are referred to as AhR^fl/fl^ controls. The tamoxifen administration protocol is described below.

Animals were housed under controlled temperature and humidity conditions on a 12h light/12h dark cycle, with food and water available *ad libitum*. Experimental colonies were maintained at the Universidad Complutense de Madrid (UCM) animal facility until April 2021 and subsequently transferred to the Centro Nacional de Investigaciones Cardiovasculares (CNIC) animal facility.

All experimental procedures were conducted in accordance with Spanish legislation governing the protection of animals used for scientific purposes (Royal Decree 53/2013) and with European Directives 86/609/EEC and 2010/63/EU. Experimental protocols were reviewed and approved by the Animal Welfare and Ethics Committees of the UCM and CNIC and authorized by the Consejería de Medio Ambiente y Ordenación del Territorio de la Comunidad de Madrid under authorization PROEX 047/16, PROEX 193.1/21 and PROEX 166.7/24. Animal experiments were designed and reported in accordance with the ARRIVE guidelines (Animal Research: Reporting of In Vivo Experiments). The study also adhered to the Transparency and Openness Promotion Guidelines. The data supporting the findings of this study are available from the corresponding authors upon reasonable request.

### In vivo treatments

#### Indoxyl-3-sulfate administration

To assess the effect of sustained AhR activation on amyloid pathology, 10-month-old *APP*^NL-F^ mice were treated with indoxyl-3-sulfate (I3S) for 6 weeks. The administration regimen was adapted from a previously described protocol using systemic I3S treatment in a model of central nervous system inflammation^18^. I3S was dissolved in dimethyl sulfoxide (DMSO) to prepare a concentrated stock solution (25 µg/µL), aliquoted and stored at −20°C. Immediately before administration, the stock solution was diluted 1:10 in sterile 0.9% saline to obtain a final concentration of 2.5 µg/µL. I3S was administered intraperitoneally at 10 µg/g body weight, equivalent to 10 mg/kg, every other day for 6 weeks. Control animals received an equivalent volume of the corresponding vehicle.

#### Methoxy-X04 (MeX04) assay

Methoxy-X04 (MeX04; Tocris Bioscience, catalog no. 4920) was initially dissolved in DMSO at 10 mg/mL. A 2 mg/mL working solution was prepared by mixing two volumes of the MeX04 stock solution with three volumes of DMSO and five volumes of 0.9% saline adjusted to pH 12.0. Mice received a single intraperitoneal injection of MeX04 at 10 mg/kg, 24 h before euthanasia. MeX04-labeled fibrillar Aβ deposits were subsequently assessed by histological analysis, whereas MeX04 uptake by cortical CD11b-positive microglia was quantified by flow cytometry as described below^53, 54^.

### Tamoxifen-induced microglial AhR deletion

To induce AhR deletion in microglia, 2-month-old APP^NL-F^/*Ahr*^fl/fl^/CX3CR1^CreER^ mice received tamoxifen by oral gavage. Tamoxifen was prepared at 5 mg/mL in corn oil containing 10% ethanol, and mice received 200 µL per administration, corresponding to 1 mg of tamoxifen per mouse. Three doses were administered on alternate days. Control mice received an equivalent volume of vehicle containing 10% ethanol in corn oil. Mice maintained until 12 months of age received an additional booster dose at 9 months. Tamoxifen-treated mice are hereafter referred to as AhR^KO-Mic^ mice, whereas vehicle-treated animals are referred to as AhR^flox^ controls.

### Behavioral testing

#### Open-field test

General locomotor activity and anxiety-like behaviour were assessed using the open-field test. Each mouse was placed in the centre of a square arena measuring 45 cm x 45 cm x 40 cm and allowed to explore freely for 7 min. Behaviour was recorded with an overhead camera and analysed using ANY-maze software. The total distance travelled, mean speed, immobility time, and time spent in the central and peripheral regions were quantified. The central region was defined as the inner 40% of the arena. The apparatus was cleaned with ethanol between animals to minimize olfactory cues.

#### Nest-building test

Nest-building behaviour was assessed as a measure of species-typical activity and daily living function. Mice were individually housed overnight in clean cages containing a pre-weighed amount of compressed cotton nesting material. The following morning, nest quality was scored using the five-point Deacon scale, in which a score of 1 indicates minimal manipulation of the nesting material and a score of 5 indicates a fully formed and organized nest. The amount of nesting material used was also determined by weighing the unshredded material remaining in the cage and expressing the result as the percentage of the initial material used for nest construction. Unused fragments weighing more than 0.1 g were collected and weighed^55^.

#### Spontaneous alternation in the Y-maze

Spatial working memory was assessed using the spontaneous alternation test in a Y-maze consisting of three identical solid-white arms arranged at 120° angles. Each arm measured 35 cm in length, 7 cm in width and had 20-cm-high walls. Each mouse was placed in the central junction and allowed to explore the maze freely for 5 min. An arm entry was recorded when the mouse entered an arm with all four paws. The sequence and total number of arm entries were recorded, and spontaneous alternation was defined as consecutive entries into all three arms without repetition, such as ABC, BCA or CAB. The percentage of spontaneous alternation was calculated as the number of observed alternations divided by the total number of arm entries minus two, multiplied by 100. Behaviour was recorded using an overhead camera and analysed by an investigator blinded to the experimental groups. The maze was cleaned with ethanol between animals to remove olfactory cues.

### Histology

#### Tissue preparation

Mice were perfused transcardially with phosphate buffer (0.1M). Left hemispheres of brains were post-fixed 24 h in 4% paraformaldehyde (PFA) and transferred to 30% sucrose. Coronal sections (30 μm) were cut using a microtome (Leica SM2000R) and stored in cryoprotective solution. Series of 1:10 coronal sections spaced 300 μm apart were used. The brains used for AhR immunofluorescence were coronal sectioned using a cryostat (Leica CM1950), obtaining 2 series of 10 µm and 8 series of 30 µm sections.

#### Immunofluorescence

Immunofluorescence was performed on free-floating sections that were incubated overnight at 4°C with the following antibodies: mouse anti-Amyloid β 1-16 (1:250, 6E10, BioLegend, 803015), rabbit anti-Iba1 (1:500, Wako, 011-27991), goat anti-AhR (1:100, Novus biologics, NB100-128SS), rat anti-CD68 (1:100, Abcam, ab53444), mouse anti-NeuN (1:100, Merck, MAB377), rabbit anti-GFAP (1:500, Agilent Technologies, Z033429-2), chicken anti-GFAP (1:500, Invitrogen, PA1-10004v), rat anti-LAMP1 (1:100, DSHB, 1D4B). The secondary antibodies used were goat anti-mouse 568 (1:500, Invitrogen, A-11031), goat anti-rabbit (1:500, Invitrogen, A-21245), donkey anti-goat biotinylated (1:300, Invitrogen, A-16003), goat anti-rat 488 (1:500, Invitrogen, A-11006), goat anti-mouse 405 (1:500, Invitrogen, A-31553), chicken anti-rabbit 647 (1:500, Invitrogen, A-21443), goat anti-chicken 488 (1:500, Invitrogen, A-11039), chicken anti-rat 647 (1:500, Invitrogen, A-21472), Streptavidin 568 (1:300, Thermo Fisher Scientific, S11226). In the immunofluorescence assays that included the anti-amyloid β antibody prior to incubation with the primary antibody, a permeabilization with 70% formic acid was performed for 20 minutes, followed by a 30-minute incubation with 1M glycine to unmask the Aβ oligomers.

#### Thioflavin-S staining

After immunofluorescence, those samples intended for amyloid plaque quantification were stained with Thioflavin-S. The mounted sections on SuperFrost (Fisher Scientific, 10149870) slides were sequentially incubated for 2 minutes in distilled water, 30 minutes in 1% Thioflavin-S (Sigma-Aldrich, T1892), 3 minutes in 50% alcohol, and 2 minutes in distilled water. Afterward, the sections were left to dry and coverslipped using Aqua-Poly/Mount (Polysciences, 18606-100).

### Image acquisition, processing, and analysis

Images were acquired using identical microscope settings within each experiment. Image processing and quantitative analyses were performed using ImageJ/Fiji (**ImageJ 1.54t)**. Anatomically matched brain sections were selected at equivalent rostrocaudal levels according to the Paxinos mouse brain atlas by Franklin and Paxinos (2008)^56^. Selected sections were located between bregma +2.34 mm and bregma −2.92 mm. Depending on the experiment, consecutive sections were collected at intervals of either 300 or 600 µm, and the number of sections analysed varied according to the specific quantification performed. Investigators performing image acquisition and quantification were blinded to the experimental groups. Intensity thresholds and image-analysis parameters were kept constant for all images included within the same experiment.

#### Amyloid plaque quantification

Aβ-positive and ThioS-positive amyloid plaques were quantified using two different imaging systems, depending on the experimental group. In a subset of experimental groups, plaques were quantified using a Zeiss Axiovert 200M microscope equipped with an ApoTome structured-illumination system at the CNIC Microscopy Unit. Eleven anatomically matched sections spanning the rostrocaudal axis were analysed per animal, with sections spaced approximately 300 µm apart. Plaque numbers were expressed either as the mean number of plaques per cortical section for each animal or as the cumulative number of plaques per hemisphere across the 11 analysed sections. In other experimental groups, images were acquired using a Leica DM6B upright microscope equipped with the Navigator imaging system. Six anatomically matched sections spanning the rostrocaudal axis, selected between bregma +2.34 mm and −2.92 mm, were analysed per animal. The number of Aβ-positive and ThioS-positive plaques was quantified using Fiji/ImageJ.

#### Cellular distribution of AhR

Confocal images were acquired using a Zeiss LSM 700 laser-scanning confocal microscope at the CNIC Microscopy Unit. Z-stacks were acquired using the same imaging parameters for all experimental groups within each analysis. To examine the cellular distribution of AhR in the cerebral cortex, its colocalization with NeuN-positive neurons, GFAP-positive astrocytes and Iba1-positive microglia was evaluated in confocal Z-stacks acquired using a 25× objective. Four serial 10-µm sections were analysed per animal.

#### Amyloid plaque size and classification

Amyloid plaque size and morphology were analysed in confocal Z-stacks acquired with a Zeiss LSM 700 confocal microscope using a 25X objective. Six cortical fields were imaged per animal across six anatomically matched sections, spaced 600 µm apart, with a section thickness of 30 µm. Between 20 and 30 plaques were analysed per animal. Plaques were automatically classified according to their Aβ and ThioS labelling using a custom-made Fiji/ImageJ macro. Dense-cored plaques were defined as Aβ-positive deposits containing a ThioS-positive fibrillar core, whereas diffuse plaques were defined as Aβ-positive deposits lacking a detectable ThioS-positive core, as previously described ^1, 26, 57^. Plaque compaction was defined as the degree of fibrillar packing of Aβ within a plaque, estimated as the proportion of dense-cored plaques relative to the total number of plaques and as the ThioS-positive core area relative to the total Aβ-positive plaque area (ThioS/Aβ area ratio); higher values indicate more compact plaques, whereas lower values indicate a more diffuse, less compacted morphology^1, 28, 58^.

#### Cortical Aβ burden and dystrophic neurites

The cortical area occupied by Aβ-positive deposits and LAMP1-positive dystrophic neurites was quantified from two tiled confocal Z-stack acquisitions per section, acquired with a Zeiss LSM 700 confocal microscope. Tile scans were acquired using 3×1 and 1× 3 arrangements with a 10X objective and 0.5 digital zoom to capture the largest possible cortical area. Four anatomically matched sections, spaced 600 µm apart, were analysed per animal. Cortical regions of interest were delineated, and fixed intensity thresholds were applied to the Aβ and LAMP1 channels; thresholds were kept constant across all animals and experimental groups within each experiment. Results were expressed as the percentage of the analysed cortical area occupied by Aβ immunoreactivity or LAMP1-positive dystrophic neurites.

#### Microglial morphology and plaque association

Microglial morphology and plaque-associated microglial responses were analysed in confocal Z-stacks acquired with a Zeiss LSM 700 confocal microscope from five serial 30-µm sections per animal, spaced 600 µm apart. Five cortical fields per section were analysed for morphological analysis, and five cortical fields per section were analysed for plaque-associated microglia. Images for morphological analysis were acquired using a 40X objective with 0.5 digital zoom, whereas plaque-associated microglia were imaged using a 40X objective. To quantify microglial coverage of amyloid plaques, maximum-intensity projections were generated and fixed thresholds were independently applied to the plaque and microglial channels. Plaques were identified using either Aβ immunoreactivity or ThioS labelling. Microglial coverage was calculated as the area of the plaque overlapping with Iba1 immunoreactivity, divided by the total plaque area and expressed as a percentage. Between 8 and 15 plaques were analysed per animal, depending on plaque availability within the acquired images.

#### Microglial Aβ uptake

To assess microglial Aβ uptake, five cortical fields were analysed per animal. The Aβ signal contained within Iba1-positive microglial profiles was quantified throughout the Z-stack using a custom-made Fiji/ImageJ macro that analysed Aβ uptake by microglia on a slice-by-slice (z-by-z) basis. Results were expressed as the percentage of total Aβ immunoreactivity contained within Iba1-positive microglia.

#### Non-plaque associated microglia

For morphological analysis of non-plaque-associated microglia, Iba1-positive cells located more than 50 µm away from amyloid deposits were randomly selected within the cerebral cortex. Approximately 8-12 cells were analysed per animal. Images were processed and skeletonized in Fiji using a custom-made macro adapted from Young and Morrison (2018)^59^, and morphological parameters, including the number of branches and endpoints per cell, were quantified following the previously described skeleton-analysis workflow.

#### Methoxy-X04-positive amyloid plaques

Methoxy-X04-positive amyloid plaques were analysed using a Leica DM6B upright microscope equipped with the Navigator imaging system at the CNIC Microscopy Unit. Images covering the entire brain section were acquired using a 5X objective. Six anatomically matched sections were quantified per animal, selected between bregma +2.34 mm and −2.18 mm. A fixed intensity threshold was applied to identify Methoxy-X04-positive deposits within the cerebral cortex; the same threshold was used for all animals and experimental groups. The cortical area occupied by Methoxy-X04 labelling and the number of Methoxy-X04-positive plaques were quantified using the Analyze Particles tool in Fiji, with a minimum particle size of 6 pixels, no upper size limit, and a circularity range of 0-1.

### SDS-PAGE and Western blot

For determination of total protein levels, samples were homogenized using a Teflon-glass grinder with lysis buffer: 50 mM Tris-HCl pH 7.5, 150 mM NaCl, 1 mM EDTA, and 0.1 % Igepal with Complete™ Protease Inhibitor Cocktail (Roche, 11873580001) and PhosSTOP™ (Roche, 4906845001). Equal amounts of total protein (30 μg) were resolved by SDS-PAGE and transferred to Immobilon-PVDF membranes (Millipore, IPVH00010). The membranes were blocked with 5% nonfat milk in TBS-T (1% Tween 20 in tris-buffered saline) and incubated with goat anti-AhR (1:250, Novus biologics, NB100-128SS). Rabbit β-actin (1:2000; Sigma-Aldrich, MABT523) was included to ensure equal protein loading. Immunoreactive bands were visualized using the corresponding horseradish peroxidase-conjugated IgG (1:2500, Thermo Scientific) and revealed by Clarity Western ECL substrate following manufacturer’s instructions (Bio-Rad). Blots were imaged using an iBright 1500 (Invitrogen) and quantified by densitometry (ImageJ). All densitometries are expressed in arbitrary units of optical density.

### Aβ40 and Aβ42 ELISAs

#### Protein extraction

To determine the levels of Aβ40 and Aβ42 in the cerebral cortex using ELISAs, a fractionation process was first performed to separate the soluble and insoluble fractions. Each cortex sample was homogenized in 500 µL of TBS protein extraction buffer (50 mM Tris-HCl pH 7.6, 150 mM NaCl, 2 mM EDTA pH 8, Complete™ Protease Inhibitor Cocktail (Roche), and PhosSTOP™ (Roche)). The samples were then centrifuged in an ultracentrifuge (Hitachi CP100NX) at 4°C for 20 minutes at 200,000 x g. This centrifugation yielded a supernatant containing TBS-soluble proteins, including the soluble Aβ fraction, which was stored at −80°C for later analysis. The pellet containing the insoluble Aβ was also obtained and frozen to facilitate extraction of the insoluble Aβ. Next, 125 µL of guanidine hydrochloride (GuHCl) buffer (6 M GuHCl in 50 mM Tris-HCl pH 7.6, 150 mM NaCl, 2 mM EDTA pH 8, Complete™ Protease Inhibitor Cocktail (Roche), and PhosSTOP™ (Roche)) was added to the pellet. The pellet was disaggregated and sonicated for 30 seconds at 10% amplitude. The sample was incubated for 60 minutes at room temperature, followed by centrifugation at 4°C for 20 minutes at 200,000 x g. The resulting supernatant was diluted in TBS to reduce the final concentration of GuHCl to 0.5 M. The diluted supernatant was stored at −80°C for subsequent analysis.

#### Enzyme-linked immunosorbent assay (ELISA)

The levels of Aβ40 and Aβ42 were measured using Aβ ELISA kits (Wako, 294-62501 and 290-62601) according to the manufacturer’s instructions. Briefly, samples were diluted in sample diluent to bring Aβ40 and Aβ42 levels within the optimized working range of the Aβ40 and Aβ42 standards at known concentrations (0-100 pM) for each extraction buffer condition: soluble fraction for Aβ40 and Aβ42 = 1:3; insoluble fraction for Aβ40 = undiluted, and insoluble fraction for Aβ42 = 1:6000. After incubation with the antibody solutions, Aβ40 and Aβ42 concentrations in the samples were determined by colorimetric assay. The colorimetric reaction was terminated by adding the provided stop solution, and absorbance at 450 nm was recorded. Absorbance values were converted to pg of Aβ40 or Aβ42 per mg of tissue by comparing samples to the Aβ40 or Aβ42 standard curves in the corresponding buffer conditions, correcting for respective dilution factors, and dividing by the tissue weight.

### Flow cytometry

The right cortex area was dissected and digested with an enzyme cocktail (20 units/ml papain and 0.005% DNase in sterile EBBS (Earle’s Balanced Salt Solution), Worthington Biochemical, LK003150) for 30 minutes at 37°C. After digestion, the tissue was ground in a 2 mL glass-glass grinder, filtered through a 70 μm filter, centrifuged, and the supernatant was discarded. The pellet was resuspended in 7 mL of 35% Percoll, with 5 mL of HBSS1X carefully layered on top to create a density gradient, and centrifuged at 4°C, 800 g, for 30 minutes without braking. After centrifugation, the myelin at the gradient interface, along with the rest of the supernatant, was discarded, leaving a pellet that was resuspended in 100 μL of FACS buffer for staining. For staining samples, 100 μL of each resuspended sample in FACS buffer were stained with the appropriate antibodies at a concentration of 1:1000 for 15 minutes on ice. After this incubation, 500 μL of DAPI 1:5000 or TOPRO3 1:10000 in FACS buffer was added, and the samples were centrifuged at 1,800 rpm, 4°C, for 5 minutes, with the supernatant decanted. Finally, 400 μL of counting beads (BD Trucount™, Absolute Counting Tubes, BD, 663028) resuspended in FACS buffer at a concentration of 20 beads/μL and 100 μL of FACS buffer were added, resulting in a final volume of 500 μL for analysis by flow cytometry. To evaluate the phagocytic capacity of microglia, the following antibodies were used: CD11b PE (BioLegend, 101208) and CD45 FITC (BD Biosciences, 553080), together with methoxy-X04 labeling of amyloid plaques, allowing the identification of microglia that had internalized fibrillar Aβ. To assess the cell types sequenced in the scRNA-seq, the following antibodies were used: CD11b VB510/50 (BioLegend, 101263), CD45 PE-Cy7 (BioLegend, 103114), ACSAII APC-Vio 770 (Miltenyi Biotec, 130-116-247), and O4 PE (Miltenyi Biotec, 130-117-357).

### AhR-responsive luciferase reporter assay

15,000 human embryonic kidney HEK-293 cells per well were plated in 96-well flat-bottom plates. Twenty-four hours after plating, cells were transfected with the transcription factor reporter from the XRE-Cignal Reporter Assay Kit (Qiagen, CCS-2045L). The reporter is a mixture of an inducible transcription factor-responsive construct and a constitutively expressing Renilla luciferase construct (40:1). The inducible transcription factor-responsive construct encodes the firefly luciferase reporter gene under the control of xenobiotic response elements (XREs), while the constitutively expressing Renilla construct encodes the Renilla luciferase reporter gene under the control of a CMV immediate early enhancer/promoter, acting as an internal control for normalizing transfection efficiency and monitoring cell viability. The transfection was performed using calcium phosphate. Per well, 2 μL of DNA (100 ng/μL), 1.25 μL of 2M CaCl₂, 6.25 μL of water, and 10 μL of HBSS 2X were added. After 24 hours, the transfected cells were incubated with Dulbecco’s Modified Eagle Medium (DMEM) supplemented with 10% mouse or patient’s serum, patient’s CSF, mouse brain extracts or mouse faecal pellets extracts. Mouse brain extracts or mouse faecal pellets extracts obtained by diluting 10 mg of tissue/faecal pellets in 100 µL of sterile 1X PBS and mechanically dissociating it. After this, it is centrifuged at 2,000g for 10 min at 4°C, and the supernatant is used at 10% in DMEM, in triplicates. Luciferase activity was analysed 24 hours later using the Dual-Luciferase Reporter System (Promega, E1980). Firefly luciferase activity was divided by Renilla luciferase activity and normalized to their respective control levels, which were set to 100%. To normalize basal reporter activity within each plate for the human samples, all Firefly/Renilla ratios were divided by the median value of the three DMSO control wells included on the same plate, thereby centring the DMSO response at 1. Inter-plate variation was subsequently estimated using the 6-formylindolo[3,2-b] carbazole (FICZ) concentrations included as common positive controls across plates. For each FICZ concentration, a plate-specific correction factor was calculated by dividing the global median response for that concentration across all plates by the corresponding response measured on the individual plate. Correction factors derived from the high, intermediate, and low FICZ concentrations were combined using their geometric means to obtain a single correction factor for each plate. When only two FICZ concentrations were available, the geometric mean of the two corresponding correction factors was used. Finally, each DMSO-normalized sample value was multiplied by the corresponding plate-specific correction factor. These corrected values were used for all subsequent statistical analyses.

### Single-nucleus RNA sequencing

#### Sample preparation and nuclei isolation

Single-nucleus RNA sequencing was performed using cortical tissue from 12-month-old male and female WT, APP^NL-F^ and APP^NL-F^/AhR^−/−^ mice. For each genotype, two separate sex-specific pools were prepared, one comprising cortical tissue from two male mice and the other from two female mice. Thus, six pooled samples were generated from a total of 12 animals. Each pool was processed independently for nuclei isolation, single-nucleus capture and library preparation, resulting in one sequencing library per sex-genotype combination. Frozen cortical tissue from one hemisphere was homogenized using a glass-glass homogenizer in lysis buffer containing 10 mM NaCl, 0.25 M sucrose, 10 mM Tris-HCl, 3 mM MgCl₂, 0.1 mM dithiothreitol, 1 mM CaCl₂, 0.1 % Triton X-100, 0.2 U/µL Protector RNase Inhibitor (Roche, cat. no. 03335402001), cOmplete Protease Inhibitor Cocktail (Roche) and PhosSTOP phosphatase inhibitor (Roche). Homogenates were filtered through a 70-µm cell strainer, incubated on ice for 5 min and centrifuged at 500 x g for 5 min at 4 °C. Pellets were resuspended in fresh lysis buffer, incubated on ice for an additional 5 min and centrifuged again at 500 x g for 5 min at 4 °C. The resulting nuclear pellets were resuspended in 500 µL of Wash Buffer 1, consisting of sterile 1X PBS, 2% bovine serum albumin and 0.2 U/µL Protector RNase Inhibitor, and centrifuged at 500×g for 5 min at 4 °C. Nuclei were subsequently resuspended in Wash Buffer 1 containing DAPI at a 1:5,000 dilution, filtered through a 35-µm cell strainer and centrifuged again at 500×g for 5 min at 4 °C. Pellets were resuspended in Wash Buffer 2, consisting of sterile 1×PBS, 2% bovine serum albumin and 0.4 U/µL Protector RNase Inhibitor, and transported to the CNIC Flow Cytometry Unit. Intact DAPI-positive nuclei were isolated by fluorescence-activated sorting and collected into sterile flow-cytometry tubes containing 100 µL of Wash Buffer 2. Sorted nuclei were centrifuged at 500 x g for 6 min at 4 °C and resuspended in an appropriate volume of Wash Buffer 2 to obtain a final concentration of 1,000-1,200 nuclei/µL.

#### Single-nucleus capture and library preparation

Nuclei were counted, and their integrity was assessed using a Countess 3 automated cell counter (Thermo Fisher Scientific). Each nuclear suspension was loaded into an individual well of a Chromium Next GEM Chip G (10x Genomics), targeting the recovery of 10,000 nuclei per sample. Single nuclei were encapsulated into gel bead-in-emulsions using a Chromium Controller (10x Genomics). Dual-indexed snRNA-seq libraries were prepared using the Chromium Next GEM Single Cell 3′ GEM, Library & Gel Bead Kit v3.1 (10x Genomics), following the manufacturer’s instructions. Library size distributions were assessed using a High Sensitivity DNA chip on a 2100 Bioanalyzer (Agilent Technologies), and library concentrations were measured using a Qubit fluorometer (Thermo Fisher Scientific).

#### Sequencing and FASTQ generation

Individual libraries were diluted to 10 nM, pooled and loaded for sequencing at 700 pM. Libraries were sequenced on a NextSeq 2000 system (Illumina) using either a P3 or P2 flow cell. The reported paired-end configuration was 28 bp for Read 1, 10 bp for Index 1, 10 bp for Index 2 and 90 bp for Read 2. FASTQ files for each sample were generated using the cellranger mkfastq pipeline (10x Genomics).

### Single-nuclei transcriptomic data analysis

#### Software and initial data processing

Sequence alignment and gene-expression quantification were performed independently for each sample using the CellRanger pipeline (version 7.0.1), including introns in the quantification, based on the mouse GRCm38 genome assembly and Ensembl genebuild (version 98), from which pseudogenes and small RNAs were removed. Ambient background was removed using CellBender. Nuclei were filtered and clustered using scuttle (v1.10.3) and Seurat (v4.4.0). Nuclei were filtered using a sequencing depth between 1,500 and 80,000, a minimum of 2,000 detected genes, a mitochondrial content below 15%, a cell-count fraction above 0.2% (the proportion of UMIs contributed by each cell to the total UMIs in the sample), a gene-expression complexity (percentage of reads from the top 50 genes) below 65%, and a haemoglobin gene-set expression below 2%, in order to remove low-quality nuclei. Doublets were identified using scDblFinder (v1.14.0)^60^ and removed. Mitochondrial and haemoglobin genes were excluded from downstream analysis. Counts were normalized using the LogNormalize method with a scale factor of 10,000, and the 2,000 most variable genes were identified using VST. Subsequent downstream analyses and visualization were performed in R version^61^ 4.5.1 using Seurat version 5.3.0. ^62, 63^

#### Cell-type annotation and neuronal classification

Cortical nuclei were annotated using the Allen Brain Atlas MapMyCells web tool (RRID:SCR_024672), applying hierarchical mapping against the Whole Mouse Brain reference taxonomy (CCN20230722)^64, 65^. Nuclei assigned to neuronal populations were retained for subsequent analyses and classified into GABAergic and glutamatergic neurons according to their reference-based annotations. The relative proportions of these two neuronal populations were calculated for each genotype.

#### Differential gene-expression analysis

Genotype-dependent transcriptional differences were investigated by comparing neuronal nuclei from APP^NL-F^/AhR^−/−^ and APP^NL-F^/AhR^+/+^ mice. Initial analyses were conducted separately for GABAergic and glutamatergic populations. Given the broadly similar genotype-associated pathway enrichment patterns observed across these neuronal classes, an additional analysis was performed on the combined neuronal population. Differential gene expression was assessed using Wilcoxon rank-sum tests implemented in Seurat. Genes with adjusted p-values below 0.05 were considered differentially expressed. To visualize the overall transcriptional response to AhR deletion, the 20 most upregulated and 20 most downregulated genes were selected according to their log_2_ fold changes among significantly differentially expressed transcripts. Genes with adjusted p-values reported as zero were excluded from this ranking. These genes were represented in dot plots displaying the magnitude and statistical significance of differential expression. Additional dot plots were generated for selected genes associated with neurodegeneration-related processes, including neuronal apoptosis, synaptic function, neuroinflammation, extracellular matrix remodelling, vascular responses and neurotrophic signalling. Volcano plots were also generated to visualize genome-wide differential expression and highlight genes of functional interest.

#### Gene set enrichment analysis

Gene set enrichment analysis (GSEA) was performed using the fgsea package to identify genotype-associated changes in neuronal biological processes. Analyses focused on selected Gene Ontology pathways relevant to neuronal function and neurodegeneration, including synaptic transmission, neurotransmitter release, synapse organization, cytoskeletal dynamics, neuronal apoptosis and cellular stress responses. Enrichment was initially examined separately in GABAergic and glutamatergic neurons and subsequently in the combined neuronal population. Enrichment results were expressed as normalized enrichment scores (NES), and p-values were adjusted for multiple testing using the Benjamini-Hochberg procedure. Pathways with adjusted p-values below 0.05 were considered significantly enriched.

#### Sex-stratified transcriptional analysis

To investigate potential sex-dependent effects of AhR deficiency, additional GSEA analyses were performed separately in male and female mice, retaining the distinction between GABAergic and glutamatergic neurons. Genotype-associated enrichment patterns were compared across four neuronal groups: male GABAergic, male glutamatergic, female GABAergic and female glutamatergic neurons. A common set of neurodegeneration- and synaptic function-related pathways was examined across all four groups, and the corresponding NES values were visualized in a heatmap to identify shared and sex-dependent transcriptional responses. Given that each sequencing library represented a pool of two animals and only one pooled sample was available per sex-genotype combination, differential gene-expression and gene set enrichment analyses were considered exploratory. Individual nuclei were not treated as independent biological replicates, and transcriptomic findings were interpreted as descriptive, hypothesis-generating observations rather than definitive genotype-or sex-dependent effects.

### Single-cell RNA sequencing

#### Animals and experimental design

Single-cell RNA sequencing was performed using 12-month-old male and female APP^NL-F^/AhR^+/+^ and APP^NL-F^/AhR^-/-^mice. Each sex-genotype group comprised two biological replicates, with each replicate corresponding to one individual mouse.

#### Isolation, sorting, and sample tagging of non-neuronal cells

Cortical tissue was mechanically dissociated, and non-neuronal cells were enriched using a Percoll density gradient. Approximately 100,000 viable cells from each biological sample were isolated at the CNIC Flow Cytometry Unit using a FACS Fusion cell sorter. Viable cells were selected based on the exclusion of DAPI. Sorted cells were collected in Eppendorf tubes and centrifuged at 2,800 rpm for 5 min at 4 °C. After removal of the supernatant, the cell pellet from each sample was resuspended in 190 µL BD Pharmingen Stain Buffer (Cat. No. 554656) supplemented with 10 µL of a sample-specific oligonucleotide-conjugated Sample Tag (Custom BD® Single-Cell Multiplexing Set, Rat anti-mouse MHC-H2 Clas I M1/42, Mat. No. 626545). Suspensions were transferred to DNA low-binding tubes (DNA LoBind® Tubes, Eppendorf®, 022431021) and incubated for 20 min at room temperature. Cells were washed twice with 1 mL of staining buffer. Each wash was followed by centrifugation at 2,800 rpm for 5 min at 4 °C and removal of the supernatant. The individually tagged cell suspensions were then pooled and resuspended in a final volume of 100 µL of sample buffer. The pooled suspension was transported on ice to the CNIC Genomics Unit for single cell capture and library preparation.

#### Single-cell capture and library preparation

Cell viability and concentration in the pooled Sample Tag-labelled suspension were assessed using a Countess III automated cell counter (Thermo Fisher Scientific). Up to 60,000 cells were loaded into a cartridge of the BD Rhapsody Single-Cell Analysis System, and cell capture and cDNA synthesis were performed according to the manufacturer’s instructions. Briefly, individual cells were distributed by gravity into nanowells and lysed. Released mRNA molecules and Sample Tag oligonucleotides were captured by barcoded beads present in the nanowells. Each bead contained a unique cell-label oligonucleotide that allowed transcripts and Sample Tags originating from the same cell to be identified. Beads were subsequently recovered from the cartridge, and reverse transcription was performed in a single pooled reaction. Separate indexed libraries were prepared for whole-transcriptome analysis and Sample Tag demultiplexing according to the manufacturer’s protocol. Library size distributions were assessed using a 2100 Bioanalyzer (Agilent Technologies), and library concentrations were measured using a Qubit fluorometer (Thermo Fisher Scientific). Whole-transcriptome and Sample Tag libraries were combined and sequenced in a paired-end 60x42-bp run using a NextSeq 2000 sequencing system with a P2 flow cell (Illumina). Primary sequencing output was processed using NextSeq 1000/2000 Control Software Suite version 1.4.1. FASTQ files were generated using BCL Convert version 3.6.3 (Illumina).

### Single-cell transcriptomic data analysis

#### Software and initial processing

Software and initial processing: Sequence alignment, unique molecular identifier (UMI) quantification, and single-cell identification were performed using the BD Rhapsody Sequence Analysis pipeline (version 2.2), based on the mouse GRCm39 genome assembly and Ensembl genebuild (version 108), from which pseudogenes and small RNAs were removed. This initial processing yielded a matrix of 29,386 genes and 12,571 cells. Single cells were filtered and clustered using scuttle (v1.10.3) and Seurat (v4.4.0). Cells were filtered using a sequencing depth between 2,000 and 40,000, a minimum of 1,000 detected genes, a mitochondrial content below 20%, a ribosomal content below 10%, a gene-expression complexity (percentage of reads from the top 50 genes) below 50%, a haemoglobin gene-set expression below 1%, and an HTO content above 50 UMIs in the hashtags. Doublets were identified using scDblFinder (v1.14.0) and removed. Counts were normalized using the LogNormalize method with a scale factor of 10,000, and the 2,000 most variable genes were identified using VST. Sample Tag libraries were used to assign individual cells to their corresponding biological samples. Subsequent downstream analyses were performed in R version 4.5.1 using Seurat version 5.3.0, with cells analysed according to their sex and genotype of origin^61, 62, 66^.

#### Cell-type annotation

Cells were annotated by hierarchical mapping with the Allen Brain Atlas MapMyCells web tool (RRID:SCR_024672), using the Whole Mouse Brain taxonomy CCN20230722 as reference^64, 65^. MapMyCells annotations were used to identify the major non-neuronal populations, including microglia, oligodendrocytes, endothelial cells, pericytes, astrocytes, border-associated macrophages, monocytes, dendritic cells, and lymphoid cells.

#### Microglial subsetting and reclustering

Cells assigned to the “334 Microglia NN” subclass by MapMyCells were extracted from the complete dataset and reanalysed using Seurat. Highly variable features were recalculated using FindVariableFeatures(), and expression values were rescaled using ScaleData(). Dimensionality reduction, graph-based clustering and UMAP visualization were subsequently repeated on the microglial subset using the standard Seurat workflow. Graph-based clustering was performed at a resolution of 0.2, resulting in eight transcriptionally distinct microglial subclusters, designated MgC0-MgC7. Cluster-specific marker genes were identified using FindAllMarkers() with a Wilcoxon rank-sum test. Genes detected in at least 20% of the cells in either the cluster of interest or the comparator population were considered. Each microglial subcluster was compared with all remaining microglial cells to identify cluster-associated genes. Seurat-adjusted p values were calculated using Bonferroni correction based on the total number of genes analysed. Genes with adjusted p≤0.05 were considered statistically significant.

#### Differential gene-expression analysis

Differential gene-expression analysis between experimental groups was performed at the single-cell level using Wilcoxon rank-sum tests implemented in Seurat. The following pairwise comparisons were examined: male APP^NL-F^/AhR^+/+^ versus male APP^NL-F^/AhR^-/-^ microglia; female APP^NL-F^/AhR^+/+^ versus female APP^NL-F^/AhR^-/-^microglia; male versus female APP^NL-F^/AhR^+/+^ microglia; and male versus female APP^NL-F^/AhR^-/-^ microglia. Sex-linked genes were excluded from the indicated sex-dependent comparison when specified. For group-level comparisons, p values were adjusted using the Benjamini-Hochberg false-discovery-rate procedure. Genes with adjusted p≤0.05 were considered differentially expressed. No additional log2 fold-change threshold was applied. Given the limited number of biological replicates per sex-genotype group (n=2 mice), these single-cell differential-expression analyses were interpreted as exploratory and not as definitive evidence of genotype-or sex-dependent effects at the animal level. Differential expression between microglial subclusters was assessed using analogous Wilcoxon rank-sum tests. Unless otherwise indicated, genes were required to be detected in at least 20% of the cells in one of the populations being compared. Depending on the specific analysis and Seurat function used, correction for multiple comparisons was performed using either the Benjamini-Hochberg or Bonferroni procedure. Heatmaps show row-scaled expression values for selected differentially expressed genes across individual microglial cells. Volcano plots display log_2_ fold changes and −log_10_-adjusted p values.

#### Ingenuity Pathway Analysis

Ingenuity Pathway Analysis (IPA; QIAGEN) was used to identify canonical pathways and biological functions associated with the differentially expressed genes obtained from the sex- and genotype-dependent comparisons. Upregulated and downregulated genes were analysed separately. Enrichment significance was assessed using Fisher’s exact test with Benjamini-Hochberg correction for multiple comparisons. The IPA activation z-score was used to predict pathway activation or inhibition when sufficient information regarding the expected direction of gene regulation was available. Absolute z-scores≥2 were considered indicative of predicted pathway activation or inhibition.

##### Gene Ontology over-representation analysis

Gene Ontology (GO) over-representation analysis was performed using clusterProfiler version 4.16.0 and org.Mm.eg.db version 3.21.0. For each microglial subcluster, genes with adjusted p≤0.05 and positive log_2_ fold change relative to the remaining microglial cells were included. All genes available in the annotation database were used as the background universe. Over-representation was assessed using a hypergeometric test, and p values were adjusted using the Benjamini-Hochberg procedure. GO biological processes with adjusted p≤0.05 were considered significantly enriched. Gene ratio was defined as the proportion of genes in the input list assigned to the corresponding GO term.

### Gene-set enrichment analysis

Gene-set enrichment analysis was performed using fgsea version 1.34.2 with 1,000 permutations. Wilcoxon rank-sum statistics were calculated using presto version 1.0.0. Depending on the analysis, genes were ranked either according to the area under the receiver-operating-characteristic curve derived from the Wilcoxon test or using a composite ranking metric incorporating differential-expression significance and log fold change. Enrichment was assessed against the human Hallmark collection (h.all.v2024.1.Hs.symbols.gmt), a custom phagocytosis-related gene set, the Gene Ontology Biological Process phagocytosis gene set, IPA CLEAR Signaling pathway molecules and a curated list of AhR target genes obtained from Harmonizome 3.0. Enrichment results were summarized using normalized enrichment scores. Positive and negative normalized enrichment scores indicate enrichment toward opposite ends of the ranked gene list. Enrichment p values were adjusted using the Benjamini-Hochberg procedure.

#### Gene-module scoring

Transcriptional module scores were calculated at the single-cell level using the Seurat AddModuleScore() function. Gene sets associated with disease-associated microglia, neurodegeneration and interferon responses were compiled from previously published studies and Gene Ontology resources. For each signature, the average expression of its constituent genes was calculated relative to control genes with comparable overall expression levels. Module-score distributions were examined across MgC0-MgC7.

#### Gene-set overlap analysis

Overlaps among differentially expressed genes associated with MgC0-MgC7 were visualized using UpSet plots. An additional focused analysis compared genes associated with MgC0, MgC1 and the remaining microglial subclusters, collectively designated “Other”.

### Faecal microbiota collection and DNA extraction

Faecal pellets were collected from male and female WT and APP^NL-F^ mice at 5 and 12 months of age and stored at −80 °C until processing. Microbial DNA extraction, 16S rRNA gene amplification, library preparation and sequencing were performed by Macrogen Corea. Microbial DNA was extracted from approximately 250 mg of faecal material using the DNeasy PowerSoil Pro Kit in a 96-well format (QIAGEN), following the manufacturer’s protocol. Samples were transferred to a PowerBead Pro Plate, and 850 µL of Solution CD1 was added to each well. Plates were incubated at 65 °C for 10 min and subsequently shaken at 20 Hz for 10 min using a 96-well plate shaker. After centrifugation at 2,900 x g for 3 min, 600 µL of supernatant was transferred to a collection plate, mixed with 200 µL of Solution CD2 by ten inversions and centrifuged again at 2,900xg for 3 min. Up to 700 µL of the resulting supernatant was transferred to a new plate and mixed with 600 µL of Solution CD3. Lysates were loaded in successive 650-µL aliquots onto a QIAamp 96 Spin Plate and centrifuged at 2,900 x g for 3 min after each loading step. The columns were washed sequentially with 500 µL of Solution EA and 500 µL of Solution C5, followed by centrifugation at 2,900 x g for 5 min and air-drying for 10 min at room temperature. DNA was eluted by adding 50 µL of Solution C6 to each well, incubating for 3 min and centrifuging at 2,900xg for 3 min.

### 16S rRNA gene amplification, library preparation and sequencing

Amplicon libraries targeting the V3-V4 hypervariable regions of the bacterial 16S rRNA gene were prepared by Macrogen Spain using its validated Metagenome Amplicon library-preparation workflow. The following primers were used:

• Bakt_341F: 5′-CCTACGGGNGGCWGCAG-3′

• Bakt_805R: 5′-GACTACHVGGGTATCTAATCC-3′

PCR amplification, amplicon purification, indexing and library pooling were performed by Macrogen Spain according to its standardized protocols. Library quality and concentration were assessed before sequencing. Pooled libraries were sequenced in a dedicated lane of an Illumina MiSeq system, with an expected total output of 5 Gb, using a 2x300-bp paired-end configuration and a 30% PhiX spike-in.

### Bioinformatic analysis of microbiota

Sequence analysis was performed mostly with mothur^67^, following the standard operating procedure^68^ for 16S rRNA-gene paired-end reads generated with Illumina’s MiSeq platform. In brief, the workflow consisted in assembling R1 and R2 reads, filtering merged sequences by length and quality, aligning selected sequences against a customized version of the 16S SILVA SEED v132 reference to select those corresponding to regions V3-V4, clustering fragments to reduce redundancy and to denoise (pre.cluster algorithm), removing chimeras with the integrated uchime algorithm, and classifying fragments with the integrated Naive Bayes classifier against SILVA nr v138_1. Sequences were grouped into phylotypes based on their taxonomic classification, and an abundance matrix containing genus-level counts for each phylotype in each sample was generated. Customized R scripts were then used to generate rarefaction curves to assess sequencing depth, to calculate alpha and beta diversity indices and to perform Bray-Curtis’s distance-based Principal Coordinate (PCoA) and PERMANOVA analyses, using package vegan, as well as to produce exploratory plots from relative abundance (RA) values. Differential abundance analyses were performed with package MaAsLin2^69^, using the LM analysis method on TSS-normalized, log transformed RA values, after discarding low-abundance taxa with RA <0.0001 and requiring presence in at least n samples, where n was automatically determined as a function of group size. Differentially abundant taxa were defined as those with FDR <0.25. Pathway abundance inference for each metagenome was generated with PICRUSt2^70^, to infer EC numbers and MetaCyc pathway abundances. Differential pathway abundance tests were performed with MaAsLin2.

### Metabolomics analysis

Metabolomic analyses were performed at the Centre for Metabolomics and Bioanalysis (CEMBIO), Facultad de Farmacia, Universidad San Pablo-CEU, CEU Universities, Urbanización Montepríncipe, 28660 Boadilla del Monte, Spain. Serum and faecal extracts were analyzed by capillary electrophoresis coupled to time-of-flight mass spectrometry (CE-TOF-MS) in positive-ionization mode using an Agilent 7100 CE system coupled to an Agilent 6230 TOF mass spectrometer. Sample preparation, analytical conditions, quality-control procedures and data preprocessing followed previously validated CE-TOF-MS workflows, with the modifications detailed in the Supplementary Methods^71, 72^. L-Tryptophan and choline chloride were semi-quantified by CE-TOF-MS in positive-ionization mode using the same analytical conditions. Indole-3-aldehyde (IAld; indole-3-carboxaldehyde), indole-3-acetamide (I3AM), trans-indole-3-acrylic acid (I3Acr), and n-butyric acid were semi-quantified by UHPLC-MS/MS using an Agilent 1290 Infinity II UHPLC system coupled to an Agilent 6495D triple-quadrupole mass spectrometer operating in dynamic multiple-reaction-monitoring mode. Semi-quantitative concentrations were estimated using analyte-specific eight-point external calibration curves prepared in solvent. Detailed sample-preparation procedures, instrumental conditions, quality-control criteria, calibration ranges, and compound-specific analytical parameters are provided in the Supplementary Methods. Metabolomic data were analysed using univariate statistical approaches. The selected pairwise comparisons presented in Figure 7H were evaluated using two-sided non-parametric tests, with p values adjusted for multiple comparisons using the Benjamini-Hochberg false-discovery-rate procedure. Adjusted q values below 0.05 were considered statistically significant. For visualization in Figure 7H, metabolite values were standardized separately for each analyte and biological matrix (faeces or serum). Z-scores were calculated by subtracting the mean and dividing by the sample standard deviation across the corresponding 12-month dataset, and group-mean Z-scores were then calculated for each sex-genotype group. This transformation was used solely to display relative profiles across analytical methods and matrices and does not represent statistical integration or directly comparable absolute metabolite concentrations. Principal component analysis (SIMCA version 17) was used to evaluate overall data structure, quality-control clustering and potential outliers.

### Human cohorts and biomarker classification

The COGNIBIO12 clinical cohort is an ongoing prospective memory-clinic cohort established in September 2023 at Hospital Universitario 12 de Octubre, Madrid, Spain, with more than 300 participants enrolled. For the present retrospective study, an analytical subcohort of 142 participants was identified from linked clinical and biospecimen records based on the availability of archived biological samples selected for AhR bioactivity assessment and associated clinical information. Because the analytical dataset incorporated archived specimens collected before the formal establishment of COGNIBIO12, the 142 participants do not represent a consecutively recruited post-September 2023 sample. Participants had been evaluated for subjective or objective cognitive complaints and received heterogeneous clinical diagnoses, including Alzheimer’s disease, primary progressive aphasia, behavioural-variant frontotemporal dementia, dementia with Lewy bodies, vascular cognitive impairment, posterior cortical atrophy, corticobasal syndrome, mixed dementia and other neurological conditions.

In the COGNIBIO12 analytical subcohort (n=142), mutually exclusive clinical diagnostic codes comprised Alzheimer’s disease (85, 59.9%; 59 women/26 men), primary progressive aphasia (9, 6.3%; 5/4), behavioural-variant frontotemporal dementia (7, 4.9%; 4/3), dementia with Lewy bodies (10, 7.0%; 4/6), mixed clinical diagnosis (4, 2.8%; 1/3), vascular cognitive impairment (2, 1.4%; 0/2), posterior cortical atrophy (2, 1.4%; 2/0), corticobasal syndrome (1, 0.7%; 0/1), other clinical diagnoses (13, 9.2%; 4/9), and missing clinical diagnosis (9, 6.3%; 3 women/3 men/3 with no recorded sex). Separately, a suspected amyloid aetiology was recorded in 87 participants (61.3%; 60 women/27 men), including 80 who also had a clinical Alzheimer’s disease diagnosis. Clinical and etiological classifications overlapped and are therefore reported separately. All percentages use 142 as the denominator, and small deviations from 100% reflect rounding. Sample collection was approved by the Research Ethics Committee with Medicinal Products (CEIm) of Hospital Universitario 12 de Octubre, Madrid (N° CEIC 21/584), which issued a favourable opinion. Serum and, when available, cerebrospinal fluid (CSF) samples were used to determine AhR-activating bioactivity. Serum AhR bioactivity was analytically valid for 125/142 patients (75 women and 50 men), and CSF bioactivity was available for 138/142 (80 women, 55 men and three with sex not recorded). Based on participant identifiers, 122 had both serum and CSF AhR measurements, three had serum-only data, 16 had CSF-only data, and one had neither valid measurement. Two serum values were excluded for technical quality-control reasons, while their CSF measurements were retained. Cross-compartment analyses included only observations matched within the same participant; other analyses used the available data for each specific variable.

Cognitively unimpaired participants from the independently recruited Atherobrain–Heart to Head (H2H) cohort constituted the external reference group (n=28; 16 women and 12 men; age 73.5 ± 5.3 years, mean ± SD). The H2H cohort was prospectively and consecutively recruited between June 2017 and December 2018, as previously reported^73^. Cognitively unimpaired status was defined by an MMSE score >27, absence of clinical signs of cognitive impairment and preserved functional status. For the present study, archived serum samples were available from 28 H2H participants; CSF samples from H2H participants were not analysed in this study.

Participants with available CSF AD biomarkers were classified according to the NIA-AA AT(N) research framework^74^. Amyloid and tau positivity were defined using in-house cutoffs established by the Clinical Biochemistry laboratory of Hospital Universitario 12 de Octubre for the Lumipulse® G System (CSF Aβ42 ≤ 599 pg/mL; p-tau181 ≥ 56.5 pg/mL), generating four groups: A−T−, A+T−, A−T+ and A+T+. Both CSF Aβ42 and p-tau181 were available in 93 participants, yielding A−T− (n=11), A+T− (n=10), A−T+ (n=15) and A+T+ (n=57). Amyloid-positive participants were also analysed jointly, irrespective of tau status, and tau-positive participants were analysed irrespective of amyloid status. CSF Aβ40 measurements were not available for all participants because of limited assay-kit availability. Initially, Aβ40 quantification was prioritized for patients with a clinical diagnosis of Alzheimer’s disease, and subsequent measurements were performed sequentially as samples were processed. Consequently, the availability of the CSF Aβ42/Aβ40 ratio was not based on random sampling. The primary AT classification was based on CSF Aβ42 (≤599 pg/mL) and p-tau181 (≥56.5 pg/mL), which were available in a larger number of participants. The CSF Aβ42/Aβ40 ratio (<0.069) was used only in exploratory sensitivity analyses restricted to participants with both measurements available. The analytical subcohort was not restricted to a clinical Alzheimer’s disease diagnosis or amyloid biomarker positivity. The dataset comprised individuals identified from linked clinical and/or biospecimen records; availability of both specimen matrices was not mandatory. Visit-matched MMSE and GDS data were available for n=42 and n=91, respectively, distinct from broader longitudinal records.

### Human sample collection and processing

Blood serum was obtained by centrifuging blood samples for 5 minutes at 2,000 g at room temperature. The supernatant was then transferred to a new tube and stored at −80 °C until further use. CSF samples were obtained by lumbar puncture following the established European preanalytical protocol. Different CSF volumes were collected depending on the analyses to be performed. An initial volume of 1-2 mL was allocated for basic determinations, including cell count, glucose, and protein levels. An additional 1.5-2 mL was subsequently collected for the assessment of biomarkers in patients, including β-amyloid, tau, and p-tau. Finally, 7-8 mL of CSF was collected into sterile polypropylene tubes (Sarstedt, 10 mL) for processing and storage. Samples were processed within a maximum of 2 hours after collection. CSF was centrifuged at 2,000 g for 10 minutes at 4 °C to remove potential cellular debris. Following centrifugation, the supernatant was distributed into pre-labelled 500-µL aliquots. Samples were finally stored at −80 °C until further analysis. The H2H samples came from an independently recruited reference cohort whose recruitment and eligibility criteria were previously reported^73^. These collection and preanalytical procedures describe the COGNIBIO12 samples. All human specimens were assayed for functional AhR-activating bioactivity using the XRE-luciferase method described above.

### Statistical analysis for preclinical data

Statistical analyses were performed using GraphPad Prism version 10.0.2 (GraphPad Software, Boston, MA, USA). Data are presented as mean±SEM unless otherwise indicated, and individual data points represent biologically independent animals unless explicitly stated otherwise. Normality was assessed using the Shapiro-Wilk test. For morphometric analyses involving multiple measurements per animal, individual plaque or cell measurements were nested within mice, which were considered the biological experimental units. Retained measurements were averaged within each mouse, and mouse-level group means were compared using two-sided Welch’s tests. For outcomes involving three groups, pairwise genotype comparisons were adjusted using Holm’s procedure within each outcome. Additional sex-adjusted analyses were performed at the animal level using linear models including genotype and sex as fixed predictors with HC3 robust standard errors. Parametric tests were used when the data satisfied the corresponding distributional assumptions; otherwise, non-parametric alternatives were applied. Comparisons between two independent groups were performed using two-tailed unpaired Student’s t tests for normally distributed data or two-tailed Mann-Whitney U tests for non-parametric data; exact Mann-Whitney U p values were reported where applicable. Comparisons involving more than two independent groups were performed using one-way ANOVA followed by the appropriate multiple-comparisons procedure or, for non-parametric data, the Kruskal-Wallis’s test followed by Dunn’s multiple-comparisons test. Experimental designs involving two independent factors, including genotype, treatment, sex or age, were analysed using two-way ANOVA, followed by Tukey’s, Šídák’s, Dunnett’s or Fisher’s least significant difference (LSD) multiple-comparisons tests, as specified in the corresponding figure legends. Associations between continuous variables were assessed using Pearson correlation for approximately normally distributed variables with a linear relationship or Spearman rank correlation when parametric assumptions were not met. Microbiota-specific permutational multivariate analyses, transcriptomic analyses and metabolomic statistics are described in their respective Methods sections. Potential outliers were identified using the ROUT method implemented in GraphPad Prism, with a false-discovery rate set at Q=0.2% and were excluded from subsequent statistical analyses. For morphometric analyses involving multiple measurements per animal, the mouse was considered the experimental unit. Individual plaque or cell measurements were retained for visualization, whereas values were averaged within each mouse for the primary statistical comparisons between experimental groups. Two-sided Welch’s t-tests were performed on mouse-level means. For analyses involving more than two experimental groups, pairwise comparisons were adjusted using Holm’s procedure within each outcome. All statistical tests were two-sided, and p<0.05 was considered statistically significant.

### Statistical analysis for human samples

Continuous variables are presented as mean ± SD or median [interquartile range (IQR)], as appropriate, and categorical variables as counts and percentages. Differences in demographic and clinical characteristics were assessed using Student’s t-test or analysis of variance for continuous variables and χ² or Fisher’s exact tests for categorical variables. Because serum and CSF AhR-activating bioactivity showed right-skewed distributions, values were natural-log transformed before parametric analyses. Group differences were examined using linear regression models. Unadjusted models included group as the only predictor, whereas adjusted models included age and sex as covariates. Heteroscedasticity-consistent HC3 standard errors were used, and regression coefficients were exponentiated and reported as geometric mean ratios with 95% confidence intervals. Analyses included comparisons between COGNIBIO12 and the primary H2H control cohort, comparisons among the four AT groups, pooled A+ versus A− and T+ versus T− analyses, and comparisons across harmonized clinical diagnostic categories. Planned contrasts between AT groups were corrected using the Holm procedure. Kruskal-Wallis and Mann-Whitney tests were used as non-parametric sensitivity analyses. Associations between AhR bioactivity and continuous CSF biomarkers were evaluated using Pearson correlations with natural-log-transformed AhR values, Spearman rank correlations as sensitivity analyses, and linear regression models adjusted for age and sex. The association between serum AhR-activating bioactivity and CSF Aβ42 concentrations was primarily examined in amyloid-positive (A+) participants, defined according to the CSF Aβ42 cutoff. Additional exploratory sensitivity analyses were performed to assess the influence of Aβ40 availability and alternative definitions of amyloid positivity. These included repeating the primary correlation in Aβ42-defined A+ participants with available Aβ42/Aβ40 measurements and reassessing the association in participants classified as A+ according to the CSF Aβ42/Aβ40 ratio (<0.069). Given the sex-dependent effects observed in the preclinical experiments, sex-stratified analyses and formal group-by-sex or biomarker-by-sex interaction terms were also examined. Only participants with complete data for the variables included in each model were analysed, and missing values were not imputed. For families of exploratory biomarker associations, p values were additionally adjusted using the Benjamini-Hochberg false-discovery-rate procedure. All tests were two-sided, and p<0.05 was considered statistically significant. The unit of analysis was the participant. Serum-versus-CSF associations included only paired samples from the same participant (122 pairs overall; n=60 paired A+ and n=51 paired A+T+ after the corresponding biomarker requirements). Within the 93 participants with paired CSF Aβ42 and p-tau181 data, AT-classified serum AhR analyses comprised n=87 and CSF AhR analyses n=92; one additional participant with available CSF AhR lacked age data, yielding n=91 for adjusted CSF models. The pooled T+ versus T− and A+ versus A− contrasts were assessed separately from the four-level AT-group test. Complete-case selection was applied within each model without imputation; unadjusted p values and FDR-adjusted q values were distinguished explicitly.

## Acknowledgments

The authors thank all members of the laboratory for their insightful feedback. We also gratefully acknowledge the staff of the Centro Nacional de Investigaciones Cardiovasculares (CNIC) for their excellent technical support.

We thank Aranzazu Rosado Díez, Sergio Callejas Alejano, and Ana Dopazo from the Genomics Unit for their support with single-cell and single-nucleus RNA sequencing; Eva Garrido, Antonio Benítez, Alberto García, Santiago Rodríguez, and Ernesto de la Cueva from the Animal Facility for animal husbandry; Elena Prieto, Raquel Nieto, Mariano Vitón, Norman Alberto Muñoz, and María Beatriz Álvarez-Flores from the Flow Cytometry Unit for cell sorting and cytometric analyses; and Elvira Arza Cuesta, Olga Giménez Sáez, Verónica Labrador Cantarero, Alfonsa Díaz Torres, and Valeria R. Caiolfa from the Advanced Microscopy and Dynamic Imaging Unit for assistance with microscopy, image acquisition, and image analysis.

## Data And Materials Availability

The data are presented in the main manuscript and in the supplementary materials. snRNA-seq data are deposited in the Genome Expression Omnibus (GEO) with accession <u>GSE353385</u>.

