## Supplementary material for "Microglial AhR Drives Sex-Biased Amyloid Pathology": Suppl. Figures and Methods

**Nieto-Vaquero et al.**

### Supplementary Methods

### Metabolomics analysis

**Chemicals:** Ultrapure water was produced in-house using a Milli-Q Direct 16 water-purification system (Millipore, Burlington, MA, USA) and was used to prepare all aqueous buffers and standard solutions. Acetonitrile, formic acid, and methanol, all of LC–MS grade, were purchased from Fisher Scientific (Pittsburgh, PA, USA). Analytical-grade ammonia solution (28%, GPR RECTAPUR®) was obtained from VWR Chemicals (Pittsburgh, PA, USA). The internal standards used for CE–MS analysis were methionine sulfone, paracetamol, 2-(N-morpholino)ethanesulfonic acid (MES), procaine, and ethyl sulfate, all purchased from Sigma-Aldrich (Burlington, MA, USA). Analytical standards for L-tryptophan, indole-3-aldehyde (IAld; indole-3-carboxaldehyde), indole-3-acetamide (I3AM), trans-indole-3-acrylic acid (I3Acr), choline chloride, and n-butyric acid were obtained from Sigma-Aldrich. Purine, HP-0922, and ammonium trifluoroacetate, used as reference-mass compounds, were purchased from Agilent (Santa Clara, CA, USA).

**Sample treatment:** Untargeted CE–MS metabolomic profiling was performed to provide broad coverage of the polar metabolome. For serum preparation, 100 µL of serum was mixed with 100 µL of 0.05 M formic acid containing 5% (v/v) acetonitrile and the following internal standards: methionine sulfone (0.4 mM), paracetamol (2 mM), MES (0.5 mM), ethyl sulphate (0.15 mM), and procaine (0.1 mM). The mixture was transferred to a centrifugal ultrafiltration device (Pall Corporation, Port Washington, NY, USA) with a 3-kDa molecular-weight cutoff and centrifuged at 16,000×g for 60 min at 4 °C. The resulting filtrate was collected for metabolomic analysis. Because of the limited amount of faecal material available, samples within each experimental group were pooled before extraction to generate five independent pooled replicates per group. Pooled faecal samples were frozen and lyophilized overnight using a LyoQuest freeze dryer (Telstar, Barcelona, Spain) until a constant dry weight was achieved. Subsequently, 10 mg of lyophilized faecal material was resuspended in 100 µL of MeOH:$H_2O$ (1:15, v/v). After vortex mixing and centrifugation at 16,000×g for 15 min at 4 °C, 100 µL of supernatant was transferred to a clean tube and evaporated to dryness using a vacuum concentrator. Dried extracts were reconstituted in 100 µL of 0.05 M formic acid containing the same internal standards at the concentrations used for serum preparation. A pooled serum quality-control sample was prepared by combining equal volumes of all serum samples and was processed in the same manner as the study samples. Samples were randomized, and quality-control samples were injected at the beginning, throughout, and at the end of the analytical batch to monitor system performance and sample-preparation reproducibility. Two procedural blanks were prepared alongside the samples and analysed at the beginning and end of the analytical sequence[1].

**Analytical strategy:** The analytical workflow combined untargeted and targeted mass-spectrometry-based approaches. Serum and faecal extracts were subjected to untargeted CE–MS metabolomic profiling in positive-ionization mode. CE–MS was also used for the targeted semi-quantification of choline and L-tryptophan, whereas n-butyric acid, IAld, I3AM and I3Acr were analysed by UHPLC–QqQ-MS/MS. Targeted metabolites were semi-quantified using external calibration curves prepared in solvent.

### Untargeted metabolomics analysis

**Analytical settings:** Positive-ionization analyses of serum and faecal extracts were performed using an Agilent 7100 capillary-electrophoresis system coupled to an Agilent 6230 time-of-flight mass spectrometer equipped with an electrospray-ionization source. Metabolites were separated in normal polarity using a fused-silica capillary (total length, 100 cm; internal diameter, 50 µm; Agilent). Before each analysis, the capillary was rinsed for 5 min at 950 mbar with background electrolyte consisting of 1 M formic acid in 10% (v/v) methanol. Samples were injected for 50 s at 50 mbar, followed by injection of background electrolyte for 20 s at 100 mbar to improve analytical reproducibility. Separation was performed at 30 kV, with an internal pressure of 25 mbar and a constant temperature of 20 °C. The observed current was approximately 50 µA, and the total run time was 35 min. The mass spectrometer was operated in positive-ionization mode over an m/z range of 60–1,000 at an acquisition rate of 1.36 spectra/s. Additional parameters were as follows: fragmentor voltage, 125 V; skimmer voltage, 65 V; OctopoleRFPeak, 750 V; drying-gas temperature, 200 °C; drying-gas flow, 10 L/min; nebulizer pressure, 0 psig; and capillary voltage, 3,500 V. The sheath liquid consisted of MeOH:$H_2O$ (1:1, v/v) containing purine (m/z 121.0509) and HP-0922 (m/z 922.0098) as reference masses and was delivered at 0.6 mL/min with a 1:100 split ratio. Data were acquired using Agilent MassHunter Workstation LC/MS Data Acquisition for 6200 Series TOF/6500 Series Q-TOF software (version B.09.00.9044.0), and raw files were inspected using MassHunter Qualitative Analysis software (version B.08.00; Agilent).

**Data preprocessing:** Raw CE–MS data were processed using Agilent MassHunter Profinder version B.10.0.2.162 to remove background noise and unrelated ions. Features were aligned across samples according to accurate mass and migration time to generate a structured feature matrix. Features with a coefficient of variation ≥30% in quality-control samples were excluded. The filtered matrix was imported into SIMCA version 17 (Sartorius, Göttingen, Germany), and principal component analysis was used to evaluate quality-control clustering, detect potential outliers, and identify biological or analytical trends. Instrumental signal drift was corrected using quality-control sample-based support-vector-regression correction implemented in MATLAB R2022a. Corrected signals were subsequently normalized to the corresponding internal standards.

### Targeted metabolite analysis

Analytical standards and compound-specific parameters are summarized in **Table S1**. Choline chloride and L-tryptophan were analysed by CE–MS in positive-ionization mode using the electrophoretic and mass-spectrometric conditions described above and identified by comparison with authentic analytical standards. n-Butyric acid, IAld, I3AM and I3Acr were analysed by UHPLC–QqQ-MS/MS.

### UHPLC–QqQ-MS/MS analysis

Samples were analysed using an Agilent 1290 Infinity II UHPLC system coupled to an Agilent 6495D triple-quadrupole tandem mass spectrometer equipped with an Agilent Jet Stream electrospray-ionization source. Analyses were performed in positive- and negative-ionization modes. Nitrogen was used as the drying, nebulizing, and sheath gas.

Source parameters were as follows: capillary voltage, 3,000 V in positive-ionization mode and 2,500 V in negative-ionization mode; drying-gas flow, 14 L/min; drying-gas temperature, 200 °C; nebulizer pressure, 50 psi; sheath-gas flow, 12 L/min; and sheath-gas temperature, 375 °C. Data were acquired in dynamic multiple-reaction-monitoring mode. Chromatographic separation was performed using an InfinityLab Poroshell 120 HILIC-Z column (2.1×150 mm, 2.7 µm) fitted with a compatible guard column (Agilent). The injection volume was 2 µL. Mobile phase A consisted of 20 mM ammonium acetate, and mobile phase B consisted of acetonitrile. The gradient was initiated at 10% A and 90% B and maintained for 1 min. Mobile phase A was increased to 22% at 8 min, 40% at 12 min, and 90% at 15 min. These conditions were maintained until 18 min. Initial conditions were restored at 19 min and maintained until the end of the run at 23 min.

Instrument control and data acquisition were performed using Agilent MassHunter Workstation software (version 06.01). Compound-specific transitions were optimized using Agilent MassHunter Optimizer software. The transitions used for quantitative analysis are provided in **Table S1**. Targeted metabolites were semi-quantified using analyte-specific eight-point external calibration curves prepared in solvent. Concentration ranges were selected according to the linear response obtained for each compound and are reported in **Table S1**. Sample responses were interpolated against the corresponding calibration curves and expressed as estimated concentrations. Because calibration standards were prepared in solvent rather than matrix-matched material, the resulting values were considered semi-quantitative.

### Supplementary Figures and Figure Legends

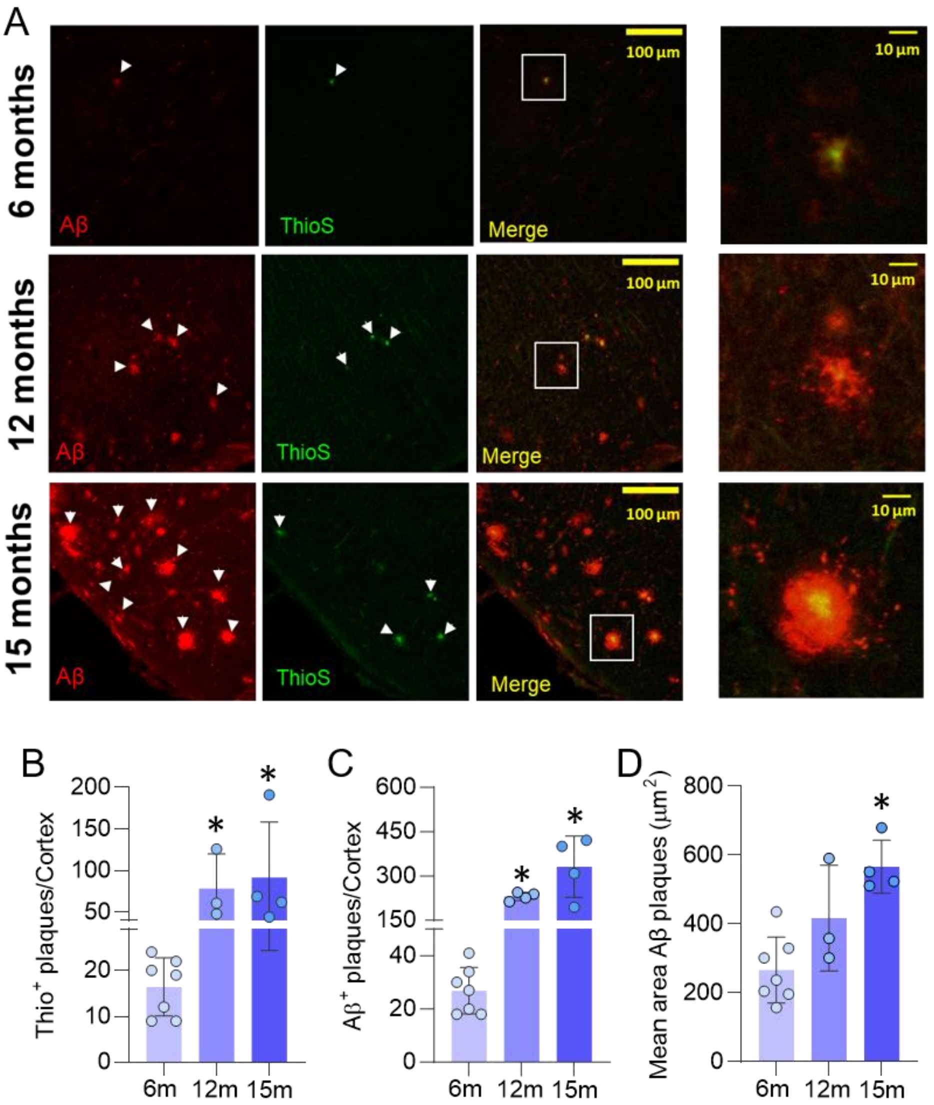


**Supplementary Figure 1. Age-dependent progression and maturation of cortical amyloid plaques in APP$^{NL-F}$ mice. (A)** Representative cortical images from 6-, 12- and 15-month-old APPNL-F mice showing total Aβ deposits detected by immunofluorescence (red), Thioflavin S-positive fibrillar amyloid cores (ThioS, green) and merged channels. Arrowheads indicate amyloid plaques. Enlarged views of the boxed regions illustrate the progressive development of compact ThioS-positive cores within Aβ-positive plaques. Scale bars = 100 µm; enlarged images, 10 µm. **(B)** Number of cortical ThioS-positive plaques at 6, 12 and 15 months of age. n=7, n=3 and n=4, respectively. Kruskal-Wallis test followed by Dunn's multiple-comparisons test. Adjusted p=0.0623 for 6 versus 12 months, p=0.0169 for 6 versus 15 months and p>0.9999 for 12 versus 15 months. **(C)** Number of cortical Aβ-positive plaques at 6, 12 and 15 months of age. n=7, n=4 and n=4, respectively. Kruskal-Wallis test followed by Dunn's multiple-comparisons test. Adjusted p=0.0609 for 6 versus 12 months, p=0.0072 for 6 versus 15 months and p>0.9999 for 12 versus 15 months**.** **(D)** Mean size of cortical Aβ-positive plaques at 6, 12 and 15 months of age. n=7, n=3 and n=4, respectively. Kruskal-Wallis test followed by Dunn's multiple-comparisons test. Adjusted p=0.3120 for 6 versus 12 months and p=0.0124 for 6 versus 15 months. Data are presented as mean±SEM. Each dot represents one biologically independent mouse. Statistical significance was considered at p<0.05.

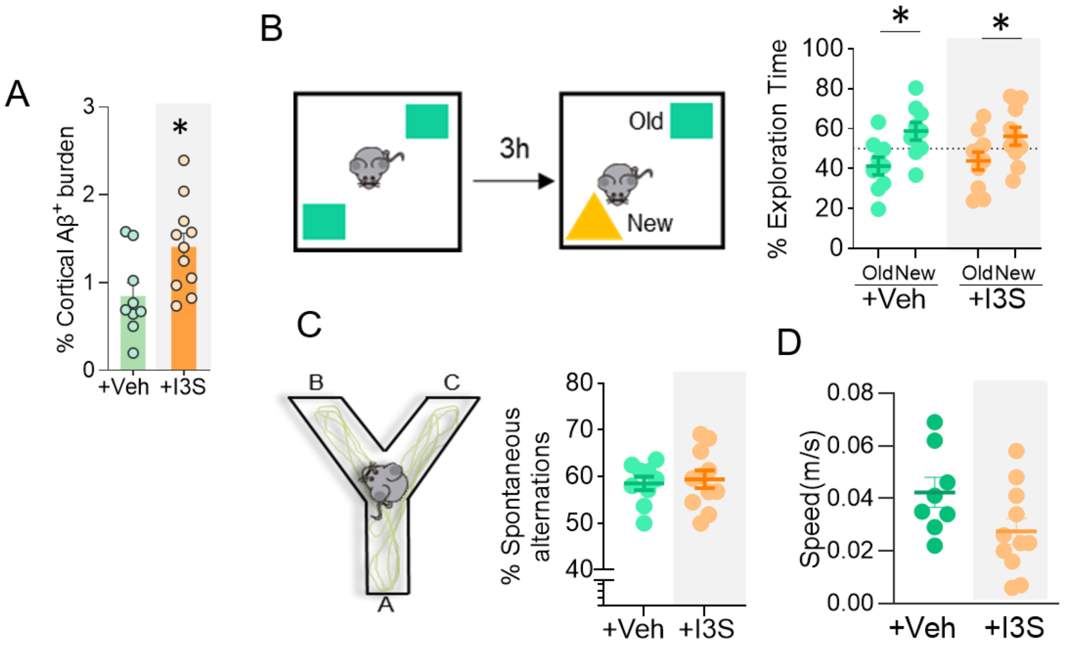


**Supplementary Figure 2. Additional amyloid and behavioural effects of indoxyl-3-sulfate treatment in APP$^{NL-F}$ mice. (A)** Percentage of cortical area occupied by Aβ-positive deposits in 12-month-old APP$^{NL-F}$ mice treated with vehicle or indoxyl-3-sulfate (I3S). Vehicle: n=9; I3S: n=11. Two-tailed exact Mann-Whitney U test, p=0.0159. **(B)** Schematic representation of the novel object recognition test and percentage of total exploration time spent investigating the familiar and novel objects in vehicle- and I3S-treated APP$^{NL-F}$ mice. Testing was performed 3h after the familiarization phase. Mice in both treatment groups spent a significantly greater proportion of time exploring the novel object than the familiar object (p<0.05). Vehicle: n=9; I3S: n=10. Two-way ANOVA followed by Sidák's multiple-comparisons test. **(C)** Schematic representation of the Y-maze and percentage of spontaneous alternations in vehicle- and I3S-treated APP$^{NL-F}$ mice. No significant difference was observed between treatment groups. **(D)** Mean speed during the open-field test in vehicle- and I3S-treated APP$^{NL-F}$ mice. Vehicle: n=8; I3S: n=11. Two-tailed exact Mann-Whitney U test, p=0.0714. Data are presented as mean±SEM. Each dot represents one biologically independent mouse. Statistical significance was considered at p<0.05.

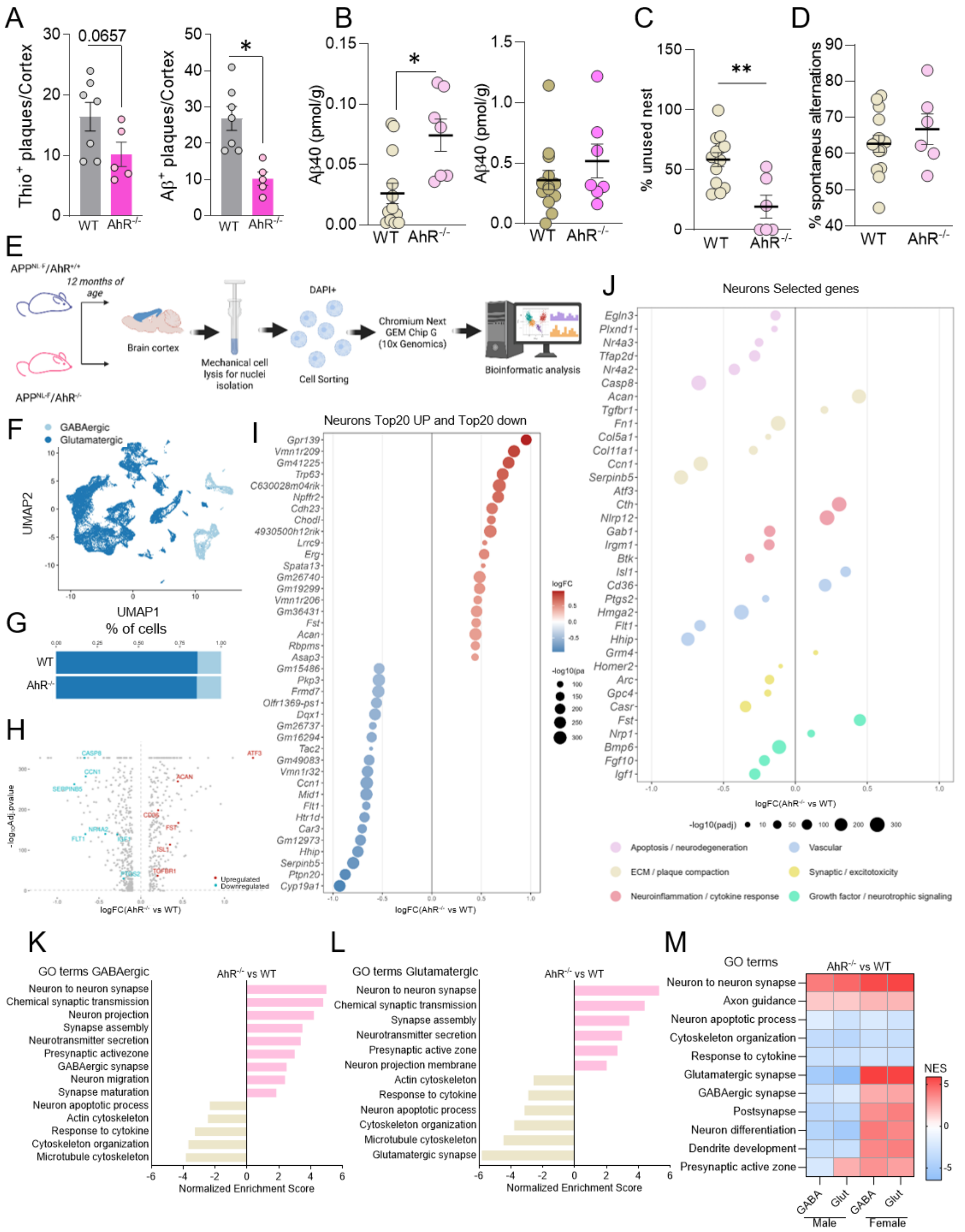

**Supplementary Figure 3. Constitutive AhR deficiency reduces early cortical amyloid plaque burden in APP$^{NL-F}$ mice. (A)** Left, number of cortical ThioS-positive plaques in 6-month-old APP$^{NL-F}$/AhR$^{+/+}$ and APP$^{NL-F}$/AhR$^{-/-}$ mice. n=7 and n=5, respectively. Two-tailed exact Mann-Whitney U test, p=0.0657. Right, number of cortical Aβ-positive plaques in 6-month-old APP$^{NL-F}$/AhR$^{+/+}$ and APP$^{NL-F}$/AhR$^{-/-}$ mice. APP$^{NL-F}$/AhR$^{+/+}$: n=7; APP$^{NL-F}$/AhR$^{-/-}$: n=5. Two-tailed exact Mann-Whitney U test, p=0.0025. **(B)** Cortical Aβ40 concentrations measured by ELISA in the TBS-soluble and guanidine-HCl-extracted insoluble fractions in APP$^{NL-F}$/AhR$^{+/+}$ and APP$^{NL-F}$/AhR$^{-}$

$^{/-}$ mice. **(C)** Percentage of unused nesting material in 12-month-old $APP^{NL-F}/AhR^{+/+}$ and $APP^{NL-F}/AhR^{-/-}$ mice. n=13 and n=6, respectively. Two-tailed exact Mann–Whitney U test, p=0.0066. **(D)** Percentage of spontaneous alternations in Y-maze in 12-month-old $APP^{NL-F}/AhR^{+/+}$ and $APP^{NL-F}/AhR^{-/-}$ mice. n=14 and n=6, respectively. Two-tailed exact Mann–Whitney U test, p=0.7181. Data are presented as mean±SEM. Each dot represents one biologically independent mouse. Sex distribution in (D) was as follows: $APP^{NL-}F/AhR^{+/+}$, 7 males/7 females; $APP^{NL-F}/AhR^{-/-}$, 4 males/2 females. **(E)** Schematic of the single-nucleus RNA-sequencing (snRNA-seq) workflow. Cortices from 12-month-old $APP^{NL-F}/AhR^{+/+}$ and $APP^{NL-F}/AhR^{-/-}$ mice were mechanically dissociated, DAPI-positive nuclei were isolated by cell sorting, and single-nucleus transcriptomes were captured using the 10x Genomics Chromium Next GEM Chip G platform followed by bioinformatic analysis. **(F)** Uniform Manifold Approximation and Projection (UMAP) of cortical nuclei, coloured by neuronal subtype (GABAergic, light blue; Glutamatergic, dark blue). **(G)** Proportion of GABAergic and Glutamatergic nuclei per genotype. **(H)** Volcano plot showing differentially expressed genes (DEGs) in cortical neurons from $APP^{NL-F}/AhR^{-/-}$ vs. $APP^{NL-F}/AhR^{+/+}$ mice. Each point represents a detected transcript, with the x-axis indicating log2 fold-change and the y-axis indicating -log10(adjusted p-value). Differential expression was assessed using the Wilcoxon rank-sum test, with adjusted p<0.05 considered statistically significant. Selected genes are highlighted and labelled according to their functional relevance. **(I)** Dot plot of the top 20 most upregulated and top 20 most downregulated genes (by log2 fold-change) in cortical neurons from $APP^{NL-F}/AhR^{-/-}$ vs $APP^{NL-F}/AhR^{+/+}$ mice, identified across all detected transcripts by Wilcoxon rank-sum test (adjusted p<0.05; genes with padj=0 excluded). Dot colour represents log2 fold-change; dot size represents -log10(adjusted p-value). **(J)** Dot plot of selected differentially expressed genes associated with the functional categories indicated in the legend. Genes were selected from significant differentially expressed transcripts (adjusted p<0.05) based on their functional relevance to the indicated categories. Dot colour represents functional category, and dot size represents -log10(adjusted p-value). **(K)** Gene set enrichment analysis (GSEA) of curated synaptic and neurodegeneration-associated pathways in GABAergic neurons, $APP^{NL-F}/AhR^{-/-}$ vs $APP^{NL-F}/AhR^{+/+}$. Bars show the normalized enrichment score (NES); pathways with Benjamini-Hochberg adjusted p<0.05 are shown. **(L)** As in (K), for glutamatergic neurons. **(M)** Heatmap of normalized enrichment scores (NES) for selected pathways across sex and neuronal subtype (male GABAergic, male glutamatergic, female GABAergic, female glutamatergic), $APP^{NL-F}/AhR^{-/-}$ vs $APP^{NL-F}/AhR^{+/+}$. WT in the graphs denotes $APP^{NL-F}/AhR^{+/+}$.

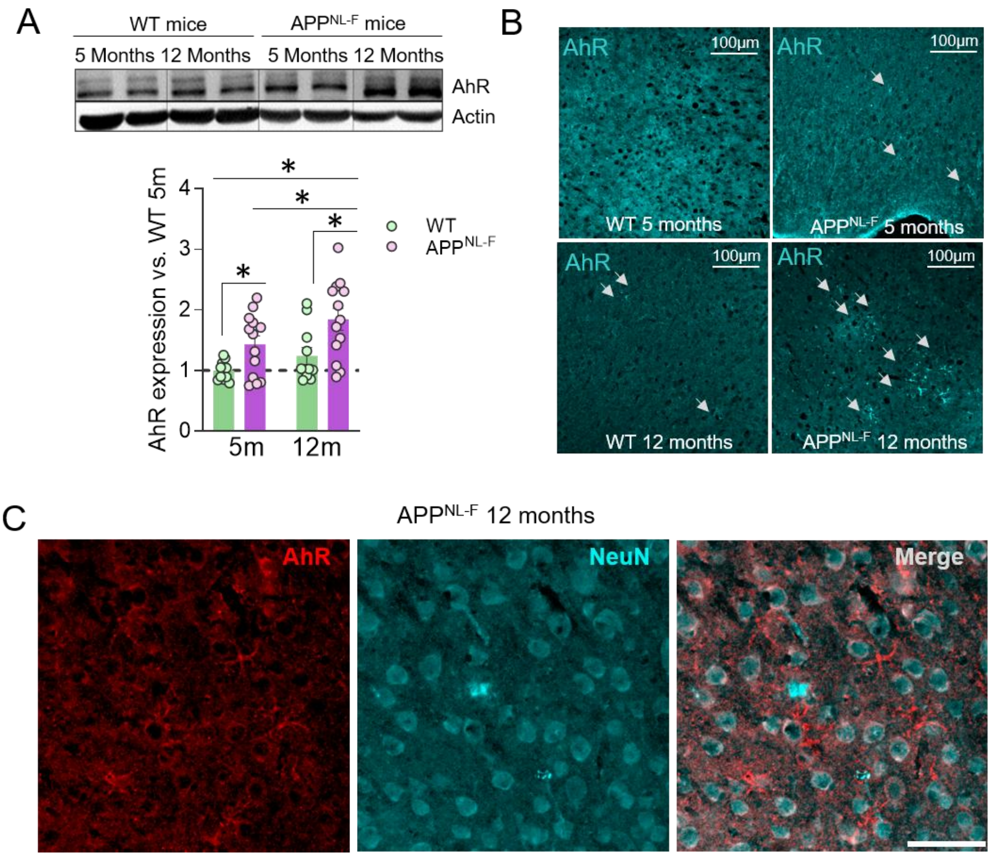


**Supplementary Figure 4. Cortical AhR expression is elevated in APP$^{NL-F}$ mice and increases with disease progression. (A)** Representative western blot and densitometric quantification of AhR protein expression in cortical extracts from WT and APP$^{NL-F}$ mice at 5 and 12 months of age. Actin was used as a loading control. AhR levels were normalized to actin and expressed relative to those measured in 5-month-old WT mice. WT: n=11 per age; APP$^{NL-F}$: n=13 per age. Two-way ANOVA followed by Fisher's LSD multiple-comparisons test. WT versus APP$^{NL-F}$: p=0.0361 at 5 months and p=0.0041 at 12 months. Five versus 12 months: WT, p=0.2510; APP$^{NL-F}$, p=0.0355. **(B)** Representative cortical images showing AhR immunoreactivity in WT and APP$^{NL-F}$ mice at 5 and 12 months of age. Arrowheads indicate cells with prominent AhR immunoreactivity. Scale bars = 100 µm. **(C)** Representative cortical images from 12-month-old APP$^{NL-F}$ mice showing AhR immunoreactivity (red), the neuronal marker NeuN (cyan) and merged channels. AhR immunoreactivity showed limited overlap with NeuN-positive neurons, consistent with its predominant localization in cortical glial cells. Scale bar = 25 µm. Data in (A) are presented as mean±SEM. Each dot represents one biologically independent mouse. Statistical significance was considered at p<0.05.

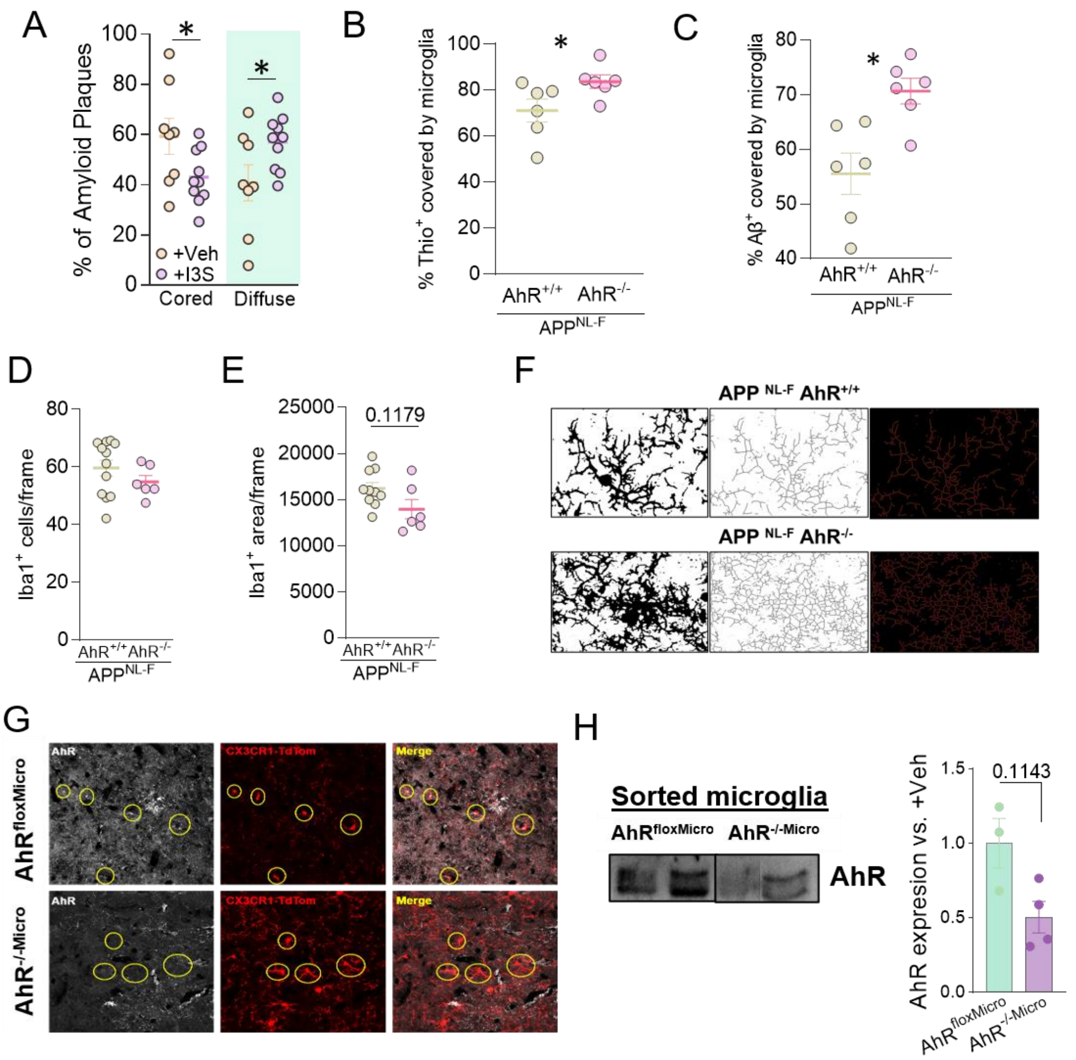


**Supplementary Figure 5. Additional analyses of plaque composition and microglial responses, and characterization of inducible microglial AhR deletion.** **(A)** Proportion of dense-cored and diffuse amyloid plaques in APP$^{NL\text{-}F}$ mice treated with vehicle or indoxyl-3-sulfate (I3S). Dense-cored plaques were defined as Aβ-positive deposits containing a ThioS-positive fibrillar core, whereas diffuse plaques lacked a ThioS-positive core. Vehicle: n=8; I3S: n=10. Two-way ANOVA followed by Fisher's LSD multiple-comparisons test. Vehicle versus I3S: p=0.0372 for dense-cored plaques and p=0.0372 for diffuse plaques. **(B)** Mean percentage of the ThioS-positive plaque area covered by Iba1-positive microglia in APP$^{NL\text{-}F}$/AhR$^{+/+}$ and APP$^{NL\text{-}F}$/AhR$^{-/-}$ mice. Measurements obtained from individual plaques were averaged for each mouse. n=6 mice per group. Two-tailed exact Mann-Whitney U test, p=0.0260. **(C)** Mean percentage of the total Aβ-positive plaque area covered by Iba1-positive microglia in APP$^{NL\text{-}F}$/AhR$^{+/+}$ and APP$^{NL\text{-}F}$/AhR$^{-/-}$ mice. Measurements obtained from individual plaques were averaged for each mouse. n=6 mice per group. Two-tailed exact Mann-Whitney U test, p=0.0087. **(D)** Number of Iba1-positive microglial cells per cortical field in APP$^{NL\text{-}F}$/AhR$^{+/+}$ and APP$^{NL\text{-}F}$/AhR$^{-/-}$ mice. APP$^{NL\text{-}F}$/AhR$^{+/+}$: n=12; APP$^{NL\text{-}F}$/AhR$^{-/-}$: n=6. Two-tailed exact Mann-Whitney U test, p=0.2807. **(E)** Mean Iba1-positive area per cortical field in APP$^{NL\text{-}F}$/AhR$^{+/+}$ and APP$^{NL\text{-}F}$/AhR$^{-/-}$ mice. APP$^{NL\text{-}F}$/AhR$^{+/+}$: n=10; APP$^{NL\text{-}F}$/AhR$^{-/-}$: n=6. Two-tailed exact Mann-Whitney U test, p=0.1179. **(F)** Representative images illustrating the image-processing and skeletonization workflow used to analyse microglial morphology in APP$^{NL\text{-}F}$/AhR$^{+/+}$ and APP$^{NL\text{-}F}$/AhR$^{-/-}$ mice. Iba1-positive microglia were segmented and skeletonized using Fiji to quantify morphological parameters. Enlarged images illustrate the identification of individual branches and endpoints within the microglial skeleton. **(G)** Representative cortical images showing AhR immunoreactivity and CX3CR1-driven tdTomato labeling of microglia in vehicle-treated control APP$^{NL-F}$/AhR$^{fl/fl}$ mice and tamoxifen-treated APP$^{NL-F}$/AhR$^{KO-Mic}$ mice. Circles indicate tdTomato-positive microglia used to assess the reduction in microglial AhR immunoreactivity following tamoxifen-induced recombination. Scale bars as indicated. **(H)** Representative western blot and densitometric analysis of AhR in microglia isolated by fluorescence-activated cell sorting from 6-month-old vehicle-treated control APP$^{NL-F}$/AhR$^{fl/fl}$ mice and tamoxifen-treated APP$^{NL-F}$/AhR$^{KO-Mic}$ mice. AhR band intensity was expressed relative to the mean value of the vehicle-treated control group. Vehicle: n=3; tamoxifen: n=4. Two-tailed exact Mann-Whitney U test, p=0.1143. Data are presented as mean±SEM. In **(A–E)** and **(H)**, each dot represents one biologically independent mouse. Sex distribution in (H) was as follows: APP$^{NL-F}$/AhR$^{fl/fl}$, 2 males/1 female; APP$^{NL-F}$/AhR$^{KO-Mic}$, 2 males/2 females. Statistical significance was considered at p<0.05.

A

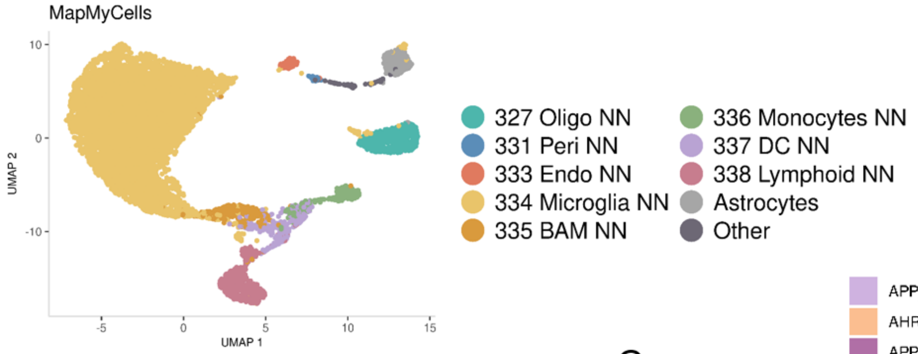


B

| | APP_Male | AHR_KO_Male | APP_Female | AHR_KO_Female |
|---|---|---|---|---|
| 327 Oligo NN | 6.66 | 8.52 | 2.95 | 4.86 |
| 331 Peri NN | 0.41 | 0.17 | 0.38 | 0.33 |
| 333 Endo NN | 0.27 | 1.09 | 0.94 | 1.72 |
| 334 Microglia NN | 73.42 | 65.29 | 80.4 | 77.27 |
| 335 BAM NN | 4.01 | 2.23 | 3.55 | 2.96 |
| 336 Monocytes NN | 1.77 | 5.66 | 1.13 | 2.19 |
| 337 DC NN | 2.58 | 5.49 | 2.42 | 2.34 |
| 338 Lymphoid NN | 6.19 | 7.26 | 5.59 | 4.75 |
| Astrocytes | 3.74 | 3.32 | 1.66 | 2.67 |
| Other | 0.95 | 0.97 | 0.98 | 0.91 |

C

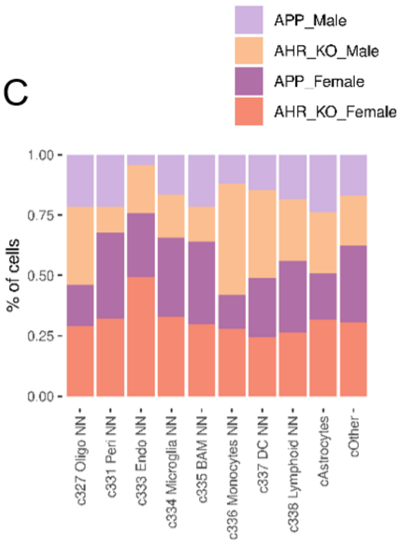


**Supplementary Figure 6. Cell-type annotation and group distribution of cortical non-neuronal cells analysed by single-cell RNA sequencing.** Cortical non-neuronal cells were enriched using a Percoll density gradient and analysed by single-cell RNA sequencing. **(A)** UMAP representation of the complete dataset showing cell-type annotations obtained by hierarchical mapping with MapMyCells against the Whole Mouse Brain reference taxonomy. Cells were assigned to oligodendrocytes (327 Oligo NN), pericytes (331 Peri NN), endothelial cells (333 Endo NN), microglia (334 Microglia NN), border-associated macrophages (335 BAM NN), monocytes (336 Monocytes NN), dendritic cells (337 DC NN), lymphoid cells (338 Lymphoid NN), astrocytes or other cell types. Cell-class labels are reported using the original MapMyCells nomenclature. **(B)** Percentage of each annotated cell population within male APP$^{NL-F}$/AhR$^{+/+}$, male APP$^{NL-F}$/AhR$^{-/-}$, female APP$^{NL-F}$/AhR$^{+/+}$ and female APP$^{NL-F}$/AhR$^{-/-}$ mice. Microglia constituted the predominant cell population in all four groups. **(C)** Relative contribution of the four sex–genotype groups to each annotated cell population. For each cell type, the contributions of male APP$^{NL-F}$/AhR$^{+/+}$, male APP$^{NL-F}$/AhR$^{-/-}$, female APP$^{NL-F}$/AhR$^{+/+}$and female APP$^{NL-F}$/AhR$^{-/-}$cells sum to 100%. No inferential statistical comparisons were performed.

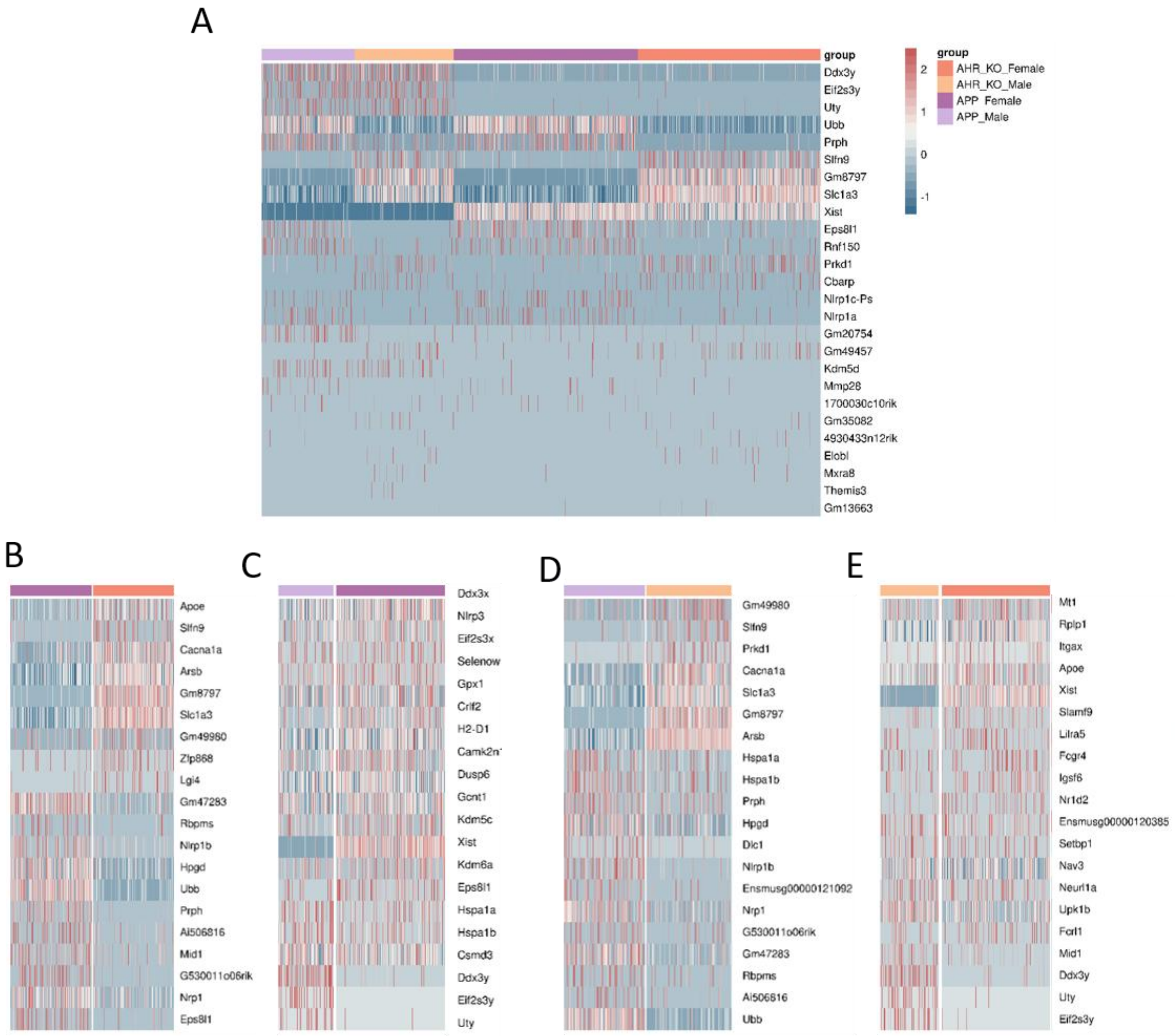


**Supplementary Figure 7. Single-cell expression patterns of sex- and AhR-dependent differentially expressed genes in APP$^{NL-F}$ microglia.** Microglial cells identified by MapMyCells annotation were subsetted and reanalysed using Seurat. Differential gene-expression analysis was performed using Wilcoxon rank-sum tests. **(A)** Heatmap showing the expression of selected differentially expressed genes across microglial cells from male APP$^{NL-F}$/AhR$^{+/+}$, male APP$^{NL-F}$/AhR$^{-/-}$, female APP$^{NL-F}$/AhR$^{+/+}$ and female APP$^{NL-F}$/AhR$^{-/-}$mice. **(B)** Expression of selected differentially expressed genes between female APP$^{NL-F}$/AhR$^{+/+}$ and female APP$^{NL-F}$/AhR$^{-/-}$ microglia. **(C)** Expression of selected sex-dependent genes between male and female APP$^{NL-F}$/AhR$^{+/+}$microglia. **(D)** Expression of selected differentially expressed genes between male APP$^{NL-F}$/AhR$^{+/+}$ and male APP$^{NL-F}$/AhR$^{-/-}$ microglia. **(E)** Expression of selected sex-dependent genes between male and female APP$^{NL-F}$/AhR$^{-/-}$ microglia. Rows represent genes and columns represent individual microglial cells. Colours indicate row-scaled gene-expression values, with red indicating higher and blue indicating lower relative expression. Genes with Benjamini-Hochberg-adjusted p≤0.05 were considered differentially expressed.

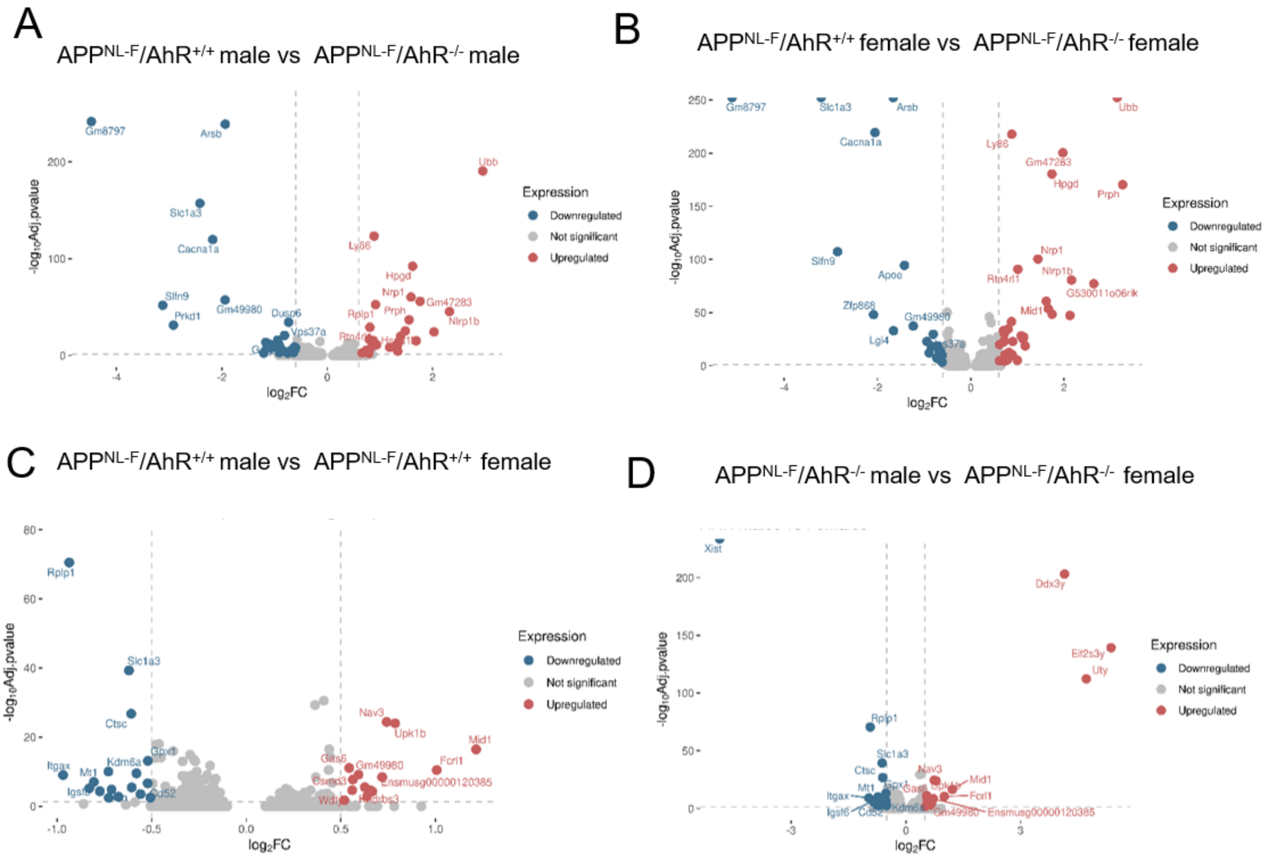


**Supplementary Figure 8. Differential gene-expression signatures associated with AhR loss and biological sex in APP$^{NL-F}$ microglia.** Volcano plots showing pairwise differential gene-expression analyses of individual microglial cells using Wilcoxon rank-sum tests. **(A)** Differentially expressed genes between male APP$^{NL-F}$/AhR$^{+/+}$ and female APP$^{NL-F}$/AhR$^{-/-}$ microglia. **(B)** Differentially expressed genes between female APP$^{NL-F}$/AhR$^{+/+}$ and female APP$^{NL-F}$/AhR$^{-/-}$ microglia. **(C)** Sex-dependent differentially expressed genes between male and female APP$^{NL-F}$/AhR$^{+/+}$ microglia, excluding sex-linked genes. **(D)** Sex-dependent differentially expressed genes between male and female APP$^{NL-F}$/AhR$^{-/-}$ microglia. The x axis represents log2 fold change, and the y axis represents −log10 of the Benjamini-Hochberg-adjusted p value. Positive log2 fold-change values indicate higher expression in the first group named in each comparison, whereas negative values indicate higher expression in the second group. Significantly upregulated and downregulated genes are shown in red and blue, respectively; non-significant genes are shown in grey. Dashed lines indicate the significance and fold-change thresholds used for differential gene-expression analysis. Selected genes are labelled. Genes with Benjamini-Hochberg-adjusted $p \leq 0.05$ were considered differentially expressed.

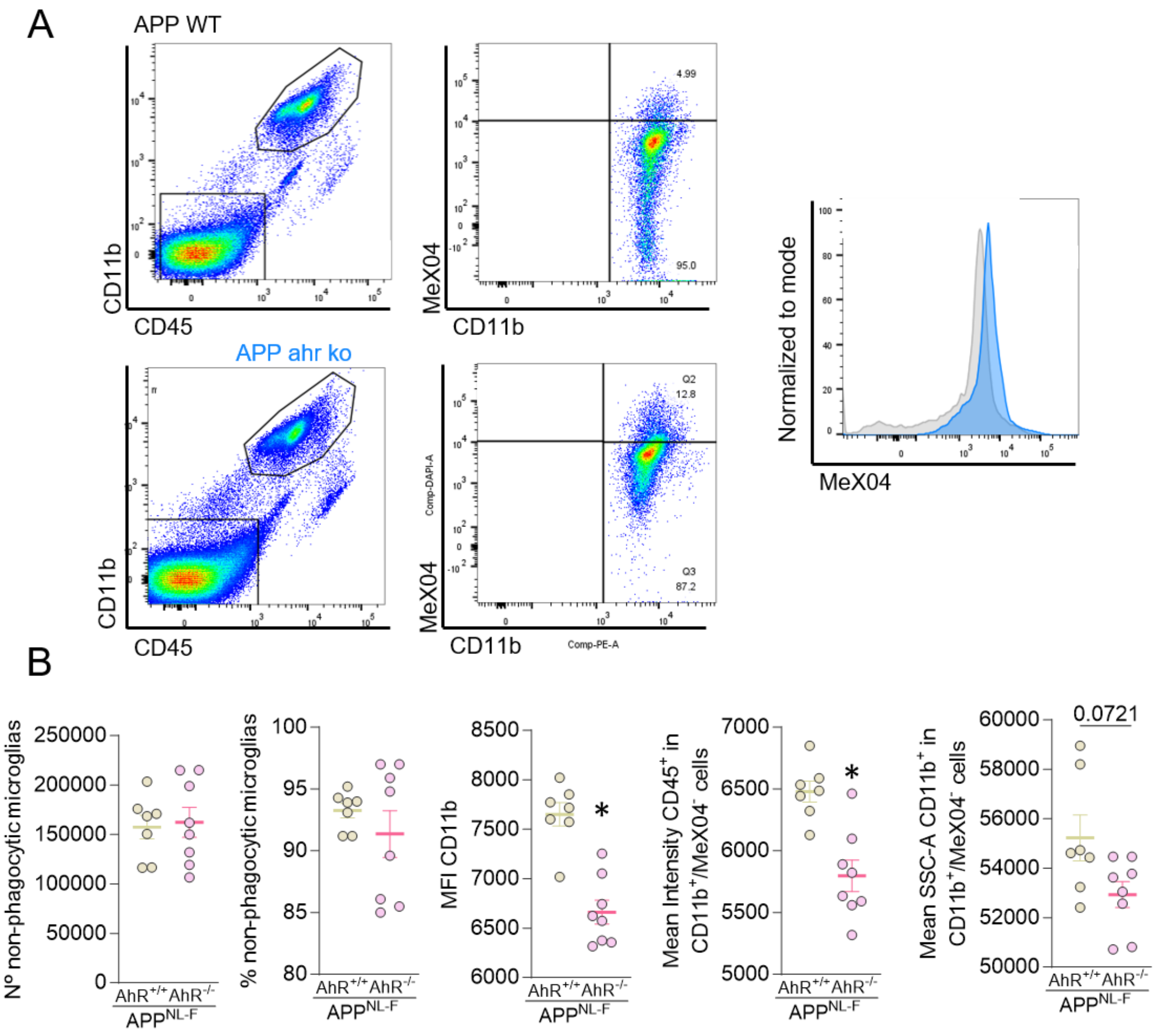


**Supplementary Figure 9. Flow-cytometric gating strategy and phenotypic characterization of Methoxy-X04-negative microglia in $APP^{NL-F}$ mice. (A)** Representative flow-cytometry plots showing the gating strategy used to identify microglia and quantify cellular uptake of Methoxy-X04-labeled fibrillar Aβ in $APP^{NL-F}$/$AhR^{+/+}$ and female $APP^{NL-F}$/$AhR^{-/-}$ mice. Microglia were identified according to their CD11b and CD45 expression, followed by separation into Methoxy-X04-positive and Methoxy-X04-negative populations. Representative histograms show the distribution of Methoxy-X04 fluorescence intensity in the indicated genotypes. **(B)** Absolute number and percentage of Methoxy-X04-negative microglia, median fluorescence intensity of CD11b, mean CD45 fluorescence intensity and mean side-scatter area (SSC-A) within the CD11b-positive/CD45-positive/Methoxy-X04-negative population from $APP^{NL-F}$/$AhR^{+/+}$ and female $APP^{NL-F}$/$AhR^{-/-}$ mice. Data are presented as mean±SEM. Each dot represents one biologically independent mouse. Statistical significance was considered at $p<0.05$.

A

% of cells per cluster

| | 0 | 1 | 2 | 3 | 4 | 5 | 6 | 7 |
|---|---|---|---|---|---|---|---|---|
| APP_Male | 61.85 | 1.76 | 13.06 | 11.94 | 7.96 | 2.04 | 0.74 | 0.65 |
| AHR_KO_Male | 4.38 | 53.33 | 20.05 | 10.51 | 7.01 | 3.15 | 1.05 | 0.53 |
| APP_Female | 62.33 | 1.36 | 14.04 | 8.6 | 8.64 | 3.52 | 0.85 | 0.66 |
| AHR_KO_Female | 1.99 | 61.97 | 14.24 | 7 | 10.03 | 3.41 | 0.9 | 0.47 |

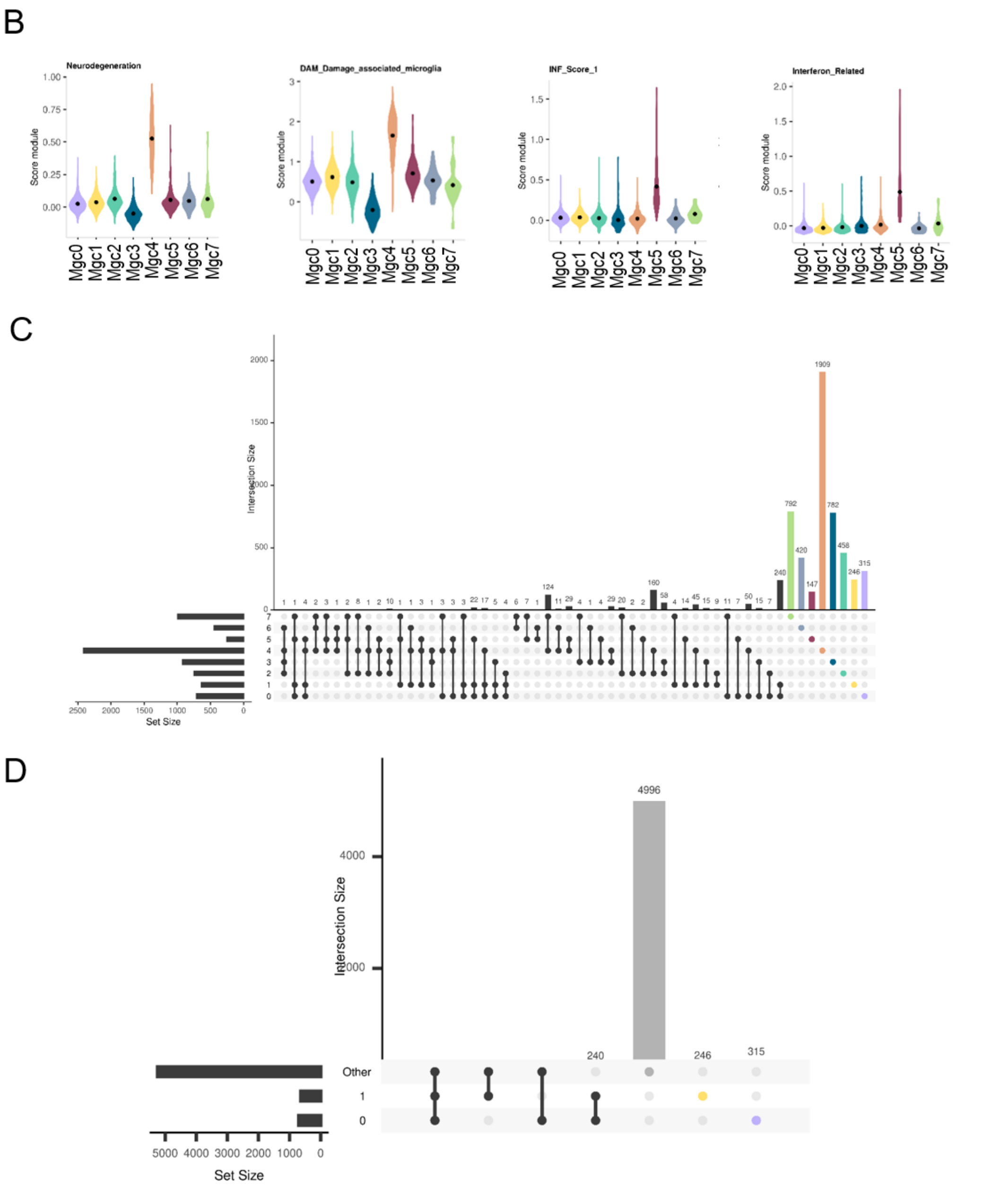


**Supplementary Figure 10. Relative abundance, disease-associated transcriptional programs and differential gene overlap across microglial subclusters. (A)** Percentage of microglial cells assigned to each subcluster, MgC0-MgC7, in male APP$^{NL-F}$/AhR$^{+/+}$, male APP$^{NL-F}$/AhR$^{-/-}$, female APP$^{NL-F}$/AhR$^{+/+}$ and female APP$^{NL-F}$/AhR$^{-/-}$ mice. **(B)** Distribution of neurodegeneration-associated, disease-associated microglia (DAM) and interferon-response module scores across MgC0–MgC7. Module scores were calculated at the single-cell level using curated gene signatures derived from previously published studies. Violin plots show the distribution of module scores across individual cells within each microglial subcluster. The genes included in each transcriptional module and their corresponding references are provided in

Supplementary Table 3. **(C)** UpSet plot showing the number and overlap of differentially expressed genes identified across MgC0-MgC7. Horizontal bars indicate the total number of differentially expressed genes associated with each subcluster, whereas vertical bars indicate the size of the gene intersections represented by the connected-dot combinations below. **(D)** Simplified UpSet plot showing the overlap of differentially expressed genes associated with MgC0, MgC1 and the remaining microglial subclusters. “Other” includes genes associated with any microglial subcluster other than MgC0 or MgC1, encompassing MgC2-MgC7. Horizontal bars indicate the total size of each gene set, and vertical bars indicate the number of genes in each intersection.

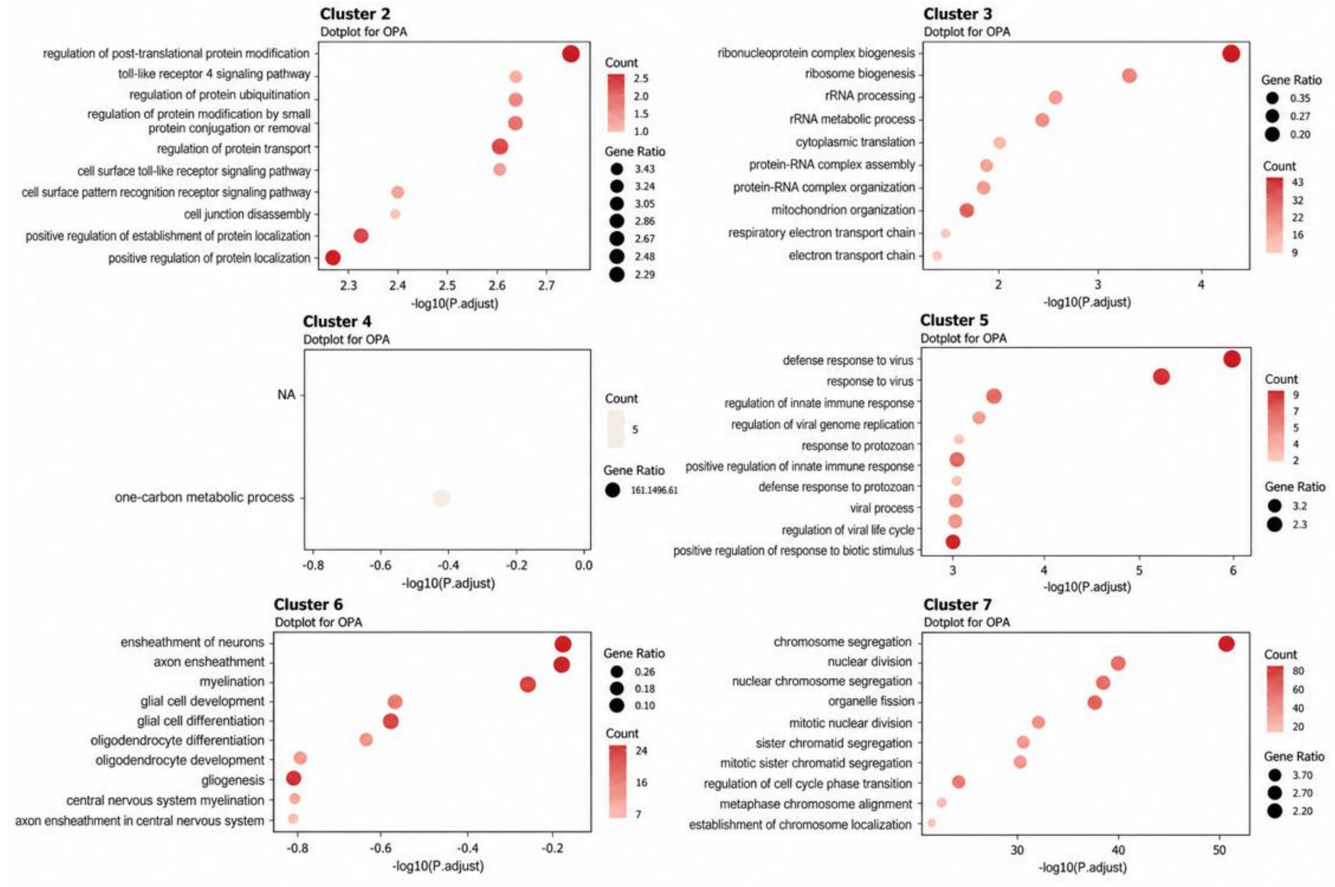


**Supplementary Figure 11. Gene Ontology enrichment analysis reveals distinct biological programs across microglial subclusters.** Gene Ontology (GO) Biological Process over-representation analysis (ORA) of cluster-associated genes in microglial subclusters MgC2-MgC7 from 12-month-old APPNL-F mice. Dot plots show enriched biological processes for MgC2, MgC3, MgC4, MgC5, MgC6 and MgC7, arranged from left to right and top to bottom. Enrichment analysis included genes significantly upregulated in each subcluster relative to the remaining microglial cells (adjusted p ≤ 0.05). Dot size represents the gene ratio, defined as the proportion of input genes associated with each GO term, whereas dot colour indicates the number of associated genes (Count). The x-axis represents $-\log_{10}$ of the adjusted p value. Enrichment was assessed using a hypergeometric test followed by Benjamini-Hochberg correction for multiple comparisons. GO terms with adjusted p ≤ 0.05 were considered significantly enriched.

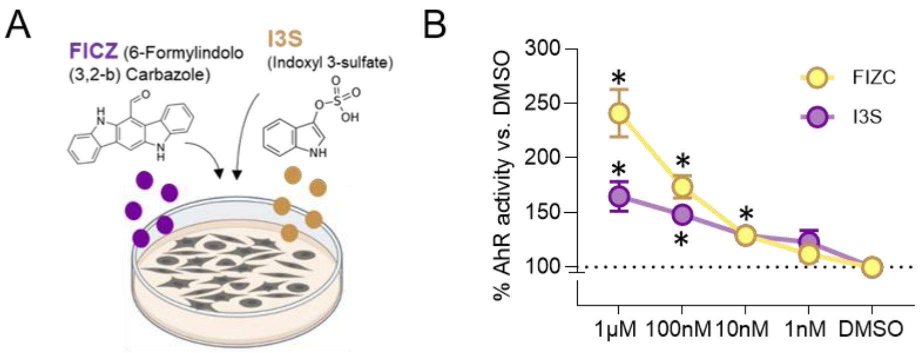


**Supplementary Figure 12. Validation of the AhR-responsive luciferase reporter assay using FICZ and indoxyl-3-sulfate. (A)** Schematic representation of the XRE-driven luciferase reporter assay used to evaluate AhR activation by the reference agonist 6-formylindolo[3,2-b]carbazole (FICZ) and indoxyl-3-sulfate (I3S). **(B)** Concentration-dependent activation of the AhR reporter following treatment with FICZ or I3S at concentrations ranging from 1 nM to 1 µM. Firefly luciferase activity was normalized to Renilla luciferase activity and expressed relative to the DMSO-treated condition, set to 100%. Kruskal–Wallis test followed by Dunn's multiple-comparisons test against DMSO. Data are presented as mean±SEM. Asterisks indicate statistically significant differences compared with DMSO, p<0.05.

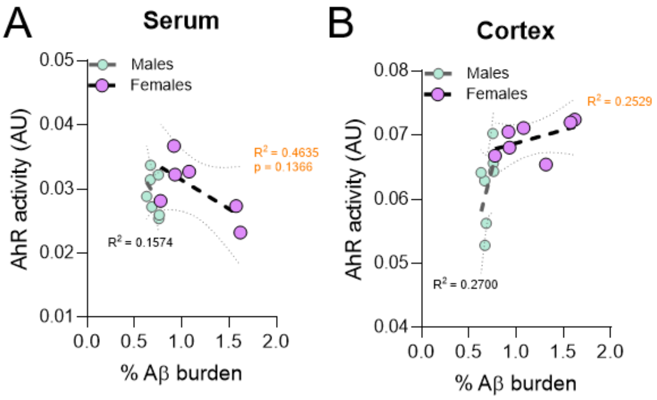


**Supplementary Figure 13. Associations between AhR-activating bioactivity and cortical amyloid burden in APP$^{NL-F}$ mice. (A-B)** Associations between cortical Aβ burden and AhR-activating bioactivity measured using the XRE-driven luciferase reporter assay in serum **(A)** and cortical extracts **(B)** from 12-month-old male and female APP$^{NL-F}$ mice. Female: n=7; male: n=7. Associations were analysed separately by sex using Spearman rank correlation. Dashed lines represent linear regression fits, and dotted lines indicate the corresponding 95% confidence intervals. $R^2$ values indicate the goodness of fit of the linear regression. Each dot represents one biologically independent mouse.

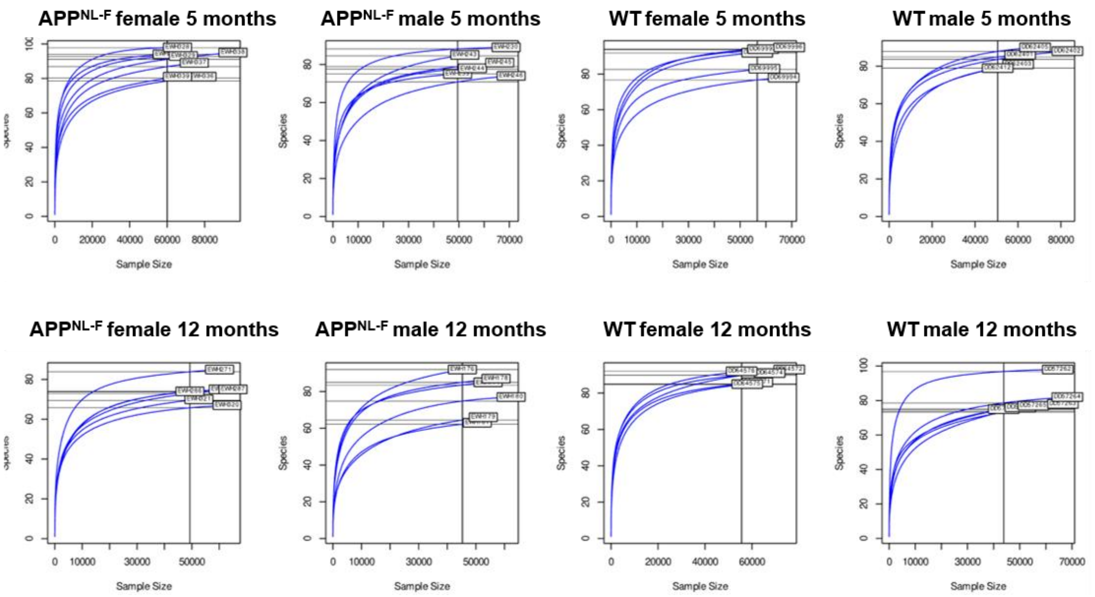


**Supplementary Figure 14. Rarefaction analysis of fecal microbiota sequencing depth in WT and APP$^{NL-F}$ mice. (A-H)** Rarefaction curves showing the number of observed bacterial species as a function of sequencing depth in faecal samples from 5-month-old female APP$^{NL-F}$ mice **(A)**, 5-month-old male APP$^{NL-F}$ mice **(B)**, 5-month-old female WT mice **(C)**, 5-month-old male WT mice **(D)**, 12-month-old female APP$^{NL-F}$ mice **(E)**, 12-month-old male APP$^{NL-F}$ mice **(F)**, 12-month-old female WT mice **(G)** and 12-month-old male WT mice **(H)**. Each curve represents one biologically independent mouse. The tendency of the curves to approach a plateau indicates that sequencing depth was sufficient to capture most of the bacterial richness present in the analysed samples.

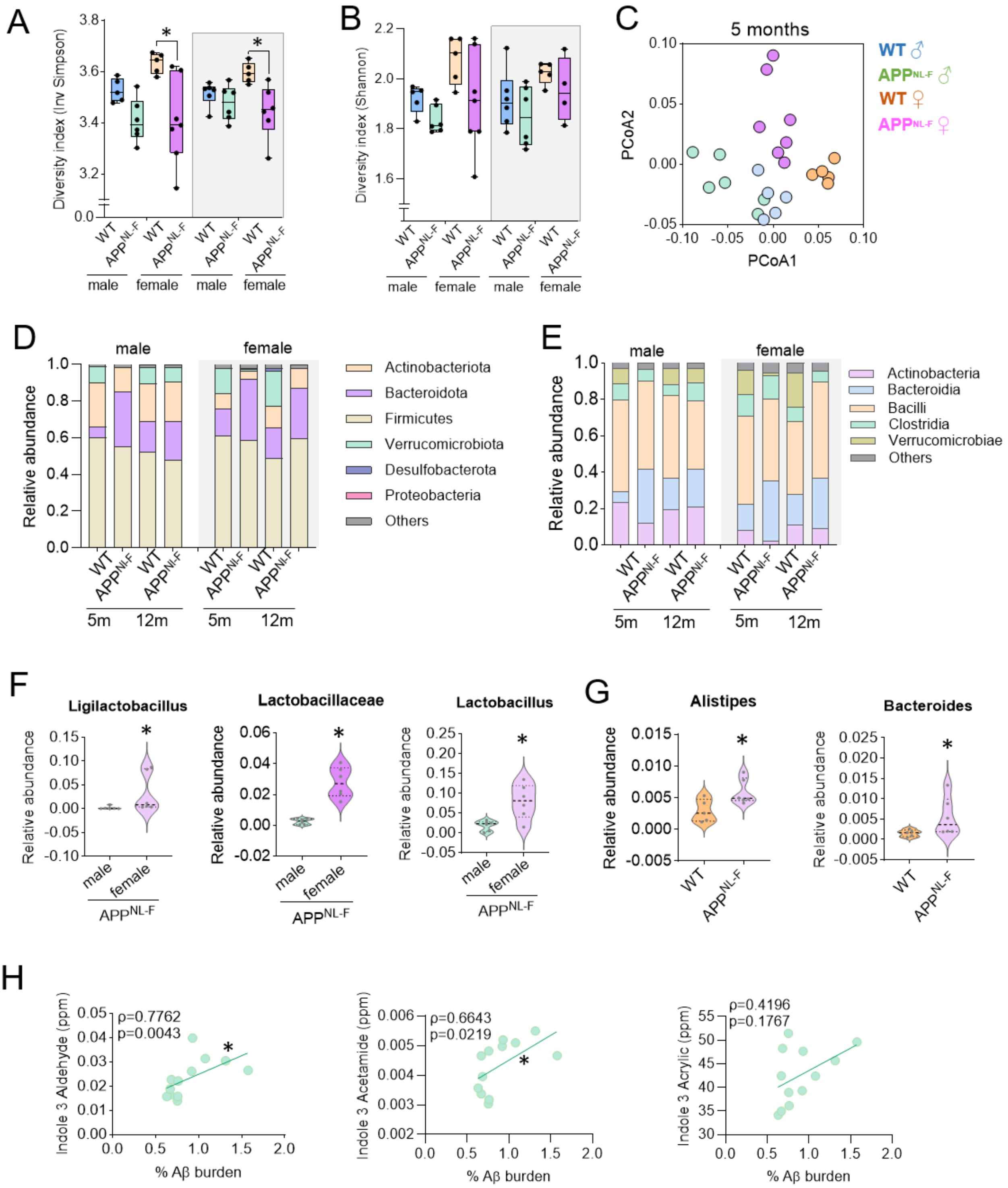


**Supplementary Figure 15. Additional sex- and genotype-dependent alterations in gut microbial diversity, taxonomic composition and AhR-related indole metabolites. (A-B)** Inverse Simpson **(A)** and Shannon **(B)** diversity indices of the faecal microbiota from male and female WT and $APP^{NL\text{-}F}$ mice at 5 and 12 months of age. At 5 months, WT males: n=5; WT females: n=5; $APP^{NL\text{-}F}$ males: n=6; $APP^{NL\text{-}F}$ females: n=7. At 12 months, WT males: n=6; WT females: n=5; $APP^{NL\text{-}F}$ males: n=6; $APP^{NL\text{-}F}$ females: n=6. Two-way ANOVA followed by multiple-comparisons tests. **(C)** Principal coordinates analysis based on Bray–Curtis dissimilarities in faecal samples from 5-month-old male and female WT and $APP^{NL\text{-}F}$ mice. Each point represents one biologically independent mouse. Overall PERMANOVA: $R^2$=0.5520, pseudo-F=7.8024, p=0.0010. Pairwise PERMANOVA with false-discovery-rate correction: $APP^{NL\text{-}F}$ versus WT females, q=0.0100; $APP^{NL\text{-}F}$ versus WT males, q=0.0120; female versus male $APP^{NL\text{-}F}$ mice, q=0.0080; and female versus male WT mice, q=0.0107. **(D-E)** Relative abundance of the predominant bacterial phyla **(D)** and classes **(E)** in the faecal microbiota of male and female WT and $APP^{NL\text{-}F}$ mice at 5 and 12 months of age. Group sizes were as indicated in **(A-B)**. **(F)** Relative abundance of Ligilactobacillus, Lactobacillaceae and Lactobacillus in male and female $APP^{NL\text{-}F}$ mice at 12 months of age. **(G)** Relative abundance of Alistipes and Bacteroides in 12-month-old female WT and $APP^{NL\text{-}F}$ mice. Comparisons were performed using two-tailed Mann–Whitney U tests. *p < 0.05. **(H)** Associations of serum indole-3-aldehyde, indole-3-acetamide and indole-3-acrylic acid concentrations with cortical Aβ plaque number and cortical Aβ burden in 12-month-old $APP^{NL\text{-}F}$ mice. n=12 mice per analysis. Spearman correlation: indole-3-aldehyde versus Aβ burden, ρ=0.7762, p=0.0042; indole-3-acetamide versus Aβ burden, ρ=0.6643, p=0.0219; indole-3-acrylic acid versus Aβ burden, ρ=0.4196, p=0.1767; and indole-3-acrylic acid versus plaque number, ρ=0.2762, p>0.05. Solid lines represent linear regression fits. Data are presented as mean±SEM where applicable. Each dot represents one biologically independent mouse. Asterisks indicate *p<0.05.

### Supplementary Tables

**Supplementary Table 1. Analytical parameters used for the targeted semi-quantification of selected metabolites by UHPLC–QqQ-MS/MS and CE–MS.** Retention times are reported for compounds analysed by UHPLC–QqQ-MS/MS, whereas migration times are reported for compounds analyzed by CE–MS. Selected MRM transitions or quantification ions, fragmentor voltages, and analyte-specific calibration ranges are shown. Eight-point external calibration curves were constructed for each compound using concentration ranges selected according to the corresponding linear response.

| Compound Name | Molecular Formula | CAS No. | RT/MT (min) | Transition / ion (m/z) | Fragmentor (V) | Calibration range (mg/L) |
|---|---|---|---|---|---|---|
| Indole-3-aldehyde (IAld; indole-3-carboxaldehyde) | $C_9H_7NO$ | 487-89-8 | 1.34 | 146.1→118.1 | 166 | 2.5–500 |
| Indole-3-acetamide (I3AM) | $C_{10}H_{10}N_2O$ | 879-37-8 | 1.16 | 175.0→130.1 | 166 | 4.1–835 |
| trans-Indole-3-acrylic acid (I3Acr) | $C_{11}H_9NO_2$ | 29953-71-7 | 5.38 | 188.1→146.1 | 166 | 2.5–50 |
| n-Butyric acid | $C_4H_8O_2$ | 107-92-6 | 7.5 | 106.0→60.3 | 166 | 2.5–50 |
| L-tryptophan | $C_{11}H_{12}N_2O_2$ | 73-22-3 | 12.4 | 204.0894 | - | 2.5–62.5 |
| Choline chloride | $C_5H_{14}ClNO$ | 67-48-1 | 8.74 | 104.107 | - | 0.5–20 |

**Supplementary Table 2. Neuronal single-nucleus RNA-seq differential expression and pathway enrichment analyses in $APP^{NL\text{-}F}$ mice following AhR deletion.** Detailed differential gene-expression and gene-set enrichment analyses of cortical neuronal nuclei from 12-month-old WT, $APP^{NL\text{-}F}$ and $APP^{NL\text{-}F}/AhR^{-/-}$ mice. Analyses focus on genotype-associated differences between $APP^{NL\text{-}F}/AhR^{-/-}$ and $APP^{NL\text{-}F}/AhR^{+/+}$ neurons, including combined neuronal populations, GABAergic and glutamatergic neurons, and sex-stratified comparisons. The workbook contains the results underlying Figure 2J and Supplementary Figure 3H–M, including selected differentially expressed genes, neuronal and neurodegeneration-related pathways, and sex-specific enrichment profiles. Differential-expression results include log2 fold changes, the proportion of nuclei expressing each gene, and unadjusted and adjusted p values. Gene-set enrichment results include enrichment scores (ES), normalized enrichment scores (NES), gene-set sizes and Benjamini–Hochberg-adjusted p values. Each sex–genotype combination was represented by one independently processed pooled sample comprising cortical tissue from two mice.

**Supplementary Table 3. Single-cell transcriptomic characterization of microglial responses to AhR deficiency in $APP^{NL\text{-}F}$ mice.** Detailed single-cell RNA-seq analyses of cortical non-neuronal cells from 12-month-old male and female $APP^{NL\text{-}F}$/$AhR^{+/+}$ and $APP^{NL\text{-}F}$/$AhR^{-/-}$ mice, with two biological replicates per sex–genotype group. The workbook includes cell-type counts and proportions underlying Figure 4C; differential gene-expression analyses comparing sex and genotype groups; Ingenuity Pathway Analysis (IPA) of canonical pathways; microglial subcluster distributions underlying Figure 5D; and marker-gene identification and Gene Ontology (GO) enrichment analyses for microglial subclusters MgC0–MgC7. Differential-expression results include log2 fold changes, the proportion of cells expressing each gene, and unadjusted and adjusted p values. IPA results include pathway enrichment significance, enrichment ratios, predicted activation z-scores and contributing genes, whereas GO analyses report enriched biological processes and associated statistics. The workbook also provides the curated gene signatures used for disease-associated microglia, neurodegeneration-related and interferon-response module scoring.

**Supplementary Table 4. Demographic, clinical and biomarker characteristics of COGNIBIO12 participants according to AT biomarker profile.** COGNIBIO12 participants with available CSF Aβ42 and p-tau181 measurements (n=93) were classified into A−T− (n=11), A+T− (n=10), A−T+ (n=15) and A+T+ (n=57) groups. Amyloid positivity (A+) was defined as CSF Aβ42 ≤599 pg/mL and tau positivity (T+) as CSF p-tau181 ≥56.5 pg/mL, using laboratory-established cutoffs. CSF Aβ40 and the Aβ42/Aβ40 ratio were available in a subset of participants (n=69) and were included as descriptive variables but were not used to define the primary AT groups. Data are presented as mean ± SD for age, median [interquartile range (IQR)] for continuous biomarker variables, and n/available n (%) for categorical variables. Separate n values indicate the number of participants with available measurements for each variable. Serum and CSF AhR-activating bioactivity was determined using an XRE-driven luciferase reporter assay and is expressed as normalized relative activity. CSF, cerebrospinal fluid; AhR, aryl hydrocarbon receptor; MMSE, Mini-Mental State Examination; GDS, Global Deterioration Scale; IQR, interquartile range; SD, standard deviation.

**Supplementary Table 5. Statistical analyses of AhR-activating bioactivity in the human cohorts.** Summary of statistical analyses examining serum and CSF AhR-activating bioactivity in independently recruited H2H cognitively unimpaired participants and the COGNIBIO12 analytical subcohort. AhR bioactivity was natural-log transformed for parametric analyses. Group comparisons were performed using linear regression models adjusted for age and sex with heteroscedasticity-consistent HC3 standard errors. Exponentiated regression coefficients are reported as adjusted geometric mean ratios with 95% confidence intervals (CI). Sex-specific group comparisons were derived from a common group-by-sex interaction model adjusted for age. For analyses across AT biomarker profiles, the global group effect was assessed first, followed by planned comparisons against the A−T− reference group with Holm correction. Associations between continuous variables were assessed using Pearson or Spearman correlations, as indicated, and age- and sex-adjusted regression models. Adjusted estimates for continuous predictors are expressed per 1-SD increase in the predictor. Benjamini–Hochberg false-discovery-rate (FDR) q values are reported where applicable. Additional sensitivity analyses assessed the influence of CSF Aβ40 availability and alternative amyloid classification using the CSF Aβ42/Aβ40 ratio (<0.069). Primary amyloid positivity was defined as CSF Aβ42 ≤599 pg/mL and tau positivity as CSF p-tau181 ≥56.5 pg/mL.

**Supplementary Table 6. Predicted microbial functional alterations associated with sex and $APP^{NL\text{-}F}$ genotype.** PICRUSt2-based analysis of predicted microbial functions showing significant differences between $APP^{NL\text{-}F}$ females and $^{APPNL\text{-}F}$ males, and between $APP^{NL\text{-}F}$ females and WT females. Significantly altered functions (FDR-adjusted $q < 0.05$) are reported together with the direction and magnitude of the change. Particular attention is drawn to functions related to aromatic compound and aldehyde metabolism, including alcohol dehydrogenase, aryl-alcohol dehydrogenase, acetaldehyde dehydrogenase, and 2-hydroxymuconate tautomerase, which were increased in $APP^{NL\text{-}F}$ females and may contribute to the sex-dependent metabolic profile associated with amyloid pathology.
